# Sex-divergent brain epigenetic reprogramming by chronic opioids

**DOI:** 10.1101/2025.10.27.684830

**Authors:** Camille Falconnier, Alba Caparros-Roissard, Hanus Slavik, Mithil Gaikwad, Victor Mathis, Margot Diringer, Charles Decraene, Salim Megat, Sebahat Ozkan, Mohamad Yassine, Yahia Hadj-Arab, Pierre Hener, Robin Waegaert, Brigitte L. Kieffer, Emmanuel Darcq, Laurence Lalanne, Karine Merienne, Anne-Laurence Boutillier, Anaïs Flore Bardet, Ipek Yalcin, Pierre-Eric Lutz

## Abstract

Opioid use disorder (OUD) is a chronic condition that exhibits sex differences in prevalence, symptoms and treatment. Yet, the epigenetic mechanisms underlying these differences remain largely unknown. Here, we investigated the nucleus accumbens, a key brain region in OUD, to define the multiomic consequences of chronic morphine exposure in male and female mice. We profiled DNA methylation, five histone post-translational modifications, and their transcriptional effects at bulk and cell-type-specific levels. Despite comparable tissue organization and neurophysiological responses to morphine, epigenetic adaptations occurred at highly sex-specific genomic loci. These adaptations nevertheless followed common mechanistic principles, acting at similar gene features and transcription factor binding sites across sexes. Strikingly, they converged on overlapping genes, biological functions, and co-expression modules, and partially recapitulated transcriptional signatures of OUD in men and women. Therefore, our findings uncover a profound epigenetic sex divergence that mediates convergent biological dysregulation, and highlight opportunities for developing improved therapeutic strategies tailored to sex-specific mechanisms.

## Introduction

Opioids are the reference treatment for severe pain. Their chronic consumption, however, is associated with serious adverse outcomes that include development of an Opioid Use Disorder (OUD) and overdose. Worldwide, opioids are responsible for an estimated 450,000 deaths annually^1^, underscoring the urgent need to better understand their biological effects.

OUD is a chronic relapsing condition characterized by tolerance, physical dependence, and invasive behaviors oriented towards consuming opioids, despite harmful consequences. In affected individuals, the risk of relapse persists for decades^2^, suggesting long-lasting brain adaptations that extend beyond the drug’s pharmacological action. A main hypothesis to explain such a protracted clinical course implicates epigenetic processes. While primarily investigated in the context of development and cancer^3,4^, these processes are increasingly recognized as contributing to psychiatric disease^5^. Here, we investigated the epigenomic consequences of chronic opioid exposure in the mouse brain. Unlike previous studies of candidate genes or single epigenetic layers (see^6^ for review), we adopted a genome-wide, multiomic strategy encompassing DNA methylation^7,8^, 5 types of histone post-translational modifications and their transcriptional impact, at bulk and cell-type specific levels.

A second major focus of this study relates to sex differences, which have been described in relation to socio-demographic and epidemiological characteristics of OUD. These differences manifest in terms of risk factors (psychiatric comorbidities in women, legal issues in men^9–11^), prevalence (higher in men^12,13^), or the types of opioids used (heroin in men, prescription drugs in women^10,11,14^). Clinically, women tend to report more pleasurable responses to drugs of abuse, whereas men exhibit more severe withdrawal symptoms but sustain longer periods of abstinence^15^. While these differences partly reflect economic, social, and cultural factors^16^, accumulating evidence from animal models suggests underlying biological mechanisms. Accordingly, female mice show greater sensitivity to the rewarding^17–19^ and motivational effects^20^ of opioids, whereas males display stronger signs of physical dependence^21,22^. Despite this, the epigenetic bases of these sex differences remain unexplored^6^.

To address these 2 objectives, we conducted parallel investigations in female and male mice, focusing on the nucleus accumbens (NAc), a major brain region implicated in OUD^6^. First, we profiled DNA methylation and histone modifications following chronic morphine exposure, using Enzymatic Methylation (EM-seq) and Cleavage Under Targets and Tagmentation (CUT&Tag) sequencing, respectively. Strong differences were observed in epigenetic effects of morphine across females and males, which nonetheless converged on shared genes and biological pathways. Second, using multiplex in situ hybridization and fiber photometry, we evaluated NAc histological organization and neuronal activity in response to morphine across sexes. No significant differences were identified, indicating that epigenomic sex differences did not stem from anatomical or neurophysiological factors. Third, to assess the functional impact of this epigenetic plasticity, we characterized morphine-induced transcriptional adaptations at both bulk and cell type-specific levels. We focused in particular on neurons expressing the mu opioid receptor (MOR), the direct pharmacological target of morphine^23^. These analyses revealed sex-specific transcriptional responses that significantly overlapped with those recently described in the human brain, in men and women with OUD^24^. Finally, through gene co-expression network analysis, we integrated multiomic data to identify the gene modules and biological pathways shaped by morphine in a sex-specific manner, and identified GABAergic interneurons as a primary target for epigenetic reprogramming. Overall, this work unravels genomic loci and cell types for developing sex-specific interventions based on epigenetic editing.

## Results

### Chronic morphine induces widespread but divergent DNA methylation changes in male & female mice

We first identified the methylomic changes induced by chronic morphine using EM-seq (Fig.1a,S1), which provides better coverage of GC-rich regions than bisulfite conversion^25^. Data were generated at 10X coverage/sample in saline-and morphine-treated mice (n=5-6/sex/group, 23 total), and used to identify differentially methylated regions (DMRs). While a good concordance has been described among DMR callers when comparing cell or tissue types^26^, whether this is also the case for lower effect sizes in behavioral neuroscience has been poorly tested. We therefore applied 3 algorithms relying on different strategies: MethylSig^27^, DSS^27,28^, and dmrseq^26^ (Fig.1b-c,S2). Because overlaps among the 3 tools were surprisingly low (with Jaccard Indices, JI, at 16% and 15% for DMR identified by at least 2 methods in females or males, respectively), we evaluated their convergence for DMR simulated in silico at increasing effect sizes, for comparison (Fig.S3). Results showed that, for DNA methylation differences between 5 and 20% (equivalent to those observed for morphine), choosing DMR identified by at least 2 methods maximized predictive power (F1-score). Consistently, morphine-induced DMR prioritized using this criteria were more significant, composed of more CG, and showed larger methylation differences than those identified by a single method (Fig.S4). This resulted in 4209 and 3954 consensus CG-DMR in females and males, respectively, indicating comparable effect sizes of morphine (Fig.1d).

**Figure 1.**
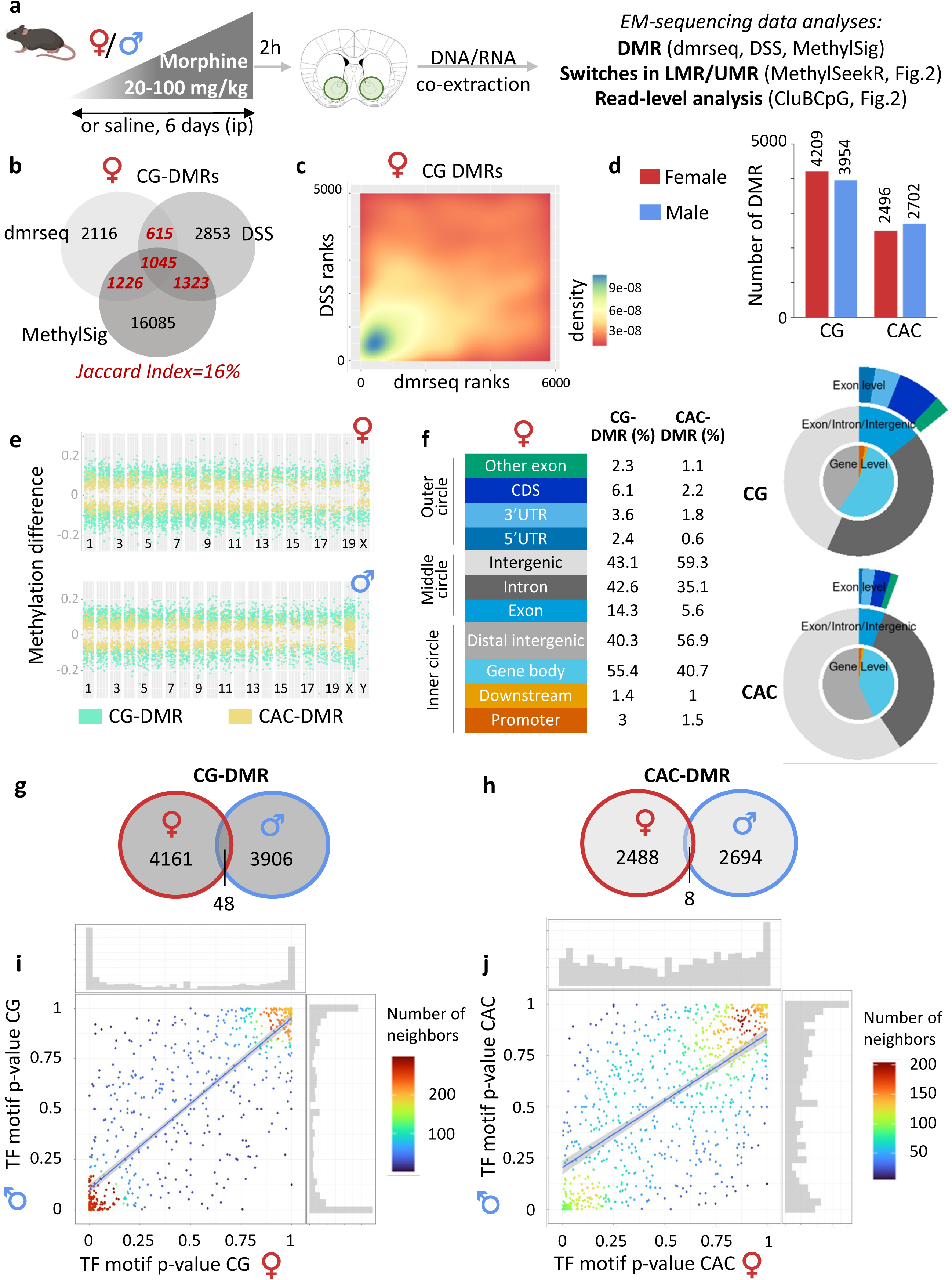
Chronic morphine induces widespread but sex-specific changes in DNA methylation. **a.** Experimental timeline. **b.** Differentially methylated regions identified in female mice in the CG context (CG-DMR) using dmrseq, DSS, and MethylSig. **c.** Density plot of ranks of DMR identified in common by both DSS and dmrseq (Female, CG context; rho=0.3, p=5e-35). **d.** Number of consensus DMRs (see main text). **e.** Genomic distribution and methylation differences of DMR. **f.** Distribution of DMR along genes in females. **g-h.** Venn diagrams of DMR identified across sexes in each cytosine context. **i-j.** Correlation of transcription factor motifs enrichment in CG-and CAC-DMRs (CG: p=4e-231; r=0.85; CAC: p=6e-97; r=0.64).

We next performed a similar analysis for non-CG methylation, considering that we^29^, and others^30–32^, have provided evidence that this neuron-enriched epigenetic substrate may contribute to neuropsychiatric disease^6,33^. We focused on the CAC context, where non-CG methylation was most abundant (Fig.S1f), as expected^31,34–36^. Using the same approach as for CG methylation, consensus DMR identified by 2 methods (Fig.S4g-l) were again prioritized, resulting in 2496 and 2702 CAC-DMRs in females and males, respectively (Fig.1d,S2b-c,S4g-l). As such, building on the recent association between non-CG methylation and OUD in the human brain^33^, we provide direct evidence for opioid-mediated regulation of this epigenetic mark under controlled settings. Of note, no DMR were identified with our consensus approach when considering contexts with lower non-CG methylation levels (data not shown), likely reflecting an unfavorable signal-to-noise ratio at our sequencing depth.

CG-and CAC-DMR showed bidirectional changes (Fig.1e), indicating that methylation and demethylation pathways were both recruited. The affected loci were distributed across all chromosomes (Fig.1e), and enabled clustering of samples based on morphine exposure (Principal Component Analysis, PCA, Fig.S5b-e). Their localization and distance with respect to genes were context-specific (Chi2, p=5e-38, Fig.1f,TableS1): CAC-DMR were predominantly located in distal intergenic regions, while CG-DMR were more frequent in gene bodies, with a higher proportion within exons. Consistently, the distance between DMR and nearest TSS was higher for CAC-(157 kb) than CG-DMR (112 kb; p<1e-16, Fig.S5f). This context-specific distribution of DMR was similar across females and males (Chi2, CG: p=0.25, CAC: p=0.93; Fig.S5g), suggesting properties inherent to each cytosine context (see also below). Next, we determined overlaps among loci affected in each sex. Surprisingly, only 48 (JI=0.6%) and 8 (0.15%) intersections were identified among female and male DMR in the CG or CAC contexts, respectively (Fig.1g-h). This dichotomy was strengthened by the fact that, when DMR for a given context were considered collectively, those identified in one sex showed no differences in DNA methylation in the other (Fig.S6a-b). Although rare, these overlaps were nevertheless more frequent than expected by chance (CG: p=1e-5; CAC: p=5e-3), reflecting the vast number of cytosines interrogated by EM-seq. In addition, CG-or CAC-DMR in one sex were located closer than expected by chance to their counterparts in the other (CG: p=1e-5; CAC: p=1e-3; 100,000 permutations, regioneR; Fig.S6c-d). We next wondered whether these sex-specific morphine effects occurred at loci marked by pre-existing differences among females and males. DMR were identified by comparing female and male saline groups using our consensus approach. These sex-DMR were found predominantly in the X chromosome, reflecting its inactivation (XCI), and mostly corresponded to decreased DNA methylation in females (Fig.S6g-h), consistent with recent data^37^. In contrast, morphine-DMR were mostly in autosomes and rarely overlapped with sex-DMR (1.3 and 1.6% of female and male morphine CG-DMR; Fig.S6e-f), suggesting indirect relationships between sex and morphine effects. Overall, results indicated that chronic morphine triggered highly sex-specific methylomic changes not predicted by baseline sex differences.

Next, we wondered which transcription factors (TF) might be modulated by morphine, and evaluated enrichment of binding motifs in DMR, using TFmotifView^38^. Interestingly, despite low genomic overlap (Fig.1g), female and male CG-DMRs were nevertheless enriched for similar motifs (Fig.1i). This was also observed in the CAC context (Fig.1j) while, in contrast, no correlation was detected within a given sex when confronting the 2 contexts (Fig.S7a-b). Therefore, enrichments of TF motifs were context-but not sex-specific, as illustrated by top findings (Fig.S7c): 7 of the 10 TF most enriched in CG-DMR were common to males and females, among which none were among the top 10 CAC-DMR motifs. Reinforcing this dissociation, no changes in CAC methylation levels could be identified as a function of morphine at CG-DMR, or vice-versa (Fig.S7d-e). Some of these TF have been previously studied in relation to fentanyl self-administration (Tfdp1/E2f^39^), cocaine exposure (Zic1^40^), or dopaminergic (DA) function (Gmeb1^41^), while others remain to be explored (Anrt, Zbtb14). Overall, chronic morphine induced methylomic adaptations at distinct loci in each cytosine context and sex, and modulated the binding of TF that were partly similar across sexes but, to some extent, specific to each cytosine context.

To define morphine effects more comprehensively, we leveraged 2 additional strategies. The first one used methySeekR^42,43^ to identify 2 methylomic features: CG-rich, unmethylated regions (UMR), and CG-poor, lowly methylated regions (LMR; Fig.2a,S8a-c). As expected, UMR overlapped with promoters (>70%), while LMR were intronic or intergenic (>80%) and showed higher inter-individual variability (Fig.S8d-f), consistent with their description as distal regulatory elements. Then, we identified regions where UMR/LMR were gained or lost as a function of morphine (Fig.2b), resulting in 177 diffUMRs and 524 diffLMRs in e.g. females. Compared to DMR, which by construction show high and homogeneous methylation differences across groups, diffLMR/UMR exhibited milder and more variable differences (Fig.S8g-h). Interestingly, while these regions rarely overlapped with DMR, they were sufficient to cluster saline and morphine groups, supporting their informational value (PCA, Fig.2c,S8i-k). For the second strategy, we analyzed single-read methylation patterns using CluBCpG. This approach, developed for cellular deconvolution^44,45^ or applied to brain maturation^46^, is based on the rationale that binary methylation states along successive CG sites of single reads (i.e., single cells) unravels differences that remain masked when methylation is averaged across reads, such as in DMR calling (Fig.2d). Compared to CluBCpG’s default value (100-bp), we considered 150-bp genomic bins (to match the length of our reads), leading to an increase in the complexity of methylation patterns (Fig.S9a). We then identified bins where changes in the frequency of such patterns were induced by morphine, defined as “diffBins”. This resulted in 8625 and 11604 diffBins in females and males, respectively (Fig.2e,S9b). Interestingly, these regions rarely intersected with DMR or with diffLMR/UMR (Fig.S9c-d), showed distinct distributions with respect to genes (Fig.S9e,TableS2), and exhibited methylation levels that, when averaged across reads, poorly separated saline and morphine groups (Fig.S9f-g), supporting the notion that they convey a distinct biological signal. Importantly, these 2 strategies again identified distinct genomic sites across sexes (Fig.S8l-m,S9h), strengthening DMR results.

**Figure 2.**
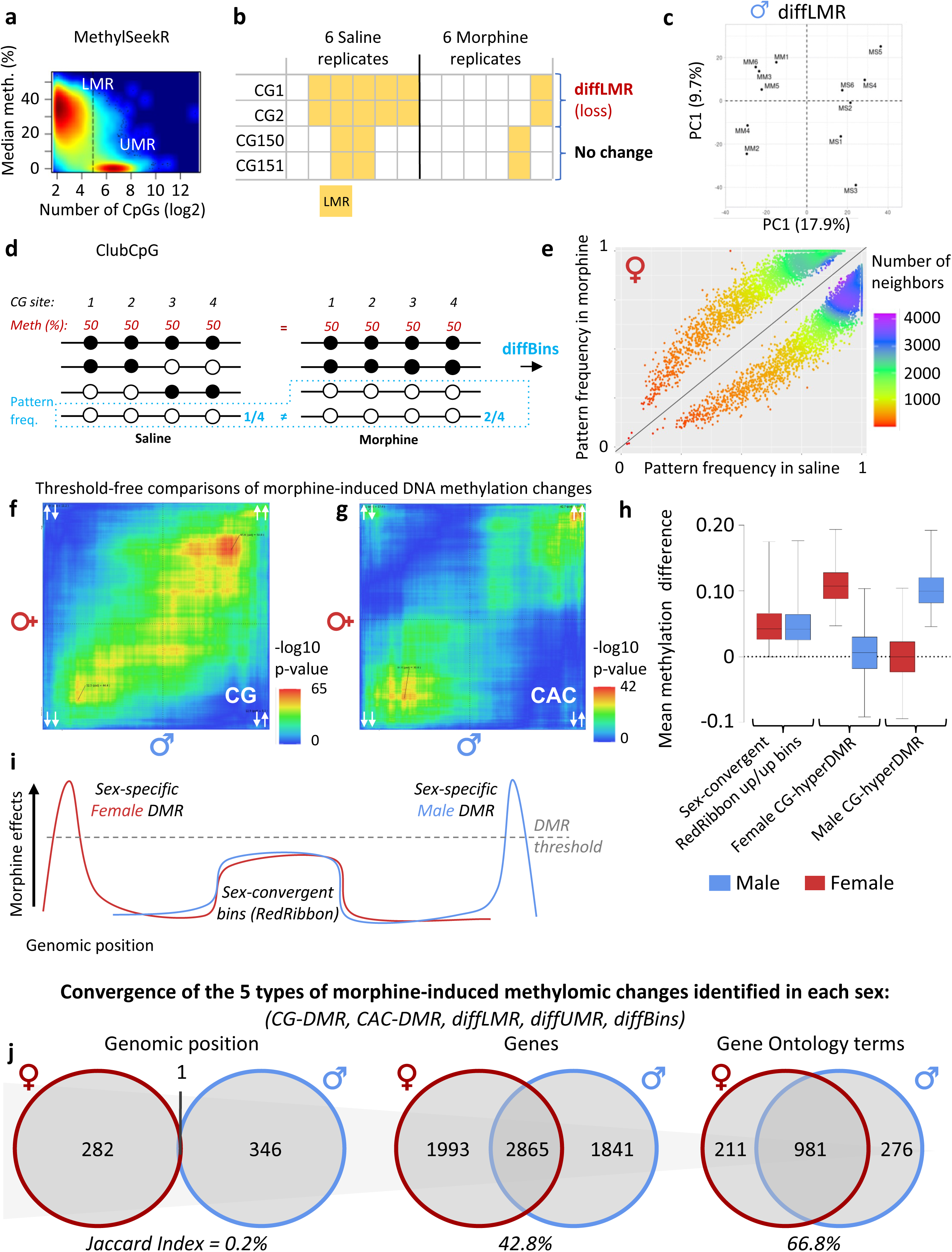
Chronic morphine induces sex-specific changes in regulatory elements and single-read methylation patterns that converge on similar genes and biological pathways. **a.** LMR and UMR identified by methylseekR for one sample. **b.** Approach for the identification of morphine-induced switches in LMR (diffLMRs) or UMR (diffUMRs) status. **c.** PCA of DNA methylation levels at diffLMRs. **d.** Approach for the identification of morphine-induced changes in single-read methylation patterns. **e.** Changes in patterns (DiffBins) in females. **f-g.** Threshold-free comparisons of morphine effects in males and females in CG (f) or CAC (g) contexts, using RedRibbon. **h.** Morphine-induced methylation differences observed in each sex at female CG-DMR, male CG-DMR, and genomic bins showing sub-threshold increase in DNA methylation in both sexes (up/up, RedRibbon). **i.** Schematic of morphine effects. **j.** Convergence of morphine-induced DNA methylation changes.

Because of these surprising sex differences, we then implemented a threshold-free approach to seek patterns of similarity. To do so, we used RedRibbon^47^, a rank-rank hypergeometric method recently developed for large datasets. Applied to morphine-induced DNA methylation changes (computed in ∼4.5 million non-overlapping 500-bp bins throughout the genome), this uncovered a significant convergence across sexes (Fig.2f-h), with numerous bins impacted by morphine in similar directions: in the CG or CAC contexts, most significant overlaps involved 163,925 or 6,340 bins showing increased methylation in both females and males, respectively (upper-right quadrant; Fig.2f-g). These sex-convergent bins showed milder methylation differences than DMR (Fig.2h,S10e), were 10 times closer to DMRs than expected by chance (e.g. in males: 3.1 kb observed compared to 40.6 kb expected; 100,000 permutations, p=1e-5 in both sexes; Fig.S10f-g), and were also frequently located between pairs of nearest male and female DMRs (1,000 permutations, p=1e-4 in the CG context). Altogether, these results indicate that morphine drove broad but sub-threshold methylomic changes at similar genomic sites across females and males (RedRibbon bins), accompanied by less frequent, more localized and sex-specific adaptations (DMR; Fig.2i).

### Functional convergence of morphine-induced sex-specific methylomic plasticity

To characterize their functional relevance, we associated DMR with genes located in *cis,* using GREAT^42^. Compared with JI observed for genomic coordinates, overlaps across sexes of the genes associated with DMR increased to 32% (CG) and 29% in the CG and CAC contexts, respectively (Fig.S11a). This indicated that morphine-induced methylomic changes affected partially overlapping genes across sexes. Gene ontology (GO) terms associated with these genes showed an even greater convergence, with JI rising to 58% (CG) and 53% (CAC), involving terms related to the regulation of membrane potential, synaptic organization, small GTPase activity or Wnt signalling, among others (Fig.S11a,TableS3). To determine whether this GO-level overlap was solely due to shared DMR-associated genes between sexes, we performed permutation testing using random gene lists with equivalent male-female overlap as in our results (Fig.S11b). Strikingly, while expected JI for GO terms averaged 12.0% (CG) and 8.7% (CAC), observed values were 5-to 6-fold higher (CG, p=5e-148; CAC, p=5e-144). This indicated that genes associated with DMR in only one sex were enriched in similar biological functions as those shared across sexes. Thus, although chronic morphine treatment induced DMR at distinct loci in males and females, these effects progressively converged on overlapping genes and biological functions.

In addition to this sex convergence, another one emerged across the 2 cytosine contexts. While rare intersections were observed among CG-and CAC-DMR (e.g. in females, N=17, JI=0.25%), overlaps among associated genes (N=1595; JI=25%) and GO terms (N=826; JI=45%) strongly increased (with similar results in male mice, Fig.S11c). These findings are consistent with our previous analyses of DMR distribution relative to genes, as well as of TF enrichment (Fig.1f,S5f-g,S7), and reinforce the hypothesis of context-specific regulatory mechanisms affecting partly similar genes and biological processes.

Finally, we combined results from the 5 methylomic features: CG-DMR, CAC-DMR, diffUMR, diffLMR, and diffBins. In females (Fig.S12a), only 1.8% of genomic sites were annotated by 2 features, and none by more. In contrast, 45% of associated genes and 61.1% of GO terms were annotated by 2 or more features. Similar findings were observed in males (Fig.S12b). Importantly, a progressive convergence again emerged across sexes (Fig.2j), with low intersections at genomic coordinates (0.2%) that strongly increased for genes (42.8%) and GO terms (66.8%), and again involved biological pathways related to synaptic plasticity, similar to those mentioned above for DMR (TablesS3-4). Overall, while previous work had characterized opioid-induced transcriptomic changes affecting such pathways, here we uncover the underlying methylomic plasticity. Furthermore, our results show that this plasticity occurs at sex-specific loci that converge on common genes and functions.

### Morphine-induced changes in histone modifications

Because we were surprised by the extent of sex differences, we opted to strengthen our findings by investigating additional epigenetic layers. We used CUT&Tag-Sequencing to analyze 5 histone marks with well-characterized roles in chromatin regulation (n=6 replicates for saline/morphine and male/female groups, n=120 libraries total). After peak calling, we observed that the data clustered according to the functional impact of each mark (repressive/activating), followed by the modification type, and sex (Fig.3a,S13a-c). As expected^29,48,49,50^, the repressive marks (H3K27me3, H3K9me3) were less abundant in gene bodies than activating ones, while H3K27ac and H3K4me1 were enriched around the transcription start site (TSS) and H3K36me3 progressively accumulated towards the transcription end site^50^ (TES, Fig.3b). Each mark also exhibited distinct positive or negative correlation with gene expression (Fig.3c).

**Figure 3.**
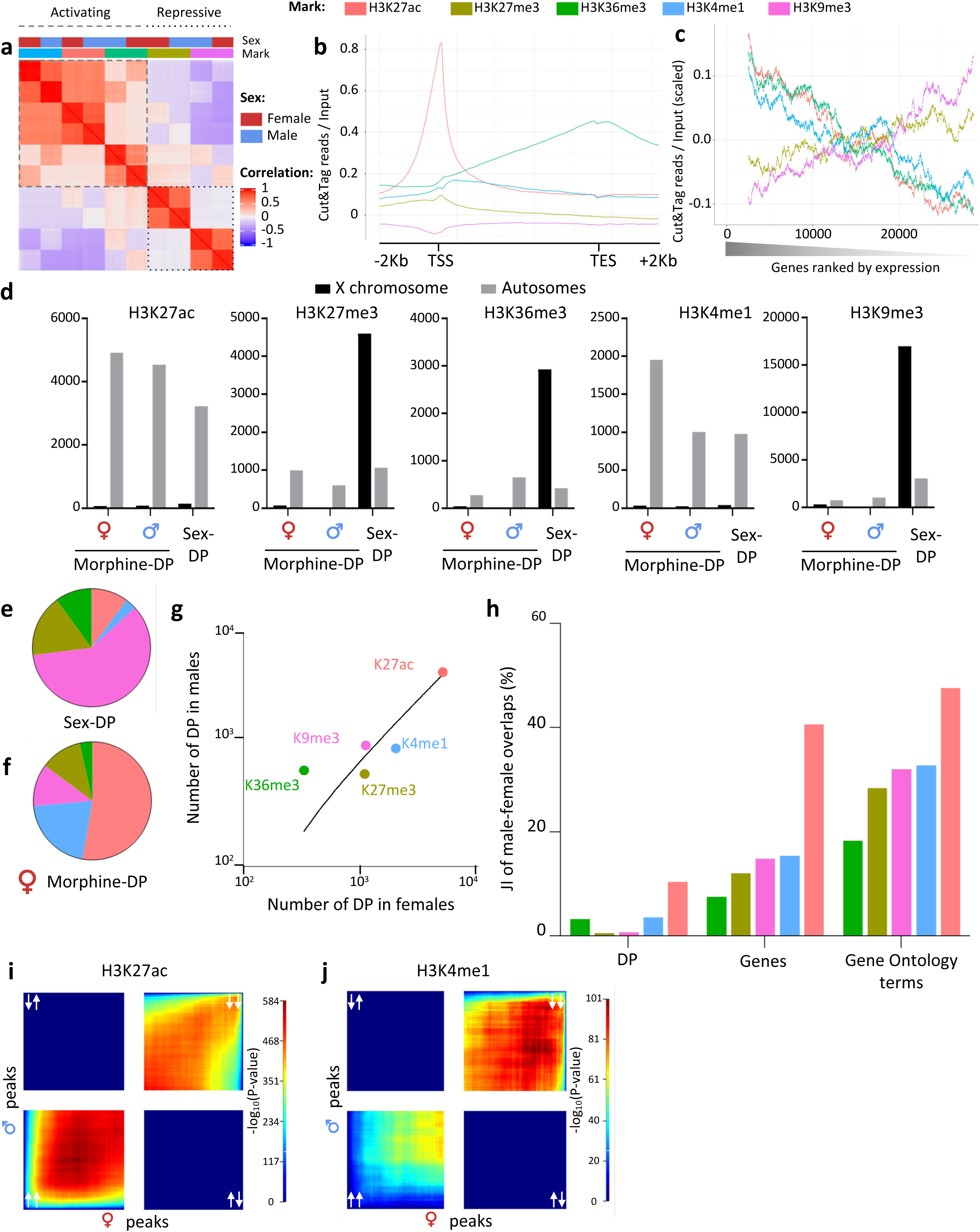
Chronic morphine induces sex-specific changes in histone modifications that show functional convergence. **a.** Clustering of histone data. **b.** Gene profiles of histone marks. **c.** Abundance of histone marks according to gene expression. **d.** Number of differential peaks (DP) identified as a function of sex or morphine. **e-f.** Proportions of each mark among sex-DP (e) and morphine-DP (f). In females, H3K27ac was the most affected by morphine (53%), followed by H3K4me1 (21%), H3K9me3 (12%), H3K27me3 (11%), and H3K36me3 (3%). **g.** Number of morphine-DP across sexes. **h.** Convergence of morphine-DP by mark. **i-j.** Threshold-free comparisons of morphine effects across sexes.

Next, similar to DNA methylation analyses, we identified changes as a function of sex or morphine (differential peaks, DP; p<0.01; Fig.S13d,TableS5). Much stronger effects of sex were observed for 3 marks (H3K27me3, H3K36me3 and H3K9me3), with DP mostly located on the X chromosome and corresponding to increased abundance in females, reflecting XCI (Fig.3d-e,S13e-f). Of note, while the accumulation of H3K27me3 and H3K9me3 on the inactive X chromosome of females is well-documented^51,52^, to our knowledge, the implication of H3K36me3 has not been described. Regarding morphine, the 2 marks most strongly impacted (H3K27ac, H3K4me1) were those that were the least involved in sex differences (Fig.3e-f). In females, for example, H3K27ac represented 53% of morphine-DP but only 5% of sex-related ones, while H3K9me3 accounted for 63% of sex-DP but 12% of morphine ones. This likely reflected the distinct biological processes involved, with sex differences originating from XCI during embryogenesis, and morphine effects from an experience during adulthood. A significant correlation was observed across sexes for the number of morphine-DP identified for each mark (r=0.96, p=0.008; Fig.3g), uncovering a conserved hierarchy in their sensitivity to this drug. Importantly, numerous similarities with previous DNA methylation results were observed. First, despite comparable effect sizes and distributions with respect to genes (TableS5), morphine-DP showed little overlaps across sexes (JI from 0.6% for H3K27me3 to 10.5% for H3K27ac; Fig.3h), as well as rare intersections with sex-DP (identified when comparing male and female control groups, Fig.S14), indicating that the principle of epigenetic divergence described above for DNA methylation extends to histone modifications. Second, threshold-free comparisons (using RRHO2) showed that, in addition to sex-specific morphine-DP, milder changes also affected broader genomic sites that were partly shared across sexes, in particular for H3K27ac (p=4.0e-585) and H3K4me1 (p=1.3e-88; Fig.3i-j). These overlaps were lower for the other marks less sensitive to morphine and primarily implicated in sex differences (H3K36me3, p=3.2e-45; H3K9me3, p=5.0e-9; H3K27me3, p=2.5e-6; Fig.S15a-c). Third, a functional convergence also emerged (Fig.3i,TableS6), as JI again increased for morphine-DP-associated genes (JI from 8% for H3K36me3 to 41% for H3K27ac) and related biological functions (JI from 8% for H3K36me3 to 56% for H3K27ac). Fourth, this convergence of GO terms was not only due to genes annotated to DP in both sexes, but reinforced by sex-specific ones (with JI 3.7 to 12.1 times higher than expected; p<7e-10 for all marks, Fig.S15d). GO terms enriched for these DP were related to the regulation of membrane potential, synapse organization or small GTPase activity, and highly correlated with GO terms uncovered during DNA methylation (r=0.86, p=6e-105; Fig.S15e). Altogether, this demonstrated that, in addition to the DNA methylome, chronic morphine drove highly sex-specific adaptations across a broad array of histone modifications, with downstream convergence.

Next, we wondered whether histone data might deepen the understanding of morphine effects on DNA methylation. An unsupervised k-means approach was used to identify CG-DMR with different histone combinations, resulting in the identification of 3 clusters (Fig.4a-d): the largest one, Cluster1, was defined by low levels of 2 activating (H3K27ac, H3K4me1) and 1 repressive (H3K27me3) marks (n=3130 in e.g. females), corresponding to poised enhancers; Cluster2, by higher levels of the 2 activating marks (n=829), defining active enhancers^53^; Cluster3, by broad H3K27me3 domains corresponding to heterochromatin (n=238). Consistently, genes annotated to Cluster2 showed highest expression, and Cluster3 the lowest (Fig.4e). These subgroups of DMR also exhibited distinct distributions along genes: Clusters1 and 3 were present in both introns and intergenic regions, while Cluster2 were mostly intronic (Fig.4f). Morphine may act at these Cluster2 intronic enhancers by modulating MeCP2 binding^54^ and the activity of the N-CoR complex^55^, an hypothesis that warrants further work.

**Figure 4.**
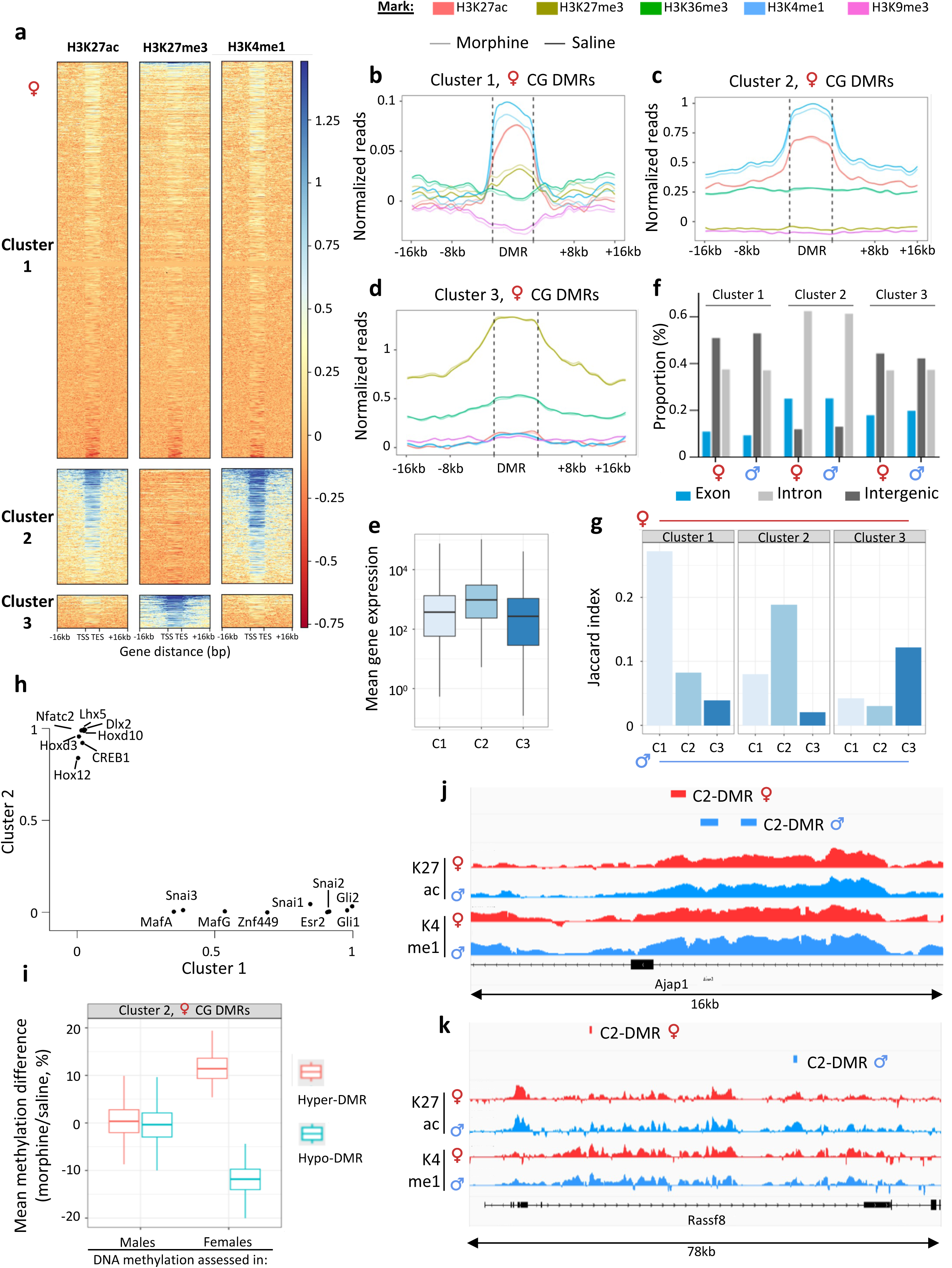
Histone marks differentiate 3 types of morphine-induced methylomic adaptations that are conserved across sexes. **a.** Heatmap of histone marks across 3 clusters of female CG-DMR. **b-d.** Profiles of histone marks for each cluster of female CG-DMR. **e.** Mean expression of genes associated with each cluster of DMR, in females (ANOVA: p=5.6e-10, p(C1vsC2)=5.1e-10, p(C1vsC3)=0.93, p(C2vsC3)=4.4e-4). **f.** Genic distribution of each cluster of DMRs. **g.** Overlap across sexes of genes associated with DMR from each cluster. **h.** Transcription factor motifs enrichment in cluster1-versus cluster2-CG-DMR. **i.** Morphine-induced methylation differences observed in each sex at female cluster2-CG-DMRs. **j-k.** Examples of sex-specific morphine-DMR falling within a common (j), or distinct (k), enhancer.

Importantly, the same clusters were observed in a similar analysis in males (Fig.S15f-h), and genes annotated to a given cluster and sex were enriched for genes associated with that same cluster in the other sex (Fig.4g). These reproducible clusters therefore likely reflect different mechanistic cross-talk among DNA methylation and histone marks. Consistently, an analysis of binding motifs restricted to cluster-specific DMR identified distinct sets of TF (Fig.S15i). Among these, 7 TF were enriched in Cluster1-DMR in both females and males, and 9 in Cluster2, while none of these were common to both Clusters (Fig.4h). While CREB1 has been extensively studied in the addiction field^56^, others have been primarily investigated in cancer (Gli, Maf) or neurodevelopment (Dlx2, Hoxd) and represent interesting targets for future work. Finally, similar to the full set of DMR (Fig.2h), genomic loci corresponding to cluster-specific DMR in one sex showed no DNA methylation differences in the other (Fig.4i,S15j), but nevertheless exhibited similar histone profiles (Fig.S15k-l). This reflected 2 situations coexisting at similar frequency: either morphine triggered methylomic changes at different subregions within common enhancers (clusters 1 and 2; Fig.4j) and heterochromatic domains (cluster3); or, alternatively, distinct enhancers and domains were affected by morphine in each sex (Fig.4k), thereby suggesting a heterogeneity in the molecular mechanisms underlying the epigenetic sex divergence. Altogether, our results show that chronic morphine recruited various types of DNA methylomic adaptations, defined by distinct histone landscapes, reproducible across sexes yet targeting sex-specific loci, and modulating specific TFs.

### No evidence for sex differences in histological organization or morphine-induced neurophysiological adaptations in the NAc

The extent of these sex differences prompted us to explore underlying factors. Opioids classically act in the NAc through direct and indirect mechanisms: direct effects involve recruiting the inhibitory MOR expressed by medium spiny neurons (MSN); indirect ones recruit MOR expressed by GABAergic interneurons of the ventral tegmental area, leading to disinhibition of the mesolimbic pathway and increased DA release, with distinct impact on MSN expressing Drd1 (D1) or Drd2 (D2) DA receptors^57^. We reasoned that under-appreciated sex differences in these factors may contribute to the epigenetic divergence described above.

First, we conducted multiplex in situ hybridization to analyze D1-, D2-, or MOR-expressing MSN populations following morphine treatment. Coronal sections were scanned (n=16 sections/group/sex; Fig.5a,S16a) and >300,000 cells analyzed with QuPath (Fig.S16b; *Methods*). D1+, D2+, and D1+/D2+ MSNs accounted for 56%, 28% and 16% of all MSNs, respectively (Fig.5b), consistent with previous work^58^. No significant changes were observed as a function of sex or morphine for any of these populations, which typically account for >90% of NAc neurons^59^. Next, we focused on the less abundant MOR-expressing MSNs, the direct cellular targets of morphine. We observed that double D1+/D2+ MSN expressing MOR were (Fig.S16c-e): i) more abundant in the rostral NAc; ii) more frequent (23%) than double D1+/D2+ neurons within the full MSN population (16%), and iii) expressed MOR at higher levels than MSN expressing a single DA receptor. While these single-receptor populations have been extensively studied^60,61^, our data highlight a potentially important role for the comparatively less explored population of double D1+/D2+ MSN in opioid responses, as recently shown for cocaine^62^. Importantly, neither sex nor morphine affected the number of MOR-expressing MSN (Fig.5c). Thus, morphine epigenetic effects did not reflect broad changes in NAc histological composition.

**Figure 5.**
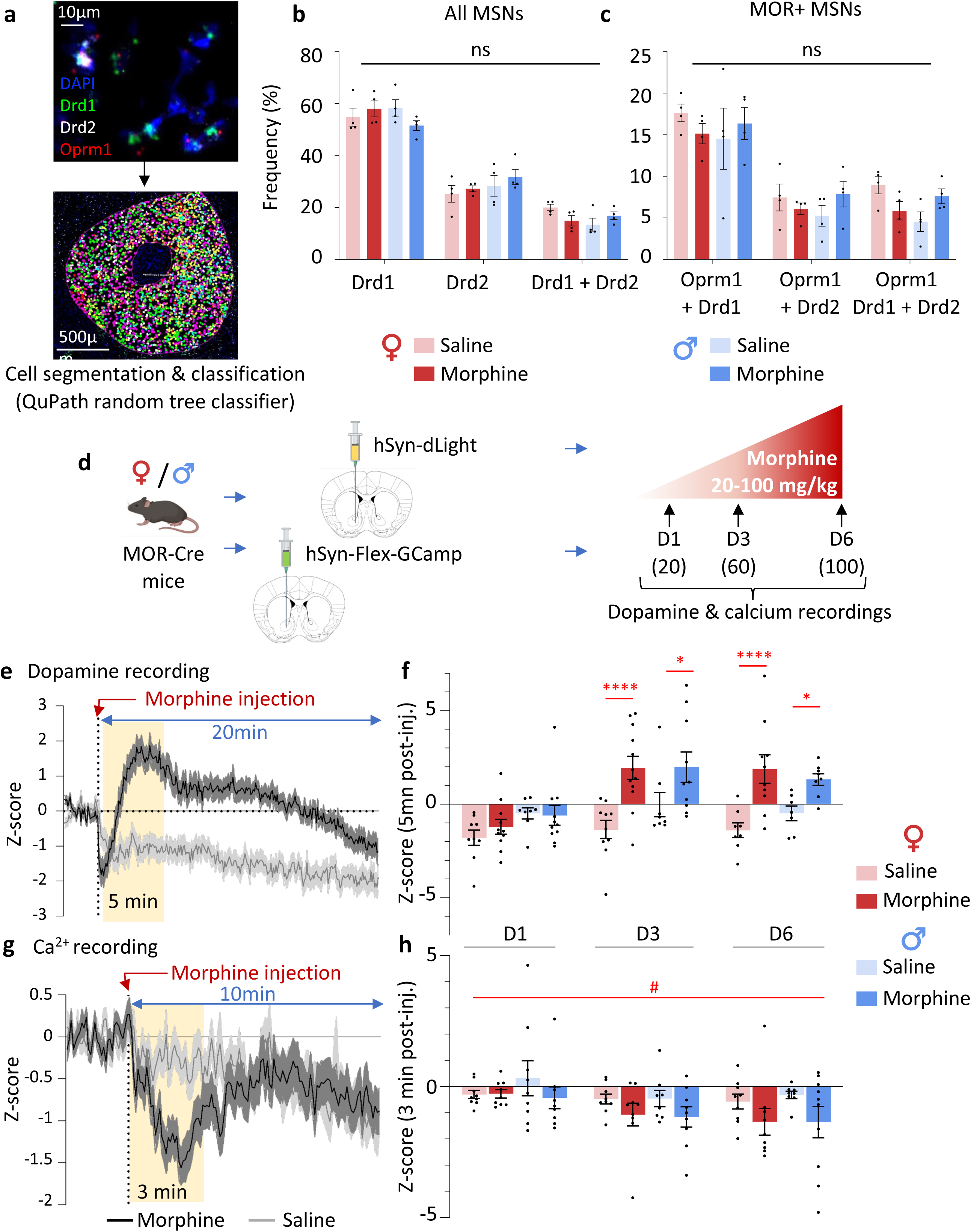
No sex differences in histological organization or morphine-induced neurophysiological adaptations in the nucleus accumbens. **a.** In-situ hybridization for Drd1, Drd2 and Oprm1. **b.** Frequency of Drd1+, Drd2+ and Drd1+/Drd2+ medium spiny neurons (MSN). No differences were observed as a function of sex (3-way ANOVA, p=0.68) or chronic morphine(p=0.68). **c.** Similarly, the frequency of Oprm1+ MSN was not significantly affected by either sex (p=0.54) or morphine (p=0.95). **d.** Fiber photometry experiments. **e.** Average DA release in response to morphine or saline injection. **f.** DA release at Day1 (D1), D3 and D6 (3-way ANOVA; time, p<0.0001; morphine, p<0.0001; sex, p=0.21; post-hocs: female morphine vs female saline, D1 p=0.43, D3 p<0.0001, D6 p<0.0001; male morphine vs male saline, D1 p=0.97, D3 p=0.01, D6 p=0.03). **g.** Average calcium activity in mu opioid receptor-expressing (MOR+) neurons in response to morphine or saline injections. **h.** Calcium activity in MOR+ neurons at D1, D3 and D6 (3-way ANOVA; time, p=0.03; morphine p=0.02; sex p=0.67). ns, non-significant; *p<0.05; ****p<0.0001; # significant effect of day and treatment.

Next, we used fiber photometry and the dLight biosensor^63^ to record DA release in the NAc of freely moving mice during morphine treatment: at day 1 (D1), D3 and D6 (Fig.5d). An initial ∼30-second decrease in DA signal was observed in both saline and morphine groups (Fig.5e), likely reflecting injection-related stress, as described for other acute stressors^63,64^. This was followed by a strong and progressive increase in DA release that occurred selectively in morphine-treated mice, particularly during the first 5 minutes, reflecting the pharmacological effect of the drug. During that period, days and morphine treatment had significant effects on DA release (Fig.5f), with an interaction between factors but, importantly, no sex differences. Thus, while opioid-induced DA release had previously been measured in males using microdialysis^65^ or biosensors^66^, we found that the magnitude of this physiological response did not differ among sexes.

We then recorded the activity of neurons recruited by morphine. A neuron-specific Cre-dependent viral vector expressing the calcium GCaMP biosensor was injected in the NAc of MOR-Cre mice (Fig.S16f), which expressed the Cre recombinase under control of the MOR locus^67^. Across the 3 recordings (D1,3,6), a decrease in Ca2+ signal was observed in morphine-but not saline-treated mice. This effect progressively increased during the first 90 seconds following each injection, and largely disappeared after 3 minutes (Fig.5g). During this period, results showed significant effects of days and morphine, without any effect of sex nor any interaction (Fig.5h). Therefore, morphine directly inhibited the activity of MOR+ neurons, consistent with the well-characterized Gi/o coupling of the receptor. This inhibition rapidly regressed, probably reflecting secondary effects of DA^68^ and homeostatic processes among basal ganglia^69^. Importantly, no sex differences were detectable for opioid-induced inhibition. Overall, these experiments indicate that the cellular composition of the NAc was similar across sexes and was not modified by morphine, and that comparable physiological adaptations affecting DA release and Ca2+ signaling occurred. Hence, none of these factors accounted for epigenetic differences between females and males.

### Chronic morphine triggers sex-specific transcriptional adaptations that partly recapitulate the human signature of OUD

We then investigated the transcriptomic impact of morphine-induced epigenetic changes, and conducted bulk RNA-seq (Fig.S17a-b) in combination with a new strategy for cell-type specific analysis of MOR+ neurons. In bulk tissue, 1073 and 1546 differentially expressed genes (DEG) were identified in females and males, respectively (adjusted p<0.1; Fig.S17c,TableS7), with 459 common DEG (JI=21%, Fig.6a), exceeding overlaps observed for DNA methylation or histone marks. Using threshold-free comparisons, a transcriptional concordance stronger than those observed for epigenetic modifications was also observed (p=1e-394 for up/up-regulated genes; Fig.6b,S17d-f), except for H3K27ac. The DEG were enriched in GO terms such as Wnt signaling, steroid hormones, or synaptic regulation (TableS8), and correlated with terms previously associated with DNA methylation (r=0.61, p=2e-17) and histone (r=0.62, p=9e-19) changes. To document the translational relevance of these results, we compared them with a recent human study that investigated NAc tissue from individuals with OUD and controls^24^. Improving on previous work^6^, this study considered equal numbers of women and men, although the impact of sex was controlled for, but not directly assessed. We therefore reprocessed the data and conducted differential expression analysis, followed by cross-species comparisons with our mouse results. When considering pooled females and males (Fig.S17g), significant similarities were identified (p=7.9e-14 and p=4.0e-10 for up/up- and down/down-regulated genes, respectively). Interestingly, analyzing each sex separately revealed further dissociations as, despite the decrease in sample size, significance levels slightly decreased in males (p=3.2e-20 and p=2.5e-20 for up/up- and down-down-regulated genes, respectively; Fig.6c) but increased in females (p=4.0e-7 and p=2.0e-10 for up/up- and down/down-regulated genes, respectively; Fig.6d,TableS9). While part of the GO terms involved in these overlaps were common to all comparisons (pooled sexes, female- or male-specific; Fig.6e), such as axon development, others were sex-specific, such as regulation of neuron apoptotic process in males, or regulation of neuron differentiation in females. Overall, these results indicated that our morphine regimen partly recapitulated sex-specific transcriptional signatures of OUD in humans.

**Figure 6.**
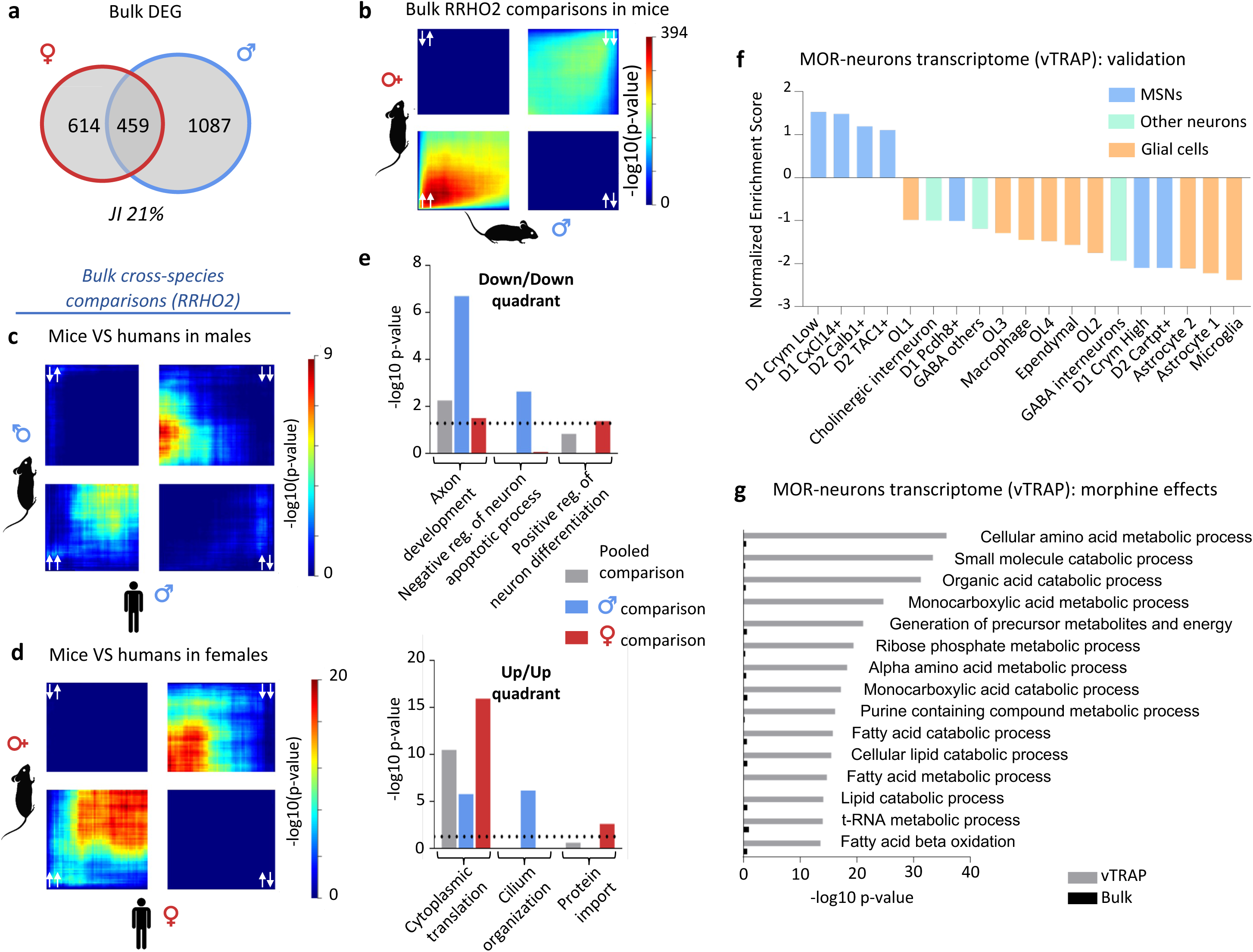
Chronic morphine induces sex-specific transcriptional adaptations in bulk tissue and morphine-responsive neurons. **a.** Overlap of morphine-induced DEG in females and males (bulk tissue). **b.** Threshold-free comparison of transcriptional morphine effects across females and males (RRHO2). **c.** Comparison of morphine effects in male mice and men with opioid use disorder (OUD)^24^. **d.** Comparison of morphine effects in female mice and women with OUD. **e.** GO enrichments of best RRHO2 overlaps (down/down and up/up quadrant) across the 3 mouse/human comparisons: using both sexes, in males, or in females only. **f.** Validation in naive mice of the capture of the transcriptome of mu opioid receptor-expressing (MOR+) neurons, using vTRAP. Results show enrichment and depletion of gene sets defining nucleus accumbens cell-types^71^), computed using GSEA. **g.** GO enrichments of best RRHO2 overlap (down/down quadrant; Fig.S19d) obtained when comparing morphine transcriptomic effects on male and female MOR+ neurons (with corresponding enrichments in bulk tissue, for comparison).

To deepen our cellular resolution, we then investigated the “translating transcriptome” of MOR+ neurons by combining the MOR-Cre line and viral translating ribosome affinity purification (vTRAP; Fig.S18a)^70^. As validation, we used RNA-seq to compare RNA fractions isolated from tagged ribosomes of MOR+ neurons (Immunoprecipitated, IP) or extracted from bulk tissue (Input), in drug-naive mice (n=3/fraction). Results were processed for Gene Set

Enrichment Analysis (GSEA) to identify overrepresentation and depletion of marker genes previously defined by single-cell RNA-seq for 26 NAc cell types^71^. As expected, genes defining most D1 and D2 MSNs (∼67% of MSN in^71^) were significantly enriched in the transcriptome of MOR+ neurons (Fig.6f), while markers of glial cells (oligodendrocytes, astrocytes, microglia) or GABA interneurons were depleted. Of note, a few MSN subgroups were also depleted (∼33% in^71^), thereby refining the identification of MSN expressing MOR. Next, we used this vTRAP approach to investigate morphine effects. IP RNA fractions from saline-and morphine-treated mice were first validated by qPCR (with no differences across sexes or treatment groups for enrichment of marker genes, Fig.S18b-e), and processed for RNA-seq and differential expression analysis (TableS7). Transcriptomic adaptations recruited by morphine in MOR+ neurons were again sex-specific, with limited overlap among DEG (pointing to corticosteroid signaling and inflammatory processes, TableS8), accompanied by broader threshold-free similarities (Fig.S19a-d,TableS10). Comparison of these vTRAP and previous bulk results also revealed significant concordance (Fig.S19e-f), indicating that while MOR+ neurons corresponded to restricted MSN subtypes, the adaptations driven by morphine and affecting both MOR+ and MOR-MSN (such as DA release) resulted in correlated transcriptomic changes. Zooming in on MOR+ neurons nevertheless uncovered findings that were not detectable at bulk level (Fig.S19g). Among these was a significant overrepresentation of GO terms related to energy metabolism (Fig.6g), indicating that the previously described opioid-induced metabolic dysregulation^72,73^ may primarily affect MOR+ neurons. Furthermore, the “opioid signaling reactome”, an established gene set related to signal transduction of opioid receptors^74^, was not affected at bulk level (Fig.S17e-f) but showed significant downregulation in MOR+ neurons (Fig.S19h-i). Interestingly, we recently obtained similar results in another brain region, the dorsal raphe nucleus^75^, where the same gene set was downregulated in MOR+ neurons but not in bulk tissue. This selective response to morphine of MOR+ neurons may therefore be conserved across brain regions, a possibility that warrants further investigation to assess its generalizability.

### Multiomic integration highlights gene modules and neuronal cell types impacted by chronic morphine in both sexes

To go further in the functional and cell-type annotation of our data, we performed multiomic integrative analyses. First, we investigated the degree of congruence detectable among omic layers. To do so, we computed permutations and odds-ratios for pair-wise overlaps among genes associated with morphine-induced differences for each one of them. Results showed that, although the narrow histone marks H3K27ac and H3K4me1 are typically associated with transcription, the broad H3K36me3 modification displayed larger and more significant odds-ratios with DEG (Fig.7a,S20a), indicating a stronger link to morphine-induced transcriptomic effects. Second, the vast majority of these overlaps were significant and correlated across sexes (Fig.S20b), although odds-ratio were relatively modest. This partly reflects technical limitations associated with the bulk resolution of our analyses (see also *Discussion*), as well as the challenge of associating epigenetic differences with genes, which may be improved by future consideration of chromatin domains^76^. At the biological level, they also suggest that part of these epigenetic changes may not immediately impact gene expression, but rather contribute to long-term adaptations that unravel upon re-exposure to the drug, or to stimuli associated with its effects^77,78^. Then, we aggregated GO enrichments obtained for each omic layer, using Stouffer p-values. Among other findings, results showed that morphine recruited multiomic adaptations that most significantly affected the histological and molecular organization of the synapse, as well as the regulation of its electrical activity, consistent with single-omic GO enrichments described above, as well as previous knowledge^6^ (Fig.S20c,TableS11).

**Figure 7.**
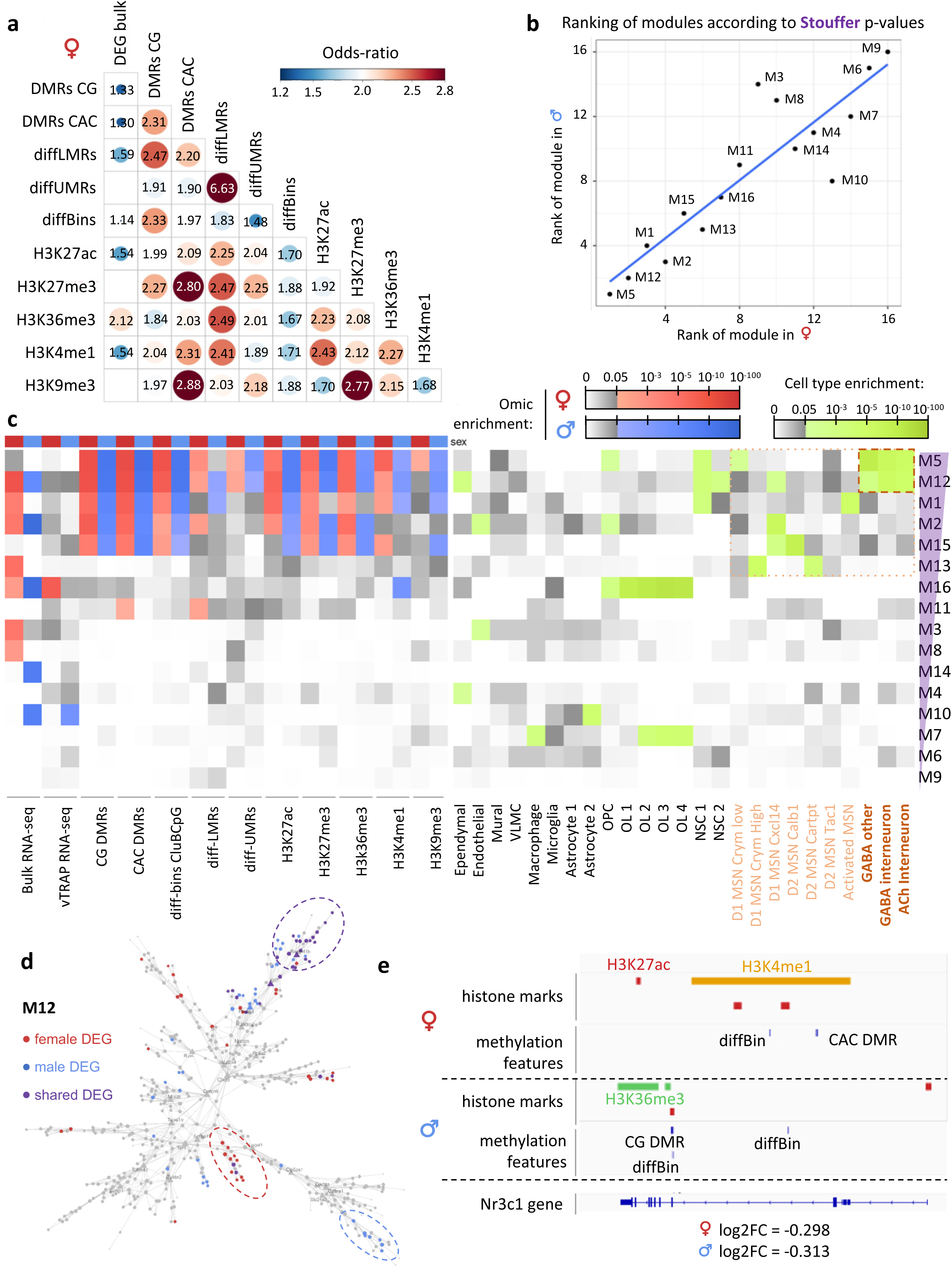
Chronic morphine epigenetically reprograms similar gene modules and cell types in both sexes. **a.** Odds-ratios of pair-wise overlaps among genes associated with morphine-induced differences observed for each omic layer. **b.** Gene co-expression modules were similarly ranked in males and females for enrichment in morphine-induced differences across all omic layers. **c.** Heatmap of module enrichment for morphine-induced differences for each omic layer (left), and gene sets defining nucleus accumbens cell-types^71^. **d.** Visualization of the co-expression structure of the M12 module, annotated with differentially expressed genes (DEG) in each sex. **e.** Representation of morphine-induced epigenetic differences annotated to *Nr3c1*, a DEG in both males and females belonging to the M12 module.

Next, we performed a gene co-expression network analysis using Multiscale Embedded Gene co-Expression Network Analysis (MEGENA). Bulk RNA-seq data were used to generate 16 parent and 111 child co-expression modules, which were then annotated for enrichment in morphine-induced differences for each omic layer. Most pair-wise correlations among layers were significant (Fig.S20d-f), refining at the level of modules the genome-wide results from aforementioned permutations. Interestingly, the modules associated with transcriptomic differences in humans with OUD^24^ correlated slightly better with our mouse methylome and histone data than with our mouse transcriptomic data. This pattern suggests that the epigenetic changes may relate more to long-term transcriptomic adaptations underlying the addiction trait than to the transient state induced by morphine exposure. Next, we summarized all enrichments to prioritize gene modules (using Stouffer p-values, excluding human data). The rankings observed across females and males were strongly correlated, with the same 4 modules most significantly impacted (r=0.90, p<1e-16; modules M5, M12, M2, and M1; Fig.7b). These results therefore reinforce at the level of modules the concept of a functional convergence stemming from sex-specific epigenetic adaptations, previously observed for individual omic layers. Then, we computed the enrichment of modules for marker genes defining NAc cell types^71^. Results showed that the 6 modules most significantly impacted by morphine were enriched for neuronal marker genes (e.g. p=1.2e-9 for “D2 MSN Calb1” and M15; Fig.7c, light orange, TableS12), among which the 3 types of local inhibitory interneurons (“GABA interneuron”, “ACh interneuron”, “GABA other”) were particularly enriched in the 2 best modules, M5 and M12 (e.g. p=2.7e-11 for “GABA other” and M5; dark orange). Therefore, while these local interneurons represent a minority of NAc MSN (∼7-8%^71^), they may be particularly affected by morphine epigenetic effects. Beyond these neuron-enriched modules, M16 showed a strong association with all types of oligodendrocytes and their progenitors (OPC), consistent with recent data suggesting that these cells strongly respond to morphine^71^. Finally, we conducted GO enrichment of modules. Among other findings, the 2 best modules, M5 and M12, were enriched in genes related to synaptic organization and function (e.g. syntaxin and synaptotagmin proteins), including for the glutamatergic synapse (TableS13), suggesting that the well-characterized opioid-induced plasticity affecting glutamatergic synapses in the NAc may particularly involve interneurons^79–81^.

As an illustration, we focused on M12 for a more detailed analysis. This module, composed of 112 genes, included 40 male-specific and 32 female-specific DEGs that affected distinct parts of the module’s co-expression organization, as well as 29 DEG common to both sexes and located elsewhere (Fig.7d). Among the latter was *Nr3c1*, the gene encoding the glucocorticoid receptor, a key regulator of the stress axis and glutamatergic transmission, known to contribute to OUD, that was identified in multiple enrichments across omic layers (TableS11)^82,83,84^. *Nr3c1* was significantly downregulated by chronic morphine and associated with 13 distinct epigenetic adaptations in both sexes (Fig.7e). These adaptations involved different loci and histone marks in each sex, yet resulted in a similar transcriptional outcome. Therefore, this individual gene illustrates the epigenetic divergence and functional convergence revealed at the genome-wide scale in this study.

## 4. Discussion

In the present work, we first analyzed 2 forms of DNA methylation occurring in the CG and CAC contexts. Both have been examined in recent post-mortem studies of individuals with OUD, although these investigations used microarrays or reduced representation bisulfite sequencing that were restricted to portions of the genome^33,85–87^ (see^6^ for review). Here, we conducted broader genome-wide analyses and, under well-controlled experimental conditions, observed substantial plasticity in both contexts in response to morphine. These findings extend previous human results to the full epigenome, and support the hypothesis that the 2 types of DNA methylation may contribute to opioid effects. Importantly, CG-and CAC-DMRs showed little genomic overlap, distinct distributions with respect to genes, and were associated with different histone modifications. These observations reproduce results from our recent study of early-life adversity and depression^29^, where we identified similar dissociations, suggesting that such principles may generalize across psychiatric disorders. Extending beyond those findings, we now show that CG- and CAC-DMR likely modulate distinct sets of TF that nevertheless converge on overlapping genes, co-expression modules, and biological pathways. These results reinforce a model in which CG and CAC methylation (although they are deposited and read by similar proteins, DNA methyltransferases and methyl-CpG-binding domain proteins, respectively^88^), represent mechanistically distinct epigenetic substrates that act synergistically to mediate opioid-induced transcriptional adaptations. Notably, these adaptations partially recapitulated changes reported in the human NAc in OUD, underscoring the translational relevance of the epigenetic plasticity identified here.

The second major aspect of this study relates to sex differences. Over the last decades, such differences have been progressively identified in brain physiology at multiple levels, including in terms of transcriptomic expression, cell type composition, histological organization, or connectivity among brain regions^89–91^. Surprisingly, however, a significant bias towards the study of male subjects persists in the preclinical literature^6,92,93^, hindering progress. This is for example illustrated by the fact that, to our knowledge, no previous study had simultaneously assessed multiple epigenetic layers across female and male rodents, with genome-wide data limited to a handful of individual comparisons (H3K4me3^94^, H3K27me3^95^, H3K27ac^95^, and DNA methylation^96,97^). Our results deepen the characterization of baseline sex differences to a broader array of epigenetic layers, and notably uncover large H3K36me3 modifications on the X chromosome in females, which may contribute to XCI.

Beyond physiology, pronounced sex differences are increasingly recognized in psychiatric disorders. This is a much-needed development, given the strong sex ratio observed in most of these conditions, including substance use disorders. Accordingly, sex-specific transcriptomic signatures have been reported in depression^98^, schizophrenia^99^, bipolar disorder^100^, autism^101^, and alcohol use disorder^102^. However, the epigenetic mechanisms that may drive such divergence remained unexplored. To address this gap, we examined how epigenetic landscapes in both sexes respond to chronic exposure to morphine, a potent psychoactive substance. Our results indicated that this reactivity occurred at extremely different loci across females and males for both CG and CAC methylation, as well as histone modifications. These effects were distributed throughout the genome and were not predictable from baseline sex differences observed in saline-treated controls, which - as expected - were mostly located on the X chromosome. Thus, although epigenetic sex differences were present at baseline, those elicited by morphine targeted different genomic loci, and were sex-specific. Despite this divergence, several commonalities documented the robustness of morphine effects. In both sexes, CG- or CAC-DMR exhibited similar distributions along genes, enrichment for similar TF, and organization into clusters of CG methylation and histone profiles that recruited distinct sets of TF. Ultimately, these sex-specific adaptations converged on similar gene modules and biological pathways. Therefore, in a long-term therapeutic perspective, targeting sex-specific loci may increase the efficiency of future epigenetic editing strategies in OUD.

This pronounced sex-dependent epigenetic reprogramming raises important questions about underlying mechanisms. Notably, this divergence occurred in the absence of detectable sex differences in histological features (proportions of D1, D2, or morphine-responsive neurons) or neurophysiological responses (Ca2+ signaling). We therefore propose that it is driven primarily by cell-autonomous processes, whereby female and male epigenomes react to behavioral experiences through shared molecular substrates that nevertheless affect sex-specific loci. This perspective opens new avenues for investigation in at least 2 directions: first, to determine how this principle extends to other psychiatric disorders and possibly to the epigenetic plasticity recruited in physiological contexts; and second, to examine how it is modulated by a combination of gonadal hormone fluctuations^103^, sex-specific chromatin conformation^104^, and sexually dimorphic epigenetic enzymes. Among the latter, the X-linked histone demethylases Kdm5c and Kdm6a - and their Y-linked homologues Uty and Kdm5d - are of particular interest, as they are already attracting attention in the context of neurodevelopmental disorders^105,106^, and may hold equal relevance for molecular psychiatry.

Finally, this study identified NAc GABAergic interneurons as a relatively rare cell population that appears particularly sensitive to morphine-induced multiomic reprogramming. Although only a few reports have implicated this population in cocaine^107,108^ and methamphetamine^109^ use disorders, our results suggest that it also plays an important role in opioid effects and OUD, and reveal the associated epigenetic plasticity. In addition, we highlight new gene candidates that may inform the long-term development of sex-specific therapeutic strategies aimed at alleviating the complex behavioral traits underlying the chronic course of OUD.

## Limitations of the study

In our systematic investigations, we did not detect sex differences in NAc histological organization or in morphine-induced neurophysiological responses. Given the large number of cells analyzed, false negatives are unlikely for the major neuronal populations (D1-, D2-, and MOR-expressing cells). However, changes in rarer cell types may have occurred during morphine treatment and could have contributed to the observed epigenetic sex differences. In the fiber photometry DA experiments, subtle sex differences may also have gone undetected. More refined or alternative methods, such as fast cyclic voltammetry, may prove helpful in future work, as suggested by earlier cocaine studies at lower anatomical resolution (whole striatum)^110,111^. Also, we used similar morphine doses in females and males for direct comparison. Nevertheless, sex differences in the pharmacokinetics and pharmacodynamics of opioids are under investigation^112,113^, and their epigenetic consequences should be investigated. Finally, although our study integrated multiple omic layers, most analyses were limited to bulk tissue. Our integrative analyses indicated that GABAergic interneurons were predominantly targeted in both sexes, suggesting that morphine-induced epigenetic adaptations were sex-specific but converged on common cell populations. Still, it remains possible that part of these adaptations originated from distinct cell types in each sex. Resolving this question will require systematic consideration of sex in future multiomic single-cell studies.

## Resource availability

### Lead contact

Requests for information, resources, and reagents should be directed to Pierre-Eric Lutz.

### Materials availability

Unique materials generated in this study will be available upon request.

### Data and code availability

EM-seq, CUT&Tag, RNA-seq and vTRAP-RNA-seq raw and processed data have been deposited at GEO under accession numbers GSE307783-GSE307786, and will be made publicly available upon publication. Code for most of the bioinformatics and fiber photometry analyses is available on GitHub: https://github.com/pelutzlab/signop. Original microscopy images have been deposited at Mendeley Data (doi:10.17632/c34rng5pgh.1) and will be made publicly available upon publication. Any additional information required to reanalyze the data reported in this paper is available from the lead contact upon request.

## Acknowledgments

We acknowledge help from the In Vitro Imaging Platform - NeuroPôle - Strasbourg (UAR3156), member of the national infrastructure France - BioImaging (https://ror.org/01y7vt929), supported by the French National Research Agency (ANR-24-INBS-0005 FBI BIOGEN). This research was supported by the CNRS (Centre National de la Recherche Scientifique), Strasbourg University (Idex Recherche Exploratoire 2022; PEL), the French National Research Agency [ANR-19-CE37-0010] (PEL), the Fondation Fyssen (Subvention de recherche 2021, PEL), Fondation Avenir (AAP 2022 Recherche médicale Appliquée; PEL), and IReSP-INCa (SPAV1-22-018; PEL). CF was supported by FRM (FDT 202204015236), and the Programme d’Investissement d’Avenir EURIDOL graduate school of pain (ANR-17-EURE-0022). ACR and MD were supported by IReSP-INCa (IReSP-INCa doctoral fellowship 2022-166 and 4th PhD fellowship 2025, respectively). MG was supported by the European Union’s Horizon 2020 research and innovation program under the Marie Sklodowska-Curie grant agreement N°955684.

## Author contributions

CF processed mouse cohorts for RNA-seq and EM-seq analyses, and performed vTRAP and fiber photometry experiments. ACR contributed to all bioinformatic analyses. HS performed Cut&Tag experiments. MG and MY performed CluBCpG analysis. VM established the fiber photometry set-up. MD contributed to qPCR experiments. CD and SM contributed to RNA-sequencing analysis. SO contributed to the preparation of viral vectors. PH performed RNA-Scope experiments. RW contributed to vTRAP experiments. BK and ED generated the MOR-Cre mouse line. AB performed quality controls, alignment, and methylation calling of EM-seq data. KM and ALB contributed to Cut&Tag experiments. IY contributed lab resources. PEL contributed to all bioinformatic analyses and performed LMR/UMR RNA-Scope data analyses. CF, ACR and PEL had access to all raw data and prepared the manuscript, which was approved by all authors.

## STAR Methods

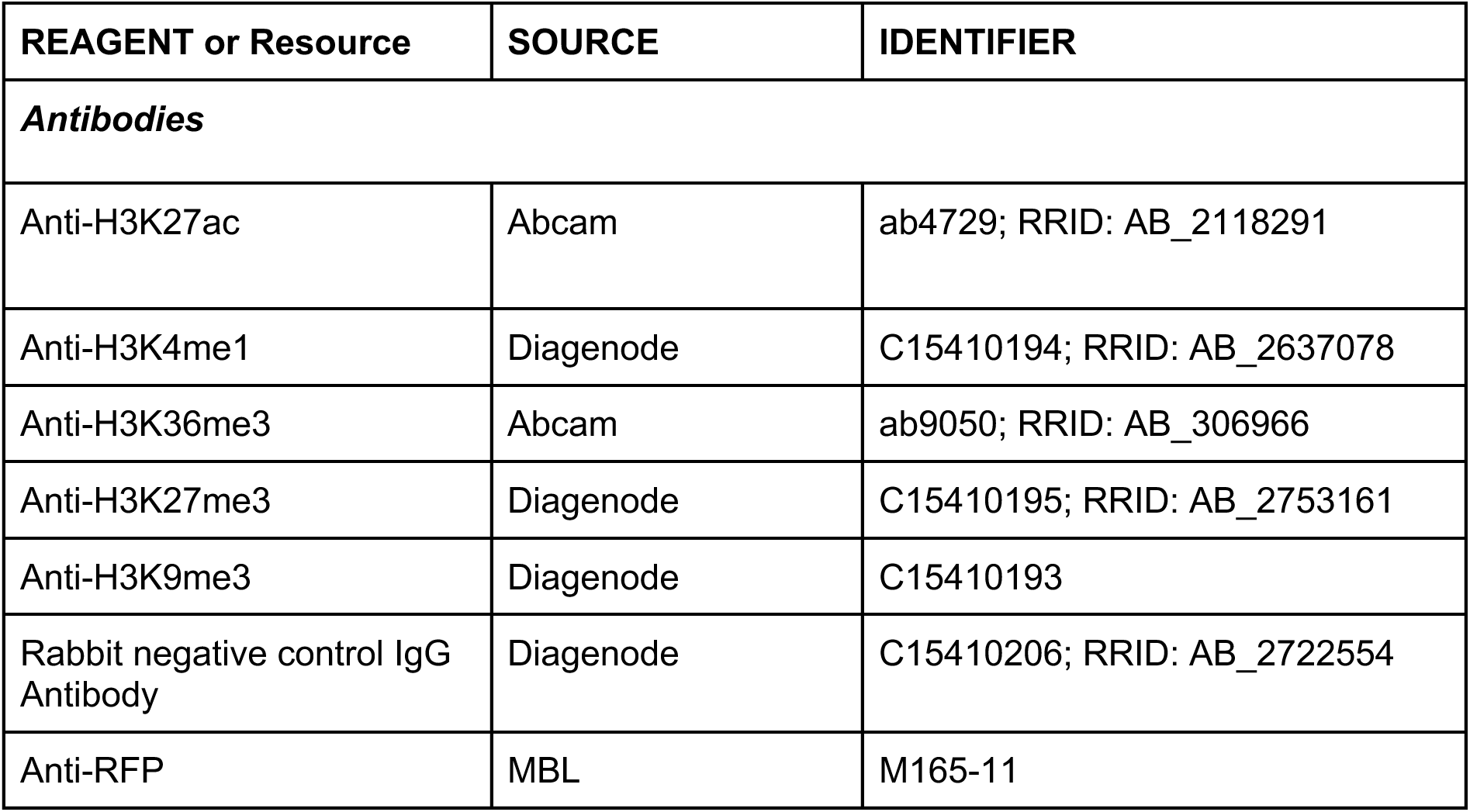
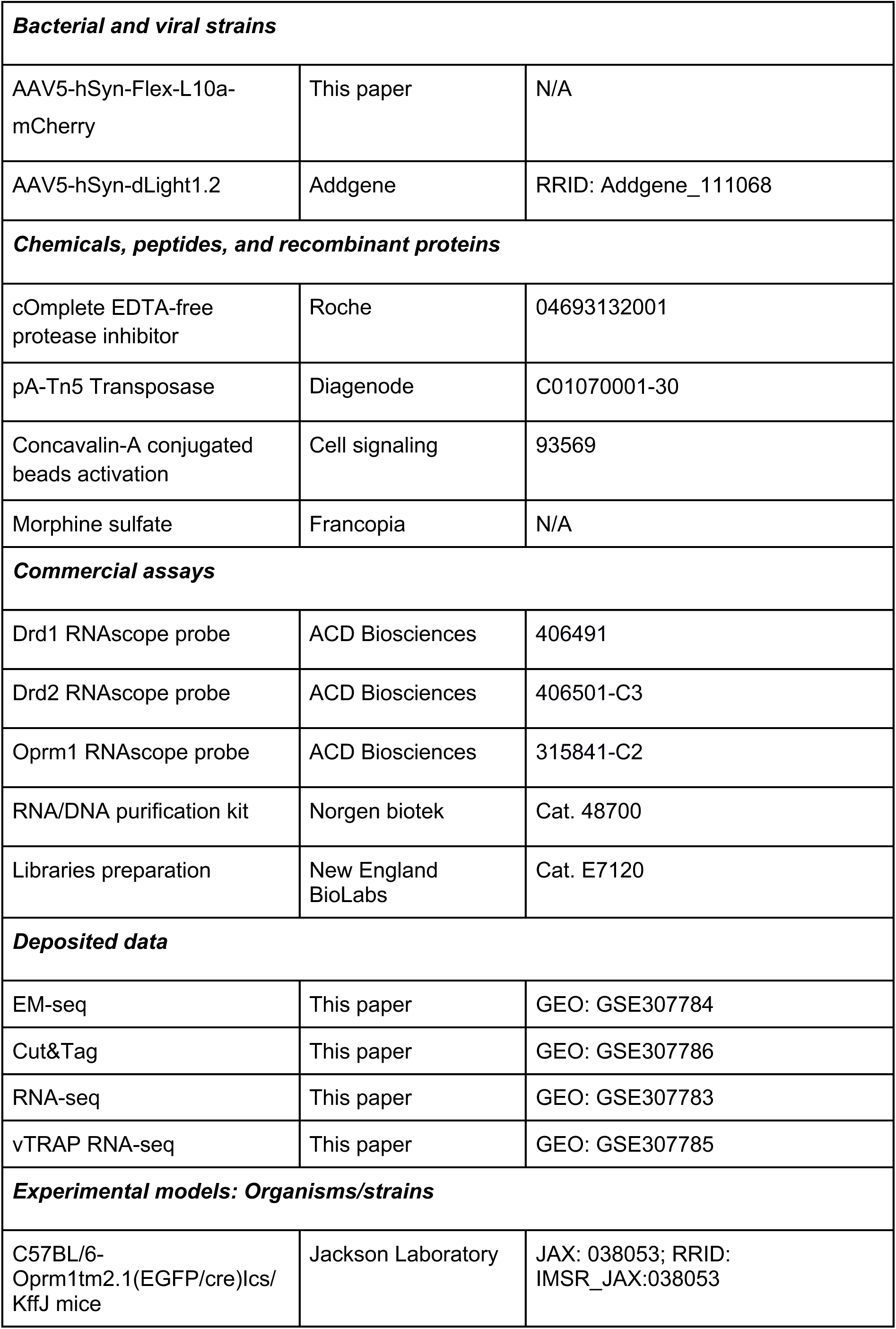
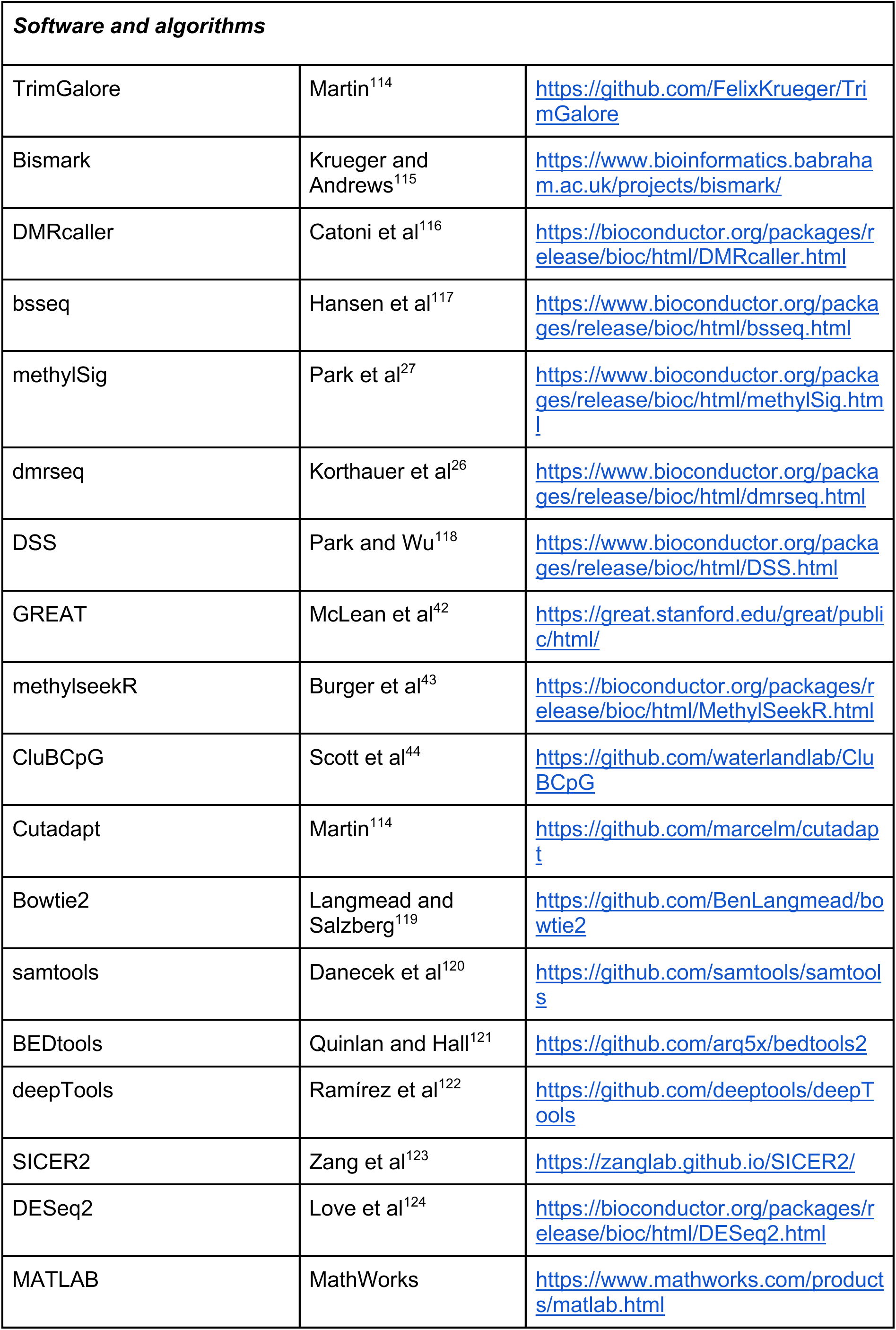
Key Resources Table

### Animals

Experiments were conducted in male and female C57BL/6N mice (Janvier Laboratory, France), 8-week old at the beginning of the experiments. Upon arrival, mice were group housed (3-5 per cage) and maintained on a 12-hr light-dark cycle (lights ON at 7 am) at 24±1°C, humidity around 50%, with *ad libitum* access to food and water. For vTRAP and fiber photometry experiments, we used the MOR-Cre knock-in mouse line, which expresses the Cre recombinase under control of the MOR promoter, on a similar C57BL/6N background^67^. Experiments were conducted at our animal facility (Chronobiotron) registered for animal experimentation (Agreement A67-2018-38), using protocols approved by the local ethical committee of the University of Strasbourg, and performed according to animal care and use guidelines of the European Community Council Directive EU 2010/63.

### Morphine treatment

Male and female mice received twice-daily intraperitoneal (i.p.) injections of morphine sulfate (Francopia, France) at increasing doses (20, 40, 60, 80, 100 mg/kg for 5 days, followed by a single injection on day 6). The control group received saline injections following the same schedule.

### RNA and DNA extraction

Two hours following the last i.p. injection from the treatment described above, animals were sacrificed by cervical dislocation and brains extracted and placed in a matrix (Electron Microscopy Sciences 69022, 30g, Coronal) on ice, then cut into 1-mm coronal slices. Dissections of the NAc were performed on 2 consecutive sections with a circular micro-dissector (1.5-mm diameter), and stored at -80°C. NAc tissue was disrupted and homogenized with a Kinematica Polytron 1600E, and total RNA and DNA co-extracted using the Norgen Biotek RNA/DNA purification kit (Cat. 48700). RNA integrity number (RIN) and concentration were assessed using a 2100 Bioanalyzer (Agilent). DNA was quantified using the Quant-iT™ PicoGreen® dsDNA Assay Kit (Life Technologies).

### EM-seq library preparation

Libraries were prepared using 200 ng of DNA and the NEBNext® Enzymatic Methyl-seq Kit, following the manufacturer’s recommendations (New England BioLabs). Libraries were quantified using the KAPA Library Quantification Complete kit (Kapa Biosystems), and the average size fragment was determined using a LabChip GXII (PerkinElmer) instrument. Sequencing was conducted on an Illumina NovaSeq 6000 (150 bp, paired-end) at an average depth of 204+/-4 million reads/library (mean+/-sem).

### EM-seq alignment, methylation calling, and identification of differentially methylated regions

EM-seq reads were trimmed at their 3’ ends before alignment using Trim Galore (0.6.4, and CutAdapt 2.6), with a >= 20 Phred Score threshold. Bismark (0.22.1)^115^ was used to align the reads against the mm10 reference genome, which was converted to bisulfite mode prior to alignment. Reads were deduplicated before running Bismark’s “methylation_extractor” with –cytosine_report and –CX options, to get methylation status for all cytosine contexts. Over-and under-conversion were assessed using pUC19 (fully methylated) and Lambda (unmethylated) DNA, respectively, resulting in 1 female-morphine sample discarded (Fig.S1). Output files from the methylation extraction step were combined into a single bsseq object for all samples using the R (4.1.0) package bsseq (1.28.0). Spatial correlation was estimated for CG and CAC context using one randomly-chosen female-morphine sample, using the function plotMethylationDataSpatialCorrelation from the DMRcaller (1.26.0) R package. Differentially methylated regions (DMRs) in the CG context were called independently in each sex on collapsed DNA strands by comparing morphine (n=5 or 6) and saline samples (n=6), using 3 R packages: methylSig (1.4.0), dmrseq (1.12.0), and DSS (2.40.0). For dmrseq, candidate regions were considered if they had >= 5% methylation difference. Ten permutations were used to generate the null model, and a p-value threshold of 0.1 was used to define DMRs. For DSS, the p-value threshold at the cytosine level was 0.05, and regions were considered as DMRs if they contained >= 5 CG and had a mean methylation difference >= 5%. For methylSig, we used a window size of 500 bp to match the average length of DMRs identified by the 2 previous approaches. Only windows with a non-null coverage in at least 5 samples per group were considered for DMR-calling, conducted using a 0.01 p-value threshold. DMR in the CAC context were called using the exact same tools and parameters. Simulation of DMR was performed on collapsed CG data using the function “simDMRs” from dmrseq (1.12.0), with the parameter “delta.max0” as effect size. To assess predictive power, we computed the F1-score equal to 2TP/(2TP+FP+FN), where TP, FP, and FN correspond to true positives, false positives, and false negatives, respectively. Gene ontology enrichment of DMR was identified in each cytosine context and sex using GREAT^42^ (4.0.4) and the default “basal plus extension” parameter. Significance of overlaps among sets of genomic ranges was computed using the “overlapPermTest” function of the regioneR package.

### Identification of differences in lowly or unmethylated regions (diffLMRs/UMRs)

We used DNA methylation data in the CG context and the R package MethylSeekR^43^ (1.36.0) to identify fully, lowly and unmethylated regions in every sample (FMRs, LMRs and UMRs, respectively). The default methylation threshold (50%) was used for segmentation, and permutation testing indicated that n=5 consecutive CG sites were required for LMR calling at a FDR=<5%. Following this sample-specific segmentation, we used in-house scripts to identify changes in the localization of UMR/LMR occurring as a function of morphine treatment, independently in each sex. To do so, diffLMR were defined as regions containing at least 10 successive cytosines that individually switched, across saline and morphine groups, from the LMR state to another (FMR/UMR) in at least 4 samples across groups. Similar criteria were used to define diffUMR (as a result, while UMR called in each individual sample contain at least 30 CG sites, diffUMR can be shorter). Gene ontology enrichment of diffLMR or diffUMs was identified using GREAT (4.0.4).

### Identification of morphine-induced changes in single-read methylation patterns (diffBins)

We used the CluBCpG (0.2.5) algorithm^44^, which focuses on the identification of methylation patterns at the level of single reads in 2 steps: first, by identifying reads that fully cover bins of defined length (“clubcpg-coverage”); then, by grouping reads into clusters presenting the same single-read methylation pattern (“clubcpg-cluster”). CluBCpG was run at default parameters for each sample, except that: 1) bin size was set at 150 bp, to match the length of sequencing reads; 2) the “noise filtering” argument was set to FALSE: while CluBCpG by default discards methylation patterns informed by 1 read only, our goal was to measure the frequency of single-read methylation patterns among all available reads, and to then identify changes in frequency induced by chronic morphine. Following the identification of methylation patterns in each biological sample using CluBCpG, we compared their frequency between the morphine and saline groups using in-house scripts, independently in each sex. Focusing on methylation patterns that were reproducibly detected (i.e., non-null coverage) in at least 4/6 samples in each saline/morphine group, frequencies across the 2 groups were compared using a Wilcoxon test, and diffBins were defined by p≤0.01. Gene ontology enrichment of diffBins was identified using GREAT (4.0.4).

### Cut&Tag-Sequencing

Two cohorts of male and female mice received chronic saline or morphine injections. NAc tissue was pooled from 2 mice in cohort1, or from 3 mice in cohort2: each tissue pool from cohort1 was used for the analysis of 2 histone marks (H3K27ac and H3K27me3), while each tissue pool from cohort2 was used for the analysis of 3 other marks (H3K4me1, H3K36me3, H3K9me3). For library preparation, fresh frozen tissue pools (stored at -80°C) were homogenized on dry ice using the KIMBLE Dounce tissue grinder set (Merck). Powder tissue was then resuspended in phosphate-buffered saline solution treated with cOmplete™ Protease Inhibitor Cocktail (PBS-PIC, Roche). Suspension was centrifuged at 1,000 g for 3 min at 4°C, the supernatant removed and the pellet resuspended in 1 mL lysis buffer containing 657.5 µl MilliQ water, 200 µl 50% glycerol, 50 µl 1M HEPES pH 7.5, 50 µl 10% NP-40, 28 µl 5M NaCl, 12.5 µl 20% Triton X-100, 2 µl 0.5M EDTA pH 8.0, and PIC. Samples were incubated at 4°C for 10 min on a rocker and transferred back to the grinder glass tube, homogenized 5 times (stroke “A”), transferred back to 1.5 mL plastic tubes, and pelleted at 1,000 g for 5 min at 4°C. Nuclei pellets were washed again with PBS-PIC, gently resuspended, and filtered using 50 µm pore size CellTrics® filters (Sysmex) to remove remaining pieces of tissue. Here, each sample was divided into equivalent fractions according to the number of studied marks (2 in cohort1, 3 in cohort2). The obtained filtrates were added with Wash Buffer for CUT&Tag to a final volume of 1.45 mL, and kept on ice during following steps. Nuclei were attached to magnetic nanoparticles using Concanavalin A Magnetic (ConA) Beads and Activation Buffer (Cell Signaling)^125^. Briefly, ConA beads with attached nuclei were resuspended and incubated on a rotator at 4°C overnight in 50 µl Antibody buffer, with 1 µl of the undiluted primary antibodies: H3K27Ac (Anti-Histone H3 (acetyl K27) antibody - ChIP Grade, Cat no. ab4729, Abcam), H3K27me3 (H3K27me3 Antibody, Cat no. C15410195, Diagenode), H3K36me3 (Anti-Histone H3 (tri methyl K36) antibody - ChIP Grade, Cat no. ab9050, Abcam), H3K4me1 (H3K4me1 Antibody, Cat no. C15410194, Diagenode), H3K9me3 (H3K9me3 Antibody, Cat no. C15410193, Diagenode), and IgG (Rabbit IgG Antibody - ChIP Grade, Cat no. C15410206, Diagenode). Afterward, the liquid was removed using a magnetic rack, and ConA beads with nuclei resuspended in 100 µl secondary antibody (Guinea Pig anti-Rabbit IgG, Heavy & Light Chain Antibody Preadsorbed, Cat no. ABIN101961, Antibodies online) diluted 100 times in Wash Buffer. Samples were incubated on a rotator at room temperature for 1 hour. After liquid removal and three washes with Wash Buffer using a magnetic rack, recombinant pA-Tn5 protein (pA-Tn5 Transposase loaded, Cat no. C01070001-30, Diagenode) was bound to the current protein-antibody complex (100 µl of 250 times diluted pA-Tn5 in DIG-300). Samples were incubated on a rotator at room temperature for 1 hour. After liquid removal and three washes with DIG-300, a tagmentation reaction was performed with 300 µl of 0.01 M MgCl2 at 37°C for 1 hour. After stopping the tagmentation reaction, nuclei were lysed, and the DNA released in solution at 55°C and extracted in 1.5 mL DNA LoBind tubes using a MinElute PCR Purification Kit (Qiagen) with double elution (15 µl and then 10 µl). Finally, tagged DNA was amplified with a dual indexing system (Nextera XT Index Kit, Illumina) and purified two times with SPRI select (Beckman Coulter): first, with 1.3 volume equivalents of SPRI; second, with 1.4 volume equivalents of SPRI. Concentration and quality of DNA libraries were assessed using Invitrogen Qubit 4 Fluorometer (ThermoFisher Scientific) and the Qubit 1X dsDNA HS Assay Kit, as well as using a BioAnalyzer 2100 (Agilent Technologies) and the Agilent High Sensitivity DNA Kit. Sequencing was conducted on an Illumina NextSeq2000 (50 bp PE) at an average depth of 16.3+/-0.3 million reads/library (mean+/-sem; Fig.S13a).

### Cut&Tag data alignment, peak-calling, and analysis

Data was preprocessed with Cutadapt^114^ (4.0) to trim adapter sequences (Nextera Transposase Sequence) from 3’ end of reads. Cutadapt was used with the following parameters ‘-a CTGTCTCTTATA -A CTGTCTCTTATA -m 25:25’. Reads were mapped to the *Mus musculus* genome (assembly mm10) using Bowtie2^119^ (2.5.0) with default parameters except for ‘--very-sensitive --no-mixed --no-discordant -I 10 -X 700’. Reads whose mapping quality is below 10 were removed using samtools (1.15.1)^126^. Then, reads falling into Encode blacklisted regions v2^127^ were removed using BEDtools intersect (2.30.0)^121^. BigWig files were generated using deepTools bamCoverage^122^ (3.5) with the following parameters ‘-bs 10 -p 10 –normalizeUsing CPM – skipNonCoveredRegions –extendReads’. Peak calling was done on pools of replicates with SICER2 (1.0.3)^123^ with the following parameters: –fragment_size {computed on sample with Homer (4.11)^128^} –gap_size {200, 400, 600, 800, 1000, 2000}. After visual inspection, we selected gap sizes of 200 bp for H3K27ac, 400 for H3K9me3, 800 for H3K27me3, and 1000 for H3K36me3 and H3K4me1. Detected peaks were combined for each comparison (morphine effects in each sex, sex effect in saline-treated groups) to get the union of peaks using Bedtools merge 2.30.0^121^. The number of reads per merged peak and sample was computed using Bedtools intersect. Data were normalized using the method proposed by Anders and Huber^129^ using read counts per peak. Comparisons of groups were performed using the DESeq2 Bioconductor library 1.42.0^130^. To plot and cluster histone mark signal across regions (e.g. DMRs), we used computeMatrix from deepTools, with a binsize equal to 160 (1/20th of the median size of all histone peaks), the display size of the DMRs equal to 8000, and the flanking regions equal to 16000bp. The clustering of the signal was performed using the kmeans function in R, with the optimal number of clusters determined using the fviz_nbclust function from the factoextra package.

### RNA-sequencing library preparation

Starting from 250 ng of total RNA, and following ribosomal RNA depletion using QIAseq FastSelect (Human/Mouse/Rat 96 rxns), cDNA synthesis was achieved with the NEBNext RNA First Strand Synthesis and NEBNext Ultra Directional RNA Second Strand Synthesis Modules (New England BioLabs). The remaining steps of library preparation were conducted using the NEBNext Ultra II DNA Library Prep Kit for Illumina (New England BioLabs). Adapters and PCR primers were purchased from New England BioLabs. Libraries were quantified using the KAPA Library Quantification Complete kit (Universal) (Kapa Biosystems). The average size fragment was determined using a LabChip GXII (PerkinElmer) instrument. Libraries were run on an Illumina NovaSeq 6000 (100 bp PE) at a depth of 64+/-3 million reads/library (mean+/-sem).

### RNA-sequencing alignment, counting and differential expression analysis

STAR V2.7.10b was used for read mapping against the mm10 genome. Quantification of gene expression was performed using HTSeq v0.6.1p1, using gene annotations from Ensembl GRCm38, release 102. Read counts were normalized across libraries with the method proposed by Anders and Huber^130^. Differentially expressed genes were identified using DESeq2, with the Benjamini and Hochberg correction for multiple testing and an adjusted p-value threshold of 0.05.

### Viral Translating Ribosome Affinity Purification (vTRAP)

We used the vTRAP approach developed by Nectow et al^70^, with 2 minor changes: i) as eGFP was already encoded by the MOR-Cre allele, we used mCherry instead for ribosome tagging; ii) expression of the mCherry-tagged Rpl10a ribosomal protein was driven by the hSyn promoter, to capture neurons but not microglial cells that express MOR^131^. We cloned and used an AAV5-hSyn-Flex-L10a-mCherry (1.2e13 genomic copies/mL) encoding the ribosomal L10a protein fused with the mCherry, prepared by triple transfection of HEK293T/17 cells using 3 plasmids: pAAV-EF1α-FLEX-eGFP-RPL10a (Addgene plasmid #98747, modified for mCherry expression under the hSyn promoter, see *Results*); pXR5 (Unc Vector Core), encoding the AAV serotype 5 capsid; and pHelper (Agilent), encoding adenovirus helper functions. After 48 h, AAV2/5 vectors were harvested, treated with Benzonase (Merck) at 100U/mL, purified by Iodixanol gradient ultracentrifugation (OptiprepTM density gradient medium), followed by dialysis and concentration against Dulbecco’s Phosphate Buffered Saline (DPBS) using centrifugal filters (Amicon Ultra-15 Centrifugal Filter Devices 100K, Millipore). Viral titres were quantified by qPCR using the LightCycler480 SYBR Green I Master mix (Roche) and primers targeting the mCherry. Bilateral stereotaxic injections of AAV-hSyn-Flex-L10a-mCherry were performed in the NAc of Oprm1-Cre mice (8-12 weeks), under Zoletil/xylazine anesthesia (50 mg/mL and 2.5 mg/mL, respectively, Centravet). Bupivacaine (2mg/kg) was delivered at the incision site before surgery, and Meloxicam (5mg/kg) at the end of the surgery, and added to the water for the next 3 days. Mice were placed in a stereotaxic frame (Kopf Instruments), and 0.5 µl of virus was injected into the NAc (AP: +1.6mm; L: +/-0.8mm; DV: -4.5mm) using a 5-µl Hamilton syringe (32G; 0.05 µl/min). The needle remained in place for 10 min before suturing. Following surgery, animals were left undisturbed for at least 2 weeks. Two hours following the last morphine or saline injection (similar to bulk RNA-seq experiments), brains were extracted and sliced into 1 mm coronal sections (Electron Microscopy Sciences 69022, 30g), and the NAc dissected on 2 consecutive sections with a circular micro-dissector (diameter 1.5 mm). Bilateral samples from 3 animals were pooled to optimize RNA yield for vTRAP, as previously described^132,133,70^. Tissue was homogenized with a Heidolph RZR 2102 control (900 rpm) in a pre-chilled lysis buffer (20 mM HEPES-KOH pH 7.4, 150 mM KCl, 12 mM MgCl2, 0.5 mM DTT, 100 μg/mL cycloheximide, 10 μl/mL of each RNAse inhibitor (RNasin® 40 U/μL, Promega, and SUPERas-InTM® 20 U/μL, Life Technologies), and cOmplete™ Protease Inhibitor Cocktail (PBS-PIC, Roche)). After centrifugation (2000g, 4°C, 10 min), the polysome-containing supernatant was recovered. Nonylphenyl polyethylene glycol 10% (NP-40) and 1,2-Diheptanoyl-sn-Glycero-3-Phosphocholine 300 mM were added (final concentration 1%) and after 5 min of incubation on ice and centrifugation (20000g, 4°C, 10min), the supernatant was collected, of which 250µL was set aside as “input” (total NAc transcriptome). Polysomes were immunoprecipitated with 200 μL anti-RFP antibody-conjugated beads (M165-11, MBL) pre-washed 3 times in low salt buffer (HEPES KOH [7.3pH], 150mM KCl, 12mM MgCl2, 0.5mM DTT, 100µg/mL cycloheximide) and incubated on a rotator (4°C, 16h). Polysomes were collected on a magnet (Dynamag-2, ThermoFischer Scientific) and washed 4 times with high salt buffer (HEPES KOH [7.3pH], 350mM KCl, 12mM MgCl2, 1% NP-40, 0.5mM DTT, 100µg/mL cycloheximide). Beads were resuspended in nanoprep lysis buffer (20mM HEPES KOH [pH 7.3], 150mM KCL, 12mM MgCl2, 1M DTT, 100µg/mL cycloheximide, protease inhibitors, RNase inhibitors and β-mercaptoethanol), vortexed and incubated (room temperature, 10 min) to release RNA. RNA was extracted by adding an equal volume of 80% sulfolane and following the protocol from the Absolutely Nanoprep kit (Agilent), including DNAse treatment. Enrichment of Oprm1 and mCherry expression in the IP over the Input fraction was assessed before RNA-sequencing. cDNA was synthesized with the M-MLV kit (Thermo Fischer Scientific, Ref: 28025013): 4 μL of oligo-dT (1 μM) and random hexamers (2 μM), 2 μL of dNTPs (10 mM) and 2 μL of H2O were added to each RNA fraction (18 μL). After incubation at 65°C (5 min), 8 μL of 5x First strand buffer (250 mM Tris-HCl pH 8.3, 375 mM KCl, 15 mM MgCl2), 4 μL of DTT (0.1M) and 2 μl of reverse transcriptase (200 U/sample) were added, and samples placed in a thermal cycler: 37°C, 1h15; then 70°C, 15 min (inactivation of reverse transcriptase). For qPCR, a mix of SYBRGreen (10 μL, Applied Biosystems), pairs of primers (final concentration: 500 nM each) and H2O (2 μL) were added to each well containing 6 μL of cDNA (diluted 5 to 10 x after RT). Each sample was run in triplicate on an ABI7300 machine (Applied Biosystems) using: 2 min at 50°C, 10 min at 95°C, 40 amplification cycles (15 s at 95 °C, 1 min at 60°C), followed by a dissociation curve (15 s at 95°, 30 s at 60°C, 15 s at 95°C). Quantification was carried out using the ΔΔCt method, normalized by Actb and Gapdh. Primer sequences: Oprm1 Fwd: CCGAAATGCCAAAATTGTCA, Rvs: GGACCCCTGCCTGTATTTTGT; mCherry Fwd: GAACGGCCACGAGTTCGAGA, Rvs: CTTGGAGCCGTACATGAACTGAGG; Gapdh Fwd: AACGACCCCTTCATTGAC, Rvs: TCCACGACATACTCAGCAC; Actb Fwd: CCTCCCTGGAGAAGAGCTATG, Rvs: TTACGGATGTCAACGTCACAC. For RNA sequencing, library preparation was performed using Clontech SMART-Seq v4 Ultra Low Input RNA and Illumina Nextera XT DNA Library Prep kits. Full-length cDNA was generated from 0.5 ng of total RNA using SMART-SeqX v4 UltraX Low Input RNA Kit for Sequencing (Takara Bio Europe, Saint Germain en Laye, France), according to the manufacturer’s instructions, with 12 cycles of PCR for cDNA amplification by Seq-Amp polymerase. Six hundred pg of pre-amplified cDNA were then used as input for Tn5 transposon tagmentation by the Nextera XT DNA Library Preparation Kit (96 samples) (Illumina, San Diego, USA), followed by 12 cycles of library amplification. Following purification with SPRIselect beads (Beckman-Coulter, Villepinte, France), the size and concentration of libraries were assessed by capillary electrophoresis. Libraries were sequenced on an Illumina HiSeq4000 machine (100bp PE), with image analysis and base calling performed using RTA version 2.7.7 and bcl2fastq version 2.20.0.422.

### Fiber photometry

We used AAV5-hSyn-dLight1.2 (addgene #11106863) and AAV1-hSyn-Flex-GCaMP6m-WPRE-SV40 (addgene #100838^134^) as viral vectors encoding dopamine and calcium sensors, respectively. Surgeries were performed as described above for vTRAP, except that viral injections were unilateral, and cannulas (MFC_400/430-0.66_4.4mm_MF1.25_FLT, Doric) implanted at the end of the surgery, above the injection site (AP: +1.6 mm from bregma, L: + /−0.8 mm, DV: -4,3 mm), and secured to the skull using a screw (diameter: 1.6mm) and Superbond (Universal kit, ref: K058E). Animals were left to rest for at least 3 weeks before experiments. Mice were habituated to handling and to be connected with a patch cord (NA 0.57, Doric Lenses) for 3 days. On test days, mice were connected with the patch cord and placed in a cage for ∼1min before starting any recordings. Each recording was synchronized with a video tracking system (ANY-maze; Aniphy®) using TTLs signals. The fiber photometry setup used 2 light-emitting LEDs: 410-420 nm LED sinusoidally modulated at 333.786 Hz and a 460-490 nm LED sinusoidally modulated at 208.616 Hz (Doric Neuroscience Studio®) merged in an ilFMC6-G2 MiniCube (Doric®) that combines the 2 wavelengths’ excitation light streams and separates them from the emission light. Online real-time demodulation of the fluorescence due to the 410-420 nm and the 460-490 nm excitations was performed by Doric Neuroscience studio software (V6; Doric®). The MiniCube was connected to a fiberoptic rotary joint (Doric®) connected to the cannula. We equalized the 410-420 and 460-490 nm signals, collected at a sampling frequency of 12 kHz, to record an equivalent signal/noise ratio. Custom-generated MATLAB® scripts were used to down-sample and normalize the fluorescence signal. The fluorescence from the control channel (F405, isobestic point) was filtered using a polyfit regression giving a fitted control (F405c). ΔF/F was calculated as (F465 – F405c)/F405c). The deviation of each sample from the averaged signal of a given period was calculated with a Z-score [z=(x-µ)/ σ]. Bins of appropriate time periods were calculated for each saline/morphine injection event and compared between groups.

### Intracardiac perfusion

Animals were anesthetized with Ketamine (300 mg/kg) and Xylazine (20 mg/kg) and perfused with 30 mL of 0.1 M PB (pH 7.4) followed by 100 mL of 4% PFA in 0.1 M PB. After perfusion, brains were removed, postfixed overnight (4% PFA, 0.1M PB), and kept at 4 °C in 0.1 M PBS (pH 7.4) until cutting. Coronal sections (30 µm) were obtained using a vibratome (VT 1000 S, Leica, Deerfield, IL), serially collected in PBS, and mounted with Fluoromount-G (Electron Microscopy Sciences, EM-17984-25). Images were acquired using an epifluorescence microscope (Nikon 80i, Cy3 filter) in order to confirm viral injection sites.

### In-situ hybridization

Two hours after the last morphine injection, brain samples were immersed in isopentane and immediately placed at −80 °C. Frozen samples were embedded in OCT compound, and 14-µm thick sections were cut on a cryostat and mounted on slides. Sections were fixed, dehydrated and pre-treated using the “RNAscope Sample Preparation and Pre-treatment Guide for Fresh Frozen Tissue using RNAscope Fluorescent Multiplex Assay” protocol (Advanced Cell Diagnostics). Hybridization of Drd1 (ACD, 406491), Oprm1 (ACD, 315841-C2), and Drd2 (ACD, 406501-C3) probes and development of the different signals with Opal 520, 590, and 690 fluorophores were performed in accordance with the “RNAscope Multiplex Fluorescent Reagent Kit v2 Assay” instructions (Advanced Cell Diagnostics). Single-layer images were acquired using a slide-scanner NanoZoomer (S60; Hamamatsu Photonics) at ×40 magnification. We used 4 animals per group (i.e. female saline, female morphine, male saline, male morphine) with 4 sections for each animal. Quantifications were done using QuPath 0.3.0 software^135^. First, the region of interest was delimited using the polygon annotation tool. Then, nuclei were detected within regions of interest using the cell detection module on the DAPI staining. To determine *Drd1*, *Drd2*, and *Oprm1* positive cells in regions of interest, object classifiers were trained in QuPath using Random trees classifiers. We selected all the features by output class (Nucleus mean, Nucleus sum, Nucleus standard deviation, Nucleus maximum, Nucleus minimum, Nucleus range, Cell mean, Cell standard deviation, Cell maximum, Cell minimum, Cytoplasm mean, Cytoplasm standard deviation, Cytoplasm maximum, Cytoplasm minimum), and annotated manually a minimum of 20 points for positive cells and negative cells. Classifiers were then applied sequentially on the whole region of interest to determine the *Drd1*, *Drd2*, and *Oprm1* positive cells.

### Gene Set Enrichment Analysis (GSEA)

Genes were ranked based on fold changes obtained during differential expression analysis. GSEA was performed as previously described^136^, using the GSEA Preranked tool.

### Rank-rank hypergeometric overlap analysis

We used either: i) for transcriptomic data and histone modifications, the Rank-Rank Hypergeometric Overlap (RRHO2) procedure^137^), using the R package available at: https://github.com/Caleb-Huo/RRHO2; or ii) for DNA methylation, RedRibbon^138^, a package recently optimized for large datasets that enabled the processing of DNA methylation data. For RRHO2, genes in each data set were ranked based on the following metric: -log10(p-value) x sign(log2 Fold Change). Then, the RRHO2 function was applied to the 2 ranked gene lists at default parameters (with step size equal to the square root of the list length). For RedRibbon, the mm10 genome was partitioned into successive 500-bp genomic bins. Each bin was ranked, in each female or male dataset, based on its p-value and the sign of the methylation difference observed when comparing saline and morphine groups, computed using MethylSig (1.4.0), according to the formula: -log10(p-value) x sign(methylation difference). Then, hypergeometric testing was iteratively conducted on the 2 ranked lists of 500-bp genomic bins (in males and females), to identify p-values. For both RRHO2 and RedRibbon results, the significance of hypergeometric testing is reported as -log10(p-value). Gene ontology enrichment of genes corresponding to the best hypergeometric p-values of each RRHO2 quadrant were determined using the fora function of the fgsea R package (1.20.0).

### Gene network co-expression analysis

We applied multiscale embedded gene coexpression network analysis (MEGENA)^139^. Gene counts were first normalized using counts adjusted according to TMM factors (CTF), and we selected only genes that had more than 1 CPM in 20 out of 24 samples (∼80%), as recommended by a recent benchmark^140^. Gene expression correlations were computed using 10 permutations, and directionality was removed at graph construction. At the clustering step, modules with p-value≤0.01 and comprising between 50 genes and half of the genes in the whole dataset, were retained for annotation. MEGENA also identified subsets of modules (“child modules”) whose genes were more co-expressed than those of their parent module. Multiomic annotation of these modules was then conducted using the following results: whole-tissue RNA-seq differential expression, vTRAP RNA-seq differential expression, CG-and CAC-DMRs, diffLMRs, diffUMRs, diffBins, and DP for the 5 studied histone marks, as well as RNA-seq differential expression in OUD subjects (data from^24^). Overlaps among DEG or genes annotated to genomic sites (e.g. DMR) were assessed using hypergeometric testing and the fora function from the fgsea R package (1.20.0). Following these annotations, their combined significance was computed according to Stouffer’s procedure, using the R package poolr (1.1-1), taking into account all layers (except human data). Annotation of modules to specific cell-types was conducted using gene markers defined in^71^.

