## Supplementary material for "Sex-divergent brain epigenetic reprogramming by chronic opioids"

**Content**

1. Legends of supplementary figures

2. Legends of supplementary tables

### 1. Legends of supplementary figures

**Supplementary Figure 1. Enzymatic Methylation sequencing quality controls and metrics. a.** Enzymatic under-conversion of unmethylated cytosines in CG context, measured using a Lambda DNA spike-in control. **b.** Enzymatic over-conversion of methylated cytosines in CG context, measured using a pUC19 DNA spike-in control. **c.** Number of reads per sample. **d.** Percentage of deduplicated reads. **e.** Percentage of mapped reads. **f.** Whole-genome average DNA methylation levels in non-CG trinucleotide contexts (ANOVA, effect of trinucleotides context p<2e-16; effect of sex p=0.01). **g.** Whole-genome average DNA methylation in the CG context (ANOVA, effect of sex p=0.09, effect of morphine treatment p=0.97). **h.** Whole-genome average DNA methylation in the CAC context (ANOVA, effect of sex p=0.03, effect of treatment p=0.25).

**Supplementary Figure 2. Convergence among DMR calling methods.** We used 3 algorithms for the identification of Differentially Methylated Regions (DMR): i) MethylSig, a method that compares methylation levels in non-overlapping genomic bins of fixed length, irrespective of the positions of cytosines; ii) DSS, which compares methylation levels of individual CG sites across biological groups, and then identifies DMR by joining successive sites meeting a significance threshold; iii) dmrseq, which uses a weighted smoothing approach to correct methylation levels according to their spatial correlation and coverage, before calling DMRs using permutations. **a.** CG-DMR identified in males. **b-c.** CAC-DMR identified in females (b) and males (c). DSS identified 7775 CAC-DMRs in females, 8259 in males; Methylsig, 15878 in females, 17491 in males; dmrseq, 681 in females, 569 in males. Of note, dmrseq identified much fewer CAC- than CG-DMRs, likely reflecting its reliance on smoothing and spatial correlation of DNA methylation in the CG, but not other, cytosine contexts (see Fig.S5). **d.** Density plot of ranks of DMR identified in common by pairs of DMR callers ($\chi^2$: Female CG: DSS vs dmrseq, p=4e-10; methylsig vs dmrseq, p=4e-4; DSS vs methylsig p=4e-10; Male CG, DSS vs dmrseq, p=7e-7; methylsig vs dmrseq p=2e-4; DSS vs methylsig, p=6e-5; Female CAC, DSS vs dmrseq, p=N/A; methylsig vs dmrseq, p=0.82; DSS vs methylsig, p=3e-1; Male CAC, DSS vs dmrseq, p=1; methylsig vs dmrseq, p=1; DSS vs methylsig, p=5e-19).

**Supplementary Figure 3. Convergence of results generated by DSS, dmrseq and MethylSig for simulated DMR.** DMRs were simulated at increasing effect size from 5 to 80%. **a.** Jaccard indices (JI) for DMRs called by 2/3 methods on simulated DMRs. JI reached a plateau around 33%, even for DNA methylation group differences up to 80%, indicating a modest convergence across the 3 tools we used, even for large differences in DNA methylation (Fig.S3). **b-d.** Number of false positives (**b**), false negatives (**c**) and true positives (**d**) called by each method on simulated DMR. **e-f.** Precision (**e**) and sensitivity (**f**) for each method on simulated DMR. **g.** F1-scores (harmonic mean of precision and sensitivity) of the DMR callers on simulated DMR.

**Supplementary Figure 4. Comparison of DMR identified by a single as opposed to at least 2 DMR callers. a.** Length of female CG-DMR found by 1 tool versus ⩾ 2 tools (1/3 = 498+/-1.42; ⩾2/3 = 683+/- 4.77; p=1e-268). **b.** Number of CGs at female CG-DMR found by 1 tool versus ⩾ 2 tools (1/3 = 6.99+/-0.05; ⩾2/3 = 10.66 +/- 0.09; p=8e-229). **c.** Absolute mean difference in DNA methylation at female CG-DMR found by 1 tool versus ⩾ 2 tools (1/3 = 0.1+/-0.06; ⩾2/3 = 0.11+/-0.03; p=3e-37). **d.** Length of male CG-DMR found by 1 tool versus ⩾ 2 tools (1/3 = 492+/-1.25; ⩾2/3 = 690+/-5.28; p=1e-255). **e.** Number of CGs at male CG-DMR found by 1 tool versus ⩾ 2 tools (1/3 = 6.47+/-0.04; ⩾2/3 = 10.36+/-0.09; p=7e-283). **f.** Absolute mean difference in DNA methylation at male CG-DMR found by 1 tool versus ⩾ 2 tools (1/3 = 0.1+/-0.06; ⩾2/3 = 0.11+/-0.03; p=5e-42). **g.** Length of female CAC-DMR found by 1 tool versus ⩾ 2 tools (1/3 = 427+/-1.16; ⩾2/3 = 537+/-2.23; p=7e-285). **h.** Number of CACs at female CAC-DMR found by 1 tool versus ⩾ 2 tools (1/3 = 16.30+/-0.07; ⩾2/3 = 19.61+/-0.30; p=1e-25). **i.** Absolute mean difference in DNA methylation at female CAC-DMR found by 1 tool versus ⩾ 2 tools (1/3 = 0.05+/-0.02; ⩾2/3 = 0.06 +/-0.02; p=3e-30). **j.** Length of male CAC-DMR found by 1 tool versus ⩾ 2 tools (1/3 = 424+/-1.17; ⩾2/3 = 534+/-1.57; P-value=0). **k.** Number of CACs in male CAC-DMR found by 1 tool versus ⩾ 2 tools (1/3 = 16.15+/-0.07; ⩾2/3 = 18.55+/-0.2; p=6.69e-30). **l.** Absolute mean difference in DNA methylation at male CAC-DMR found by 1 tool versus ⩾ 2 tools (1/3 = 0.05+/-0.03; ⩾2/3 = 0.06+/-0.02; p=5.79e-42). Results are expressed as mean+/-sem.

**Supplementary Figure 5. Metrics and genomic distribution of DMR. a.** Correlation of DNA methylation levels at neighboring cytosines in CG and CAC contexts as a function of distance between them. **b.** Principal component analysis of female CG-DMR. **c.** Principal component analysis of male CG-DMR. **d.** Principal component analysis of female CAC-DMR. **e.** Principal component analysis of male CAC-DMR. **f.** Distance in base pairs between DMR and their associated genes, identified using GREAT (2-way ANOVA; sex effect: [F = 3.39; p=0.07]; context effect: [F=506; p<2e-16]; interaction sex x context [F =2.81; p=0.09]). **g.** Male DMR localization with respect to genes.

**Supplementary Figure 6. Comparison of sex- and morphine-related DMR. a-b.** Mean differences in DNA methylation levels *observed* in females or males at loci corresponding to morphine-induced DMR initially *identified* in each sex. **c.** Permutation testing of distances between nearest female & male DMR in the CG context. **d.** Permutation testing of distances between nearest female & male DMR in the CAC context. **e.** Overlap of female or male morphine-induced DMR with sex-DMR (identified when comparing male saline-treated and female saline-treated groups), in the CG context. **f.** Overlap of female or male morphine-induced DMR with sex-DMR, in the CAC context. **g.** Localization on the X chromosome and autosomes of female and male morphine-DMR, and of sex-DMR, in each cytosine context. **h.** Distribution of DMR into subgroups where decreased (hypo-DMR) or increased (hyper-DMR) DNA methylation levels were observed as a function of morphine, or sex, in each cytosine context. For sex-DMRs, hyper-DMRs correspond to higher DNA methylation levels in females.

**Supplementary Figure 7. Transcription factor (TF) binding motif enrichment analyses.** **a.** Correlation of TF binding motifs enriched in CG- or CAC-DMR in females (p=0.004, r=-0.099). **b.** Correlation of TF binding motifs enriched in CG- or CAC-DMR in males (p=4.91e-09; r=-0.2). **c.** List of the 10 most significantly enriched TF in each cytosine context and sex. As observed for CG-DMR (see main text), 4 of the TF enriched in CAC-DMR were common to males and females, with none among the top 10 in CG-DMR. JASPAR Matrix IDs of TF: ARNT_HIF1A, MA0259.1; AHR_ARNT, MA0006.1; GMEB1, MA0615.1; HIF1A, MA1106.1; ATF1, MA0604.1; TFDP1, MA1122.1; E2F6, MA0471.2; ZIC1_ZIC2, MA1628.1; ZBTB14, MA1650.1; ARNT, MA0004.1; ZNF263, MA0528.2; FOXN1, MA1684.1; ZIC3, MA0697.2; KLF9, MA1107.2; EGR1, MA0162.4; PAX4, MA0068.2; DLX2, MA0885.2; DLX5, MA1476.2; DRGX, MA1481.1; EOMES, MA0800.1; TBX3, MA1566.2; LIN54, MA0619.1; MGA, MA0801.1; BARX2, MA1471.1; ALX4, MA0853.1; LHX1, MA1518.2; HOXB9, MA1503.1; ARID3B, MA0601.1; ALX1.MA0854.1. **d-e.** Mean differences in DNA methylation levels *observed* in the CG or CAC context at loci corresponding to morphine-induced DMR initially *identified* in each CG (d) or CAC (e) cytosine context (NB: results for DMR identified initially are similar to those in Fig.S6).

**Supplementary Figure 8. Characteristics of lowly methylated regions (LMR), unmethylated regions (UMR), and morphine-induced changes affecting these features.** **a.** Number of LMR and UMR across groups. **b.** Percentage of methylation of LMR and UMR across groups. **c.** Size (in base pairs) of LMR and UMR across groups. **d-e.** Repartition of UMR (d) or LMR (e) in the genome with respect to gene features (for 1 representative sample). **f.** Jaccard Indices of pairwise overlaps of LMR or UMR among samples. **g-h.** Absolute values of DNA methylation differences observed in males (g) or females (h) at morphine-induced CG-DMR, diffLMR and diffUMR. **i-k**. Principal component analysis of female diffUMR (i), male diffUMR (j), and female diffLMR (k). **l-m.** Venn diagrams of diffUMR in males and females (l), and of diffLMR in males and females (m).

**Supplementary Figure 9. Single-read DNA methylation pattern analysis. a.** Relationship between the number of bins and the number of CG sites per bins observed when considering 100bp or 150bp bins using ClubCpG. **b.** Morphine-induced diffBins results showing the frequencies of their DNA methylation patterns across saline- and morphine-treated groups, in males**. c-d.** Principal component analysis of female (c) and male (d) diffBins. **e-f.** Genomic overlaps among diffUMR, diffLMR, CAC-DMR, CG-DMR in males (e) or females (f). **g.** Comparative distributions with respect to genes of male and female CG-DMR, CAC-DMR, diffLMR, diffUMR and diffBins. **h.** Venn diagram of male and female ClubCpG diffBins. LMR: Lowly methylated region; UMR, Unmethylated region.

**Supplementary Figure 10. Threshold-free comparisons across sexes of morphine-induced DNA methylation changes. a.** Venn diagram of the best hypergeometric p-value from the up/up quadrant of the male/female Redribbon comparison, in the CG context (Fig.2f). **b.** Venn diagram of the best hypergeometric p-value from the down/down quadrant of the male/female Redribbon comparison, in the CG context (Fig.2f). **c.** Venn diagram of the best hypergeometric p-value from the up/up quadrant of the male/female Redribbon comparison, in the CAC context (Fig.2g). **d.** Venn diagram of the best hypergeometric p-value from the down/down quadrant of the male/female Redribbon comparison, in the CAC context (Fig.2g). **e.** Mean morphine-induced DNA methylation differences observed: (i) in the CG context, at genomic bins common to both sexes identified from the male/female RedRibbon comparison (down-down quadrant), as well as hypo-DMR identified in males or females (top-left panel) ; (ii) in the CAC context, at genomic bins common to both sexes identified from the male/female RedRibbon comparison (up-up quadrant), as well as hyper-DMR identified in males or females (top-right panel) ; (iii) in the CAC context, at genomic bins common to both sexes identified from the male/female RedRibbon comparison (down-down quadrant), as well as hypo-DMR identified in males or females (bottom-left panel). **f.** Permutation testing of distances between sex-specific morphine-induced DMR identified in males and sex-convergent 500-bp genomic bins identified using RedRibbon, also in males. **g.** Permutation testing of distances between sex-specific morphine-induced DMR identified in females and sex-convergent 500-bp genomic bins identified using RedRibbon, also in females.

**Supplementary Figure 11. Functional convergence of morphine-induced methylomic changes. a.** Convergence observed across the 2 sexes for morphine-DMR in terms of genomic position, genes associated to those positions, and GO terms associated with those genes. Similar results were obtained in the 2 cytosine contexts. **b.** Description of the method used to determine whether genes associated with DMR in only one sex were enriched in similar biological functions as those shared across sexes (see main text for details). **c.** Convergence observed across the 2 cytosine contexts for morphine-DMR in terms of genomic position, genes associated to those positions, and GO terms associated with those genes. Similar results were obtained in the 2 sexes. **d.** Intersection among coordinates (upper), genes (middle) and biological pathways (bottom) associated with DMR identified across 2 sexes and 2 cytosine contexts: female CG-DMR, female CAC-DMR, male CG-DMR and male CAC-DMR. Regarding genomic coordinates, the vast majority (99.3%) of DMR were unique to each set, with only 0.7% (N=97) corresponding to intersections among 2 of them, and none to intersections among more than 2. For DMR-associated genes, the number of genes unique to a single set dropped to 51.9% (N=4669), while 26.8% (N=2405), 14.3 (N=1289) and 7% (N=625) were common to 2, 3 or 4 sets. Lastly, for DMR-associated GO terms, the number of terms enriched for 2, 3 or 4 sets increased to 22.3% (N=490), 14% (N=307) and 27.5% (N=603), respectively. Most enriched GO terms common to all 4 sets were related to general processes such as system development or cell differentiation (Supplementary Table 2).

**Supplementary Figure 12. Convergence across methylomic features. a.** Convergence at the level of genomic coordinates, genes and GO terms of morphine-induced methylomic changes observed in females across 5 types of features: CG-DMR, CAC-DMR, diffUMR, diffLMR and diffBins. **b.** Convergence in males across the same 5 features.

**Supplementary Figure 13. Analysis of 5 histone marks. a.** H3K27ac, H3K27me3, H3K36me3, H3K4me1 and H3K9me3 were analyzed by Cut&Tag sequencing, with IgG controls generated in each sex and for each batch of library preparation (see *Methods*). The table shows the mean sequencing depth achieved, and number of peaks identified, per biological replicate in females (F) or males (M). **b-c.** Correlation-based clustering of the 5 histone marks for every sample, when normalizing by sex-specific IgG controls only (b), or by batch-specific IgG controls only (c). In comparison, normalizing by both IgG controls (sex and batch) led to a better clustering (see main text and Fig.3a). **d.** Identification of differential peaks (DP) using various p-value thresholds. **e.** Number of DP for the 3 comparisons (morphine-induced differences in males or in females, sex-differences in saline controls), and 5 marks, using the $p<0.01$ significance threshold (see main text). **f.** Fold-changes (log2-transformed) at DP identified in the X chromosome when comparing females and males (saline-treated controls), for the 3 marks showing large sex differences (H3K27me3, H3K36me3 and H3K9me3).

**Supplementary Figure 14. a-e.** Venn diagrams of overlaps among differential peaks (DP) identified as a function of morphine in each sex (morphine-DP), or as a function of sex (sex-

DP, comparing saline-treated groups), for H3K27ac (a), H3K27me3 (b), H3K36me3 (c), H3K4me1 (d) and H3K9me3 (e). **f.** Table of Jaccard indices of overlaps among sex and morphine-effects, in each sex, for each of the 5 marks.

**Supplementary Figure 15. a-c.** RRHO2 comparisons of morphine-induced changes in abundance of H3K27me3 (a), H3K36me3 (b) and H3K9me3 (c) observed across females (x-axis) and males (y-axis; considering regions identified as peaks for each mark). Significant enrichments in concordant directions (bottom-left and upper-right quadrants) were detected for the 3 marks (H3K36me3, p=3.2e-45; H3K9me3, p=5.0e-9; H3K27me3, p=2.5e-6), with a discordance for H3K27me3 only (p=7.9e-14). **d.** Odds-ratios and p-values for the permutation analyses of the functional convergence at the GO level. **e.** Correlation among parent GO terms enriched for morphine-induced changes in DNA methylation (x-axis) and histone marks (y-axis), ranked according to the number of individual GO terms significantly enriched, aggregated using the rrvgo R package. **f-h.** Profiles of histone marks across the 3 clusters of male CG-DMRs (as identified in Fig.1) based on the patterns of said histone marks in saline (opaque line) and morphine (translucent line) samples, with a flanking region +/-16kb. **i.** Comparison of transcription factor binding motifs enrichment in Cluster2 female CG-DMR and the full set of female CG-DMR. Note that some of the TF binding motifs enriched in Cluster2 DMR were not enriched in the full set of DMR. Similar results were obtained in males (data not shown). **j.** Morphine-induced DNA methylation differences observed in each sex at male cluster2-CG-DMRs. Similar results were found for cluster 3 (data not shown). **k-l.** Profiles of histone marks observed in each sex at cluster2 CG-DMR identified initially in females (k) or in males (l). Similar results were obtained for cluster3 DMR (data not shown).

**Supplementary Figure 16. In-situ hybridization experiments. a.** Experimental design (see also *Methods*). **b.** Proportion of cells per class. **c.** Frequency of mu opioid receptor-expressing (MOR+) medium spiny neurones (MSNs) across the 4 slices (ANOVA, effect of section [F(3,53)=1.97; p=0.13], effect of MOR+ MSN subtype [F(1.975,104.7)=209.9; p<1e-4], interaction [F(6,106)=2.76; p=0.01], post hoc: MOR+Drd1+Drd2+ section 1 vs section 4 p=0.02; MOR+Drd1+Drd2+ section 2 vs section 4 p=0.003). **d.** Comparison of MSN subtypes across the general MSN population and the MOR+ population in females and males (2-way ANOVA, female MOR expression effects [F(1,14)=0; p>0.99]; MSN subtypes effect [F(2,28)=160.2; p<1e-10]; interaction [F(2,28)=3.41; p=0,04]; post-hoc: Drd1+Drd2+ in all MSNs versus MOR+ MSNs p=0,01; male MOR expression effects [F(1,14)=0.36 ; p=0.55]; MSN subpopulation effect [F(2,28)=122.4; p<1e-10]; interaction [F(2,28)=3.667; p=0.04]; post-hoc: Drd2+ in all MSNs versus MOR+ MSNs p=0.02; Drd1+Drd2+ in all MSNs versus MOR+ MSNs p=0.03. **e.** MOR expression (z-score) across sexes and treatment groups (3-way

ANOVA, effect of MSN subtype [$F(2,24)=55.84$; $p<1e-4$]). **f.** Representative image of hSyn-FLEX-GCamp virus expression in the nucleus accumbens. *, $p <0.05$.

**Supplementary Figure 17. RNA sequencing quality control and results. a.** Number of reads per sample across groups. **b.** Percentage of uniquely mapped reads across groups. **c.** Volcano plots of morphine-induced differentially expressed genes (DEG) between saline and morphine-treated mice in female or male mice. **d**. GO enrichment analysis of genes implicated in the best RRHO2 p-value when comparing male and female bulk tissue morphine-induced transcriptomic adaptations (Fig.5c). The top 10 GO terms associated with upregulated and downregulated genes are shown. **e.** Enrichment score (Gene Set Enrichment Analysis, GSEA; see *Methods*) of female RNAseq results for the "opioid signaling" reactome gene set ($p=1$, NES=0.40). **f.** Enrichment score of male RNAseq results for the same "opioid signaling" reactome gene set ($p=1$, NES=0.50). **g.** RRHO2 comparisons of gene expression changes associated with OUD in men and women combined (x-axis) and female and male mice combined (y-axis).

**Supplementary Figure 18. Viral translating ribosome affinity purification (vTRAP) experimental design and quality controls. a**. Experimental design for vTRAP experiments. The vTRAP virus (AAV-hSyn-Flex-L10a-mCherry) was injected in the nucleus accumbens of MOR-Cre mice by stereotaxic surgery, in order to tag ribosomes from MOR+ neurons. A representative image of the expression pattern of the vTRAP virus is shown. **b.** Experimental design of quality controls for the quantification of MOR and mCherry expression in the immunoprecipitated (IP) RNA fraction as opposed to the Input RNA fraction (corresponding to bulk tissue). **c.** qPCR results for MOR and mCherry enrichment in IP RNA fractions. No differences were observed across sexes or treatment groups. **d**. Percentage of duplicated reads across groups. **e**. Percentage of uniquely mapped reads across groups.

**Supplementary Figure 19. Viral translating ribosome affinity purification (vTRAP) results. a-b**. Volcano plot representing differentially expressed genes (DEG) as a function of morphine in males (a) or females (b). **c**. Venn diagram of overlap among female and male vTRAP DEG. **d**. RRHO2 of male vTRAP (x-axis) versus female vTRAP (y-axis) transcriptional effects of chronic morphine treatment. **e-f**. RRHO2 of vTRAP (y-axis) versus bulk tissue (x-axis) transcriptional effects of morphine in females (e) or males (f). **g**. Box plot of p-values associated with the 100 most enriched GO terms in RRHO2 male vs female down-down quadrant comparing vTRAP and whole tissue results ($p=3e-26$). **h-i**. GSEA enrichment scores of females (h) and males (i) vTRAP results for the "opioid signaling" reactome gene set ($p=0.11$, NES=-1.21).

**Supplementary Figure 20. a.** Odds-ratio from permutation testing among genes associated with differences observed for each omic layer, in males. **b.** Correlation of odds-ratios from permutation testing conducted using male and female data (p=4e-50, r=0.93). **c.** Aggregated parent GO terms generated using rrvgo (obtained using the Stouffer procedure to summarize enrichment across all omic layers for each GO term). **d.** Correlations among enrichments of modules for pairs of omic layers. Strongest correlations were observed between DMR and diffBins, followed by correlations between methylomic features and histone marks (see also below). Lower, but still significant, correlations were detected for omic features less frequently affected by morphine (diffLMR, diffUMR, H3K36me3), overall supporting the validity of the complementary strategies employed alongside DMR calling. **e.** Correlation of module rankings based on epigenetic features (y-axis) and transcriptomic (x-axis) features (p=0.02, r=0.56), in females. Interestingly, out of the 16 parent modules composing our network, M3 and M8 were ranked 2nd and 6th for enrichment in morphine-induced transcriptomic changes, respectively, but only 12th and 15th for epigenetic modifications. M3 was also significantly associated with endothelial cells. It is possible to speculate that dysregulations affecting these 2 modules may preferentially reflect acute effects of the last morphine injection, which may rely less on the epigenetic plasticity that is progressively recruited upon repeated injections. Future work will be necessary to test this hypothesis. **f.** Correlation of module rankings based on histone-related features and methylation-related features (p=3.9e-8, r=0.94), in females.

### 2. Legends of supplementary tables

**Supplementary Table 1.** Coordinates, DNA methylation difference and localization with respect to genes of consensus Differentially methylated regions (DMR) identified as a function of chronic morphine treatment (in each sex), or as a function of sex (comparing male and female saline-treated controls).

**Supplementary Table 2.** Coordinates and localization with respect to genes of switches in UMR/LMR status (diffUMR/LMR), or changes in single-read DNA methylation patterns, identified as a function of morphine in each sex, using methylseekR or CluBCpG, respectively (see *Methods*).

**Supplementary Table 3.** Gene Ontology (GO) terms significantly associated in each sex with morphine-induced differences observed for each methylation feature (CG-DMR, CAC-DMR, diffUMR, diffLMR, diffBins), as well as reduced GO terms (obtained using rrvgo).

**Supplementary Table 4.** Genes (or GO terms) associated at least once with a methylomic feature, and the number of features they were associated with (using GREAT).

**Supplementary Table 5.** Coordinates, p-value, fold-change and localization with respect to genes of differential peaks (DP) identified, for each histone modification, as a function of chronic morphine treatment (in each sex), or as a function of sex (comparing male and female saline-treated controls).

**Supplementary Table 6.** Gene Ontology (GO) terms significantly associated in each sex with chronic morphine-induced differential peaks (DP) for each histone modification, as well as reduced GO terms (obtained using rrvgo).

**Supplementary Table 7.** Differential expression analysis (fold-changes, nominal p-values and Benjamini-Hochberg adjusted p-values) of RNA-sequencing data generated in bulk-tissue, or focusing on mu opioid receptor-expressing neurons. FS, female mice treated with saline injections; FM, female mice treated with morphine injections; MS, male mice treated with saline injections; MM, male mice treated with morphine injections.

**Supplementary Table 8.** Gene Ontology (GO) terms associated with differentially expressed genes (in each sex) in bulk-tissue or MOR-expressing neurons, as well as reduced GO terms (obtained using rrvgo).

**Supplementary Table 9.** *Mus musculus* genes corresponding to best p-values of each quadrant of the RRHO2 comparing results from our bulk tissue differential expression analysis in the mouse with results from the differential expression analysis conducted in humans with OUD by Seney et al[1], in each sex.

**Supplementary Table 10.** Genes differentially expressed both in bulk-tissue and by MOR-expressing neurons, in each sex (sheet 1). Genes corresponding to the best p-value of each quadrant of the RRHO2 comparing our bulk-tissue and MOR-expressing neuron-specific differential expression results, in each sex (sheets 2 and 3).

**Supplementary Table 11.** Combined p-values (using the Stouffer method) of Gene Ontology (GO) terms associated with 12 omic features (methylation, histone modifications, transcriptome), as well as reduced GO terms (obtained using rrvgo).

**Supplementary Table 12.** Module gene composition (sheet 1), p-values of significance of association to the 12 omic layers in females (sheet 2) or males (sheet 3), p-values of the significance of the association to cell types (using marker genes from[2]; sheet 4).

**Supplementary Table 13.** Gene Ontology (GO) terms associated with 5 modules most affected by morphine, as well as reduced GO terms (obtained using rrvgo).
