## Supplementary figures and images for "Sex-divergent brain epigenetic reprogramming by chronic opioids"

### Supplementary Figure 1

**Supplementary Figure 1**

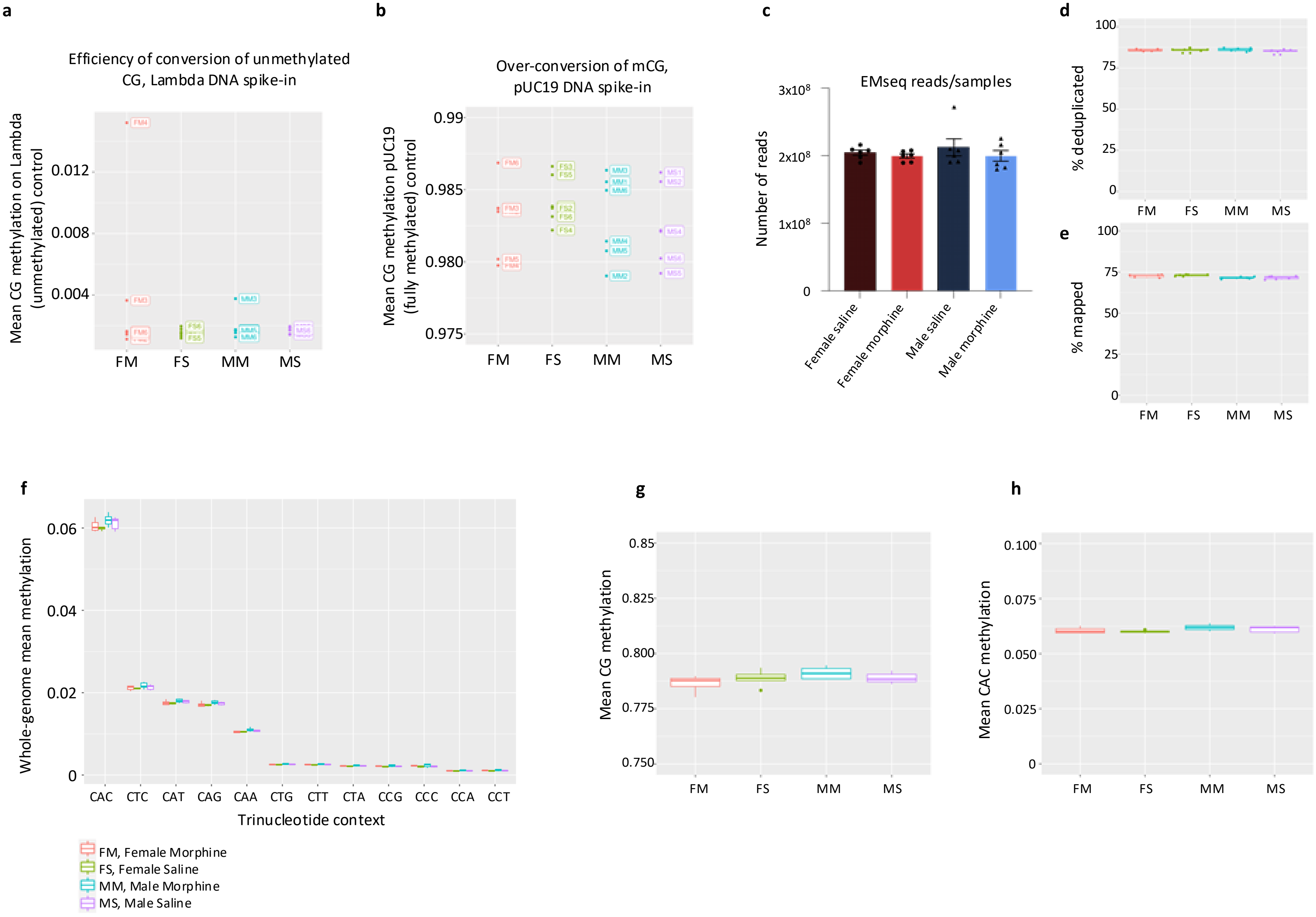

### Supplementary Figure 2

##### Supplementary Figure 2

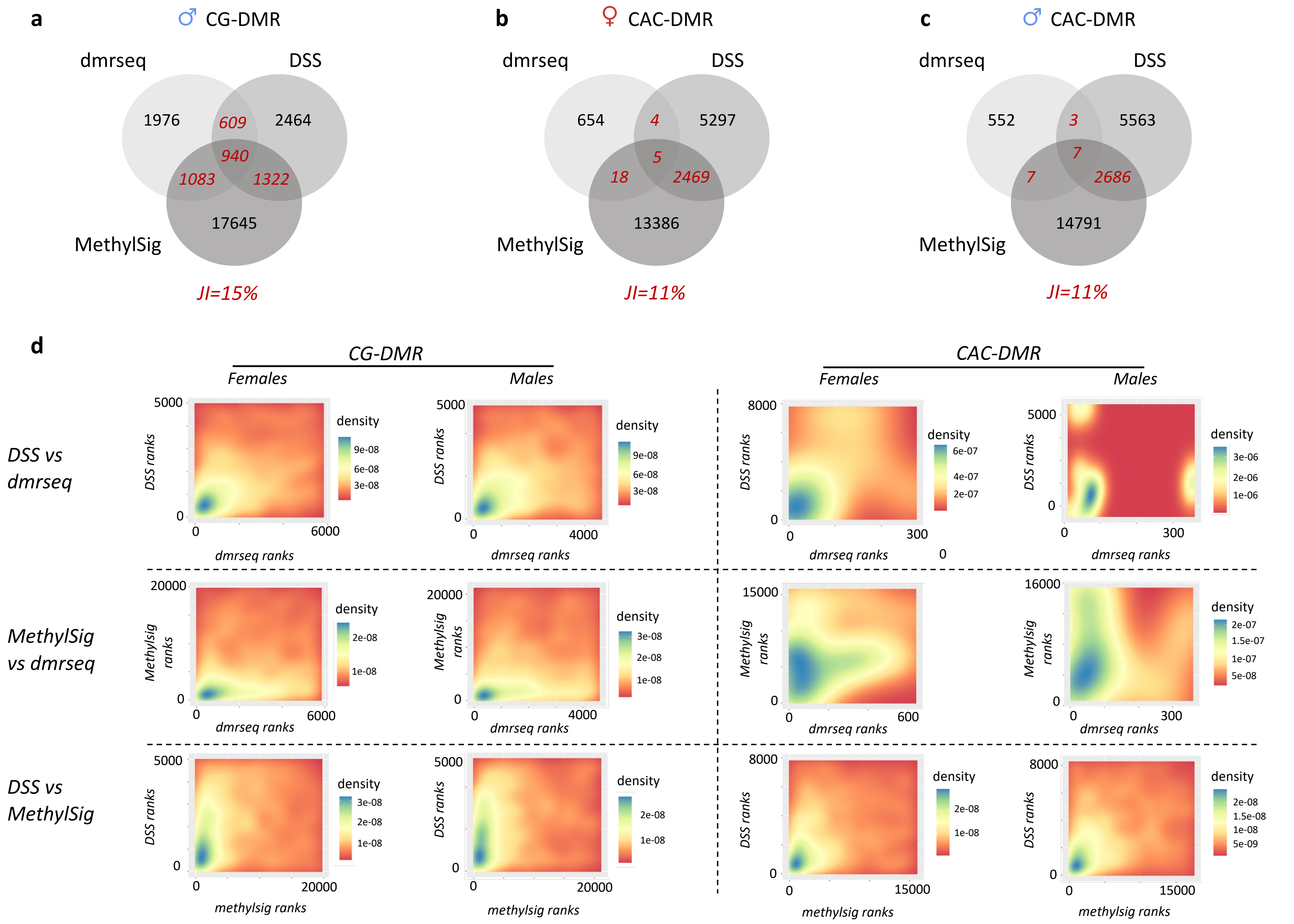

### Supplementary Figure 3

**Supplementary Figure 3**

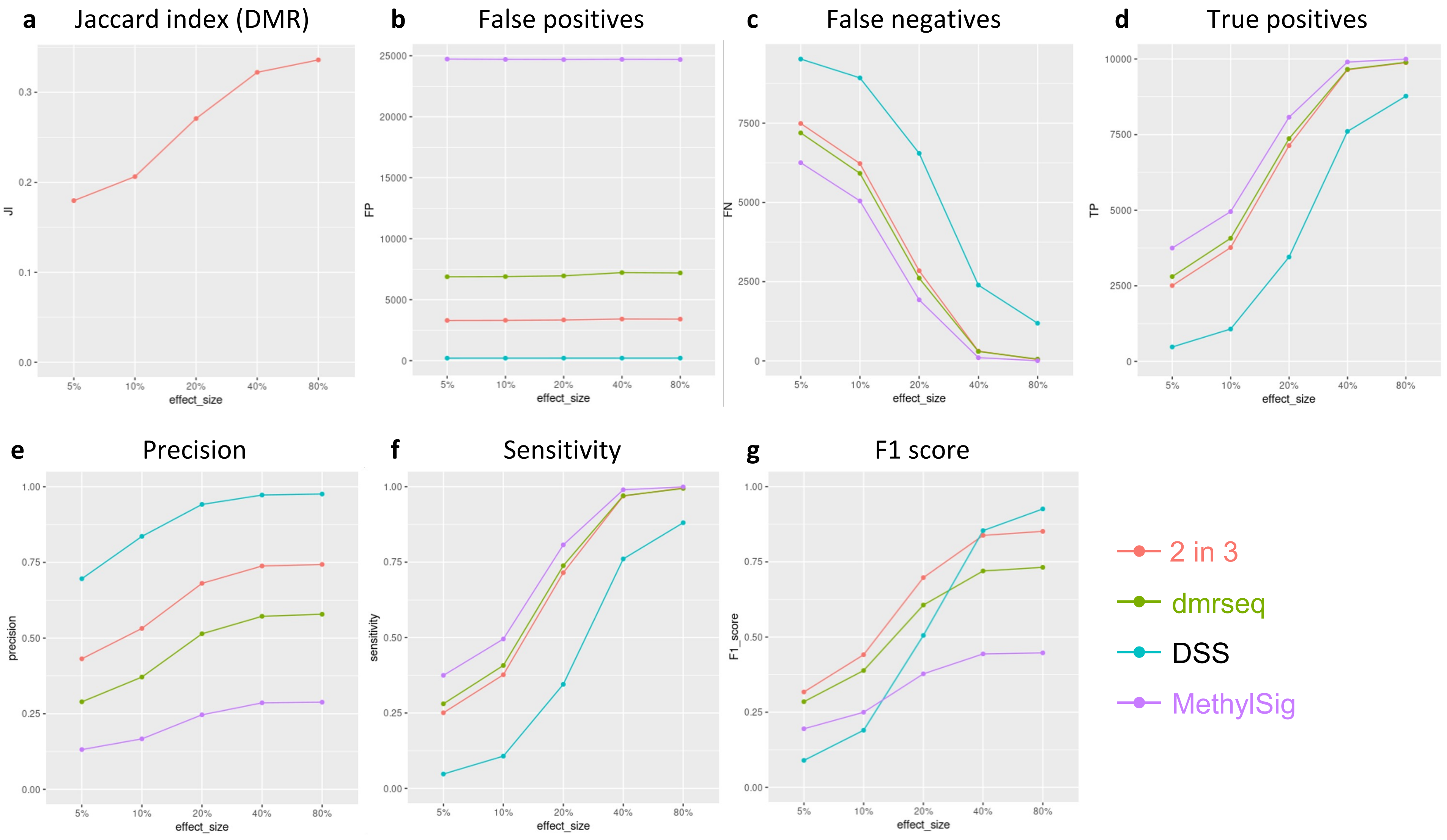

### Supplementary Figure 4

**Supplementary Figure 4**

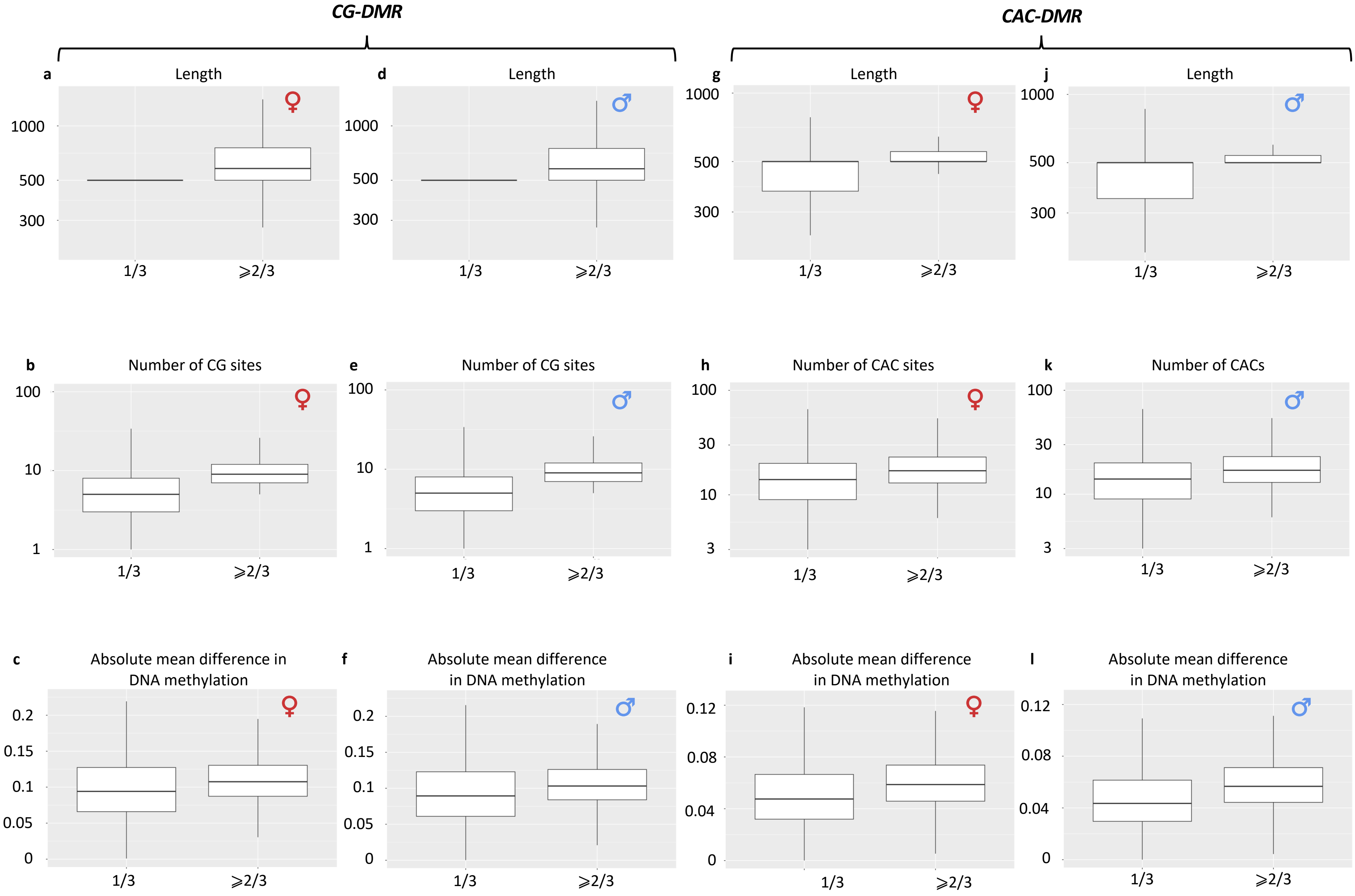

### Supplementary Figure 6

**Supplementary Figure 6**

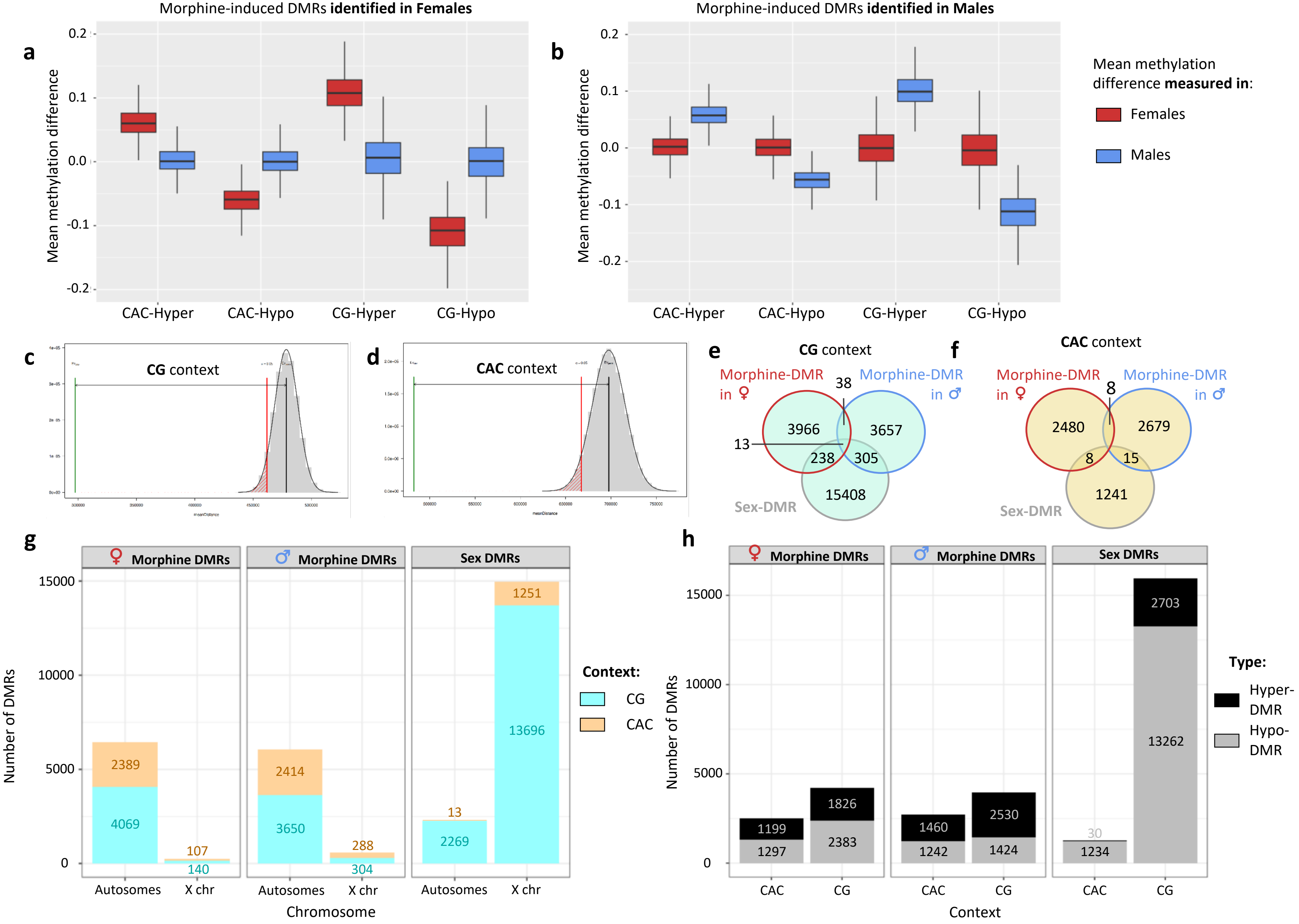

### Supplementary Figure 8

#### Supplementary Figure 8

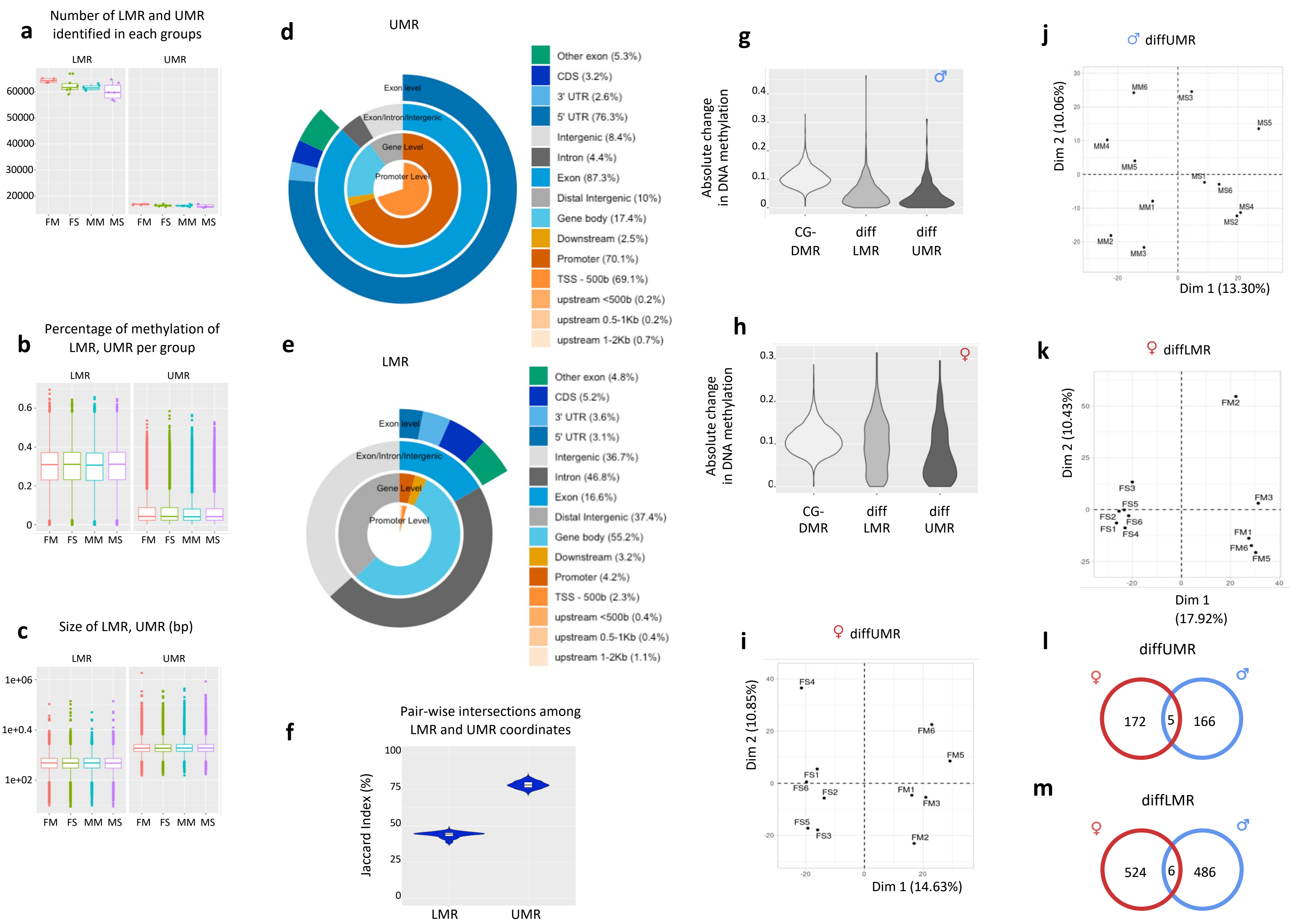

### Supplementary Figure 9

#### Supplementary Figure 9

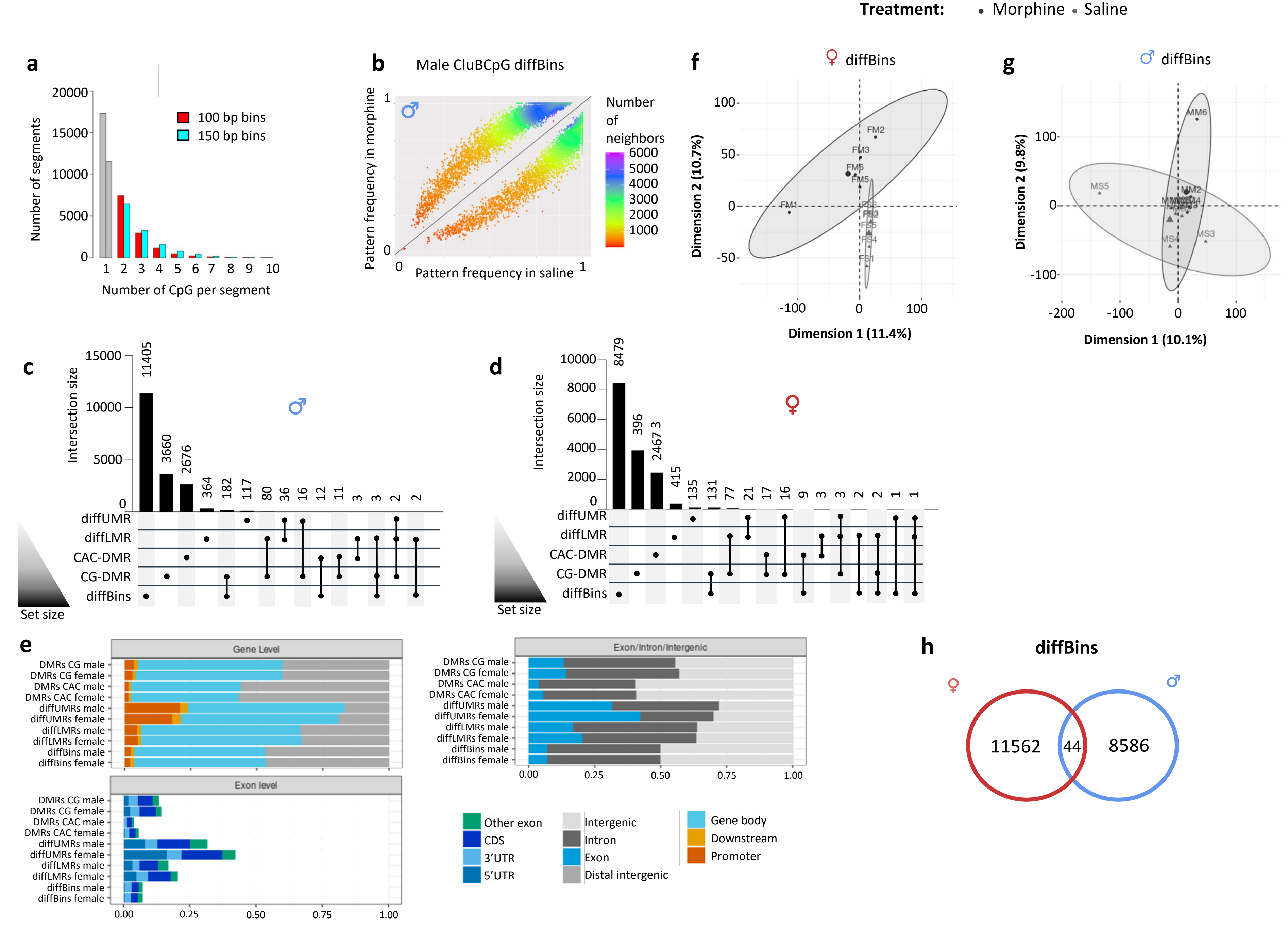

### Supplementary Figure 11

#### Supplementary Figure 11

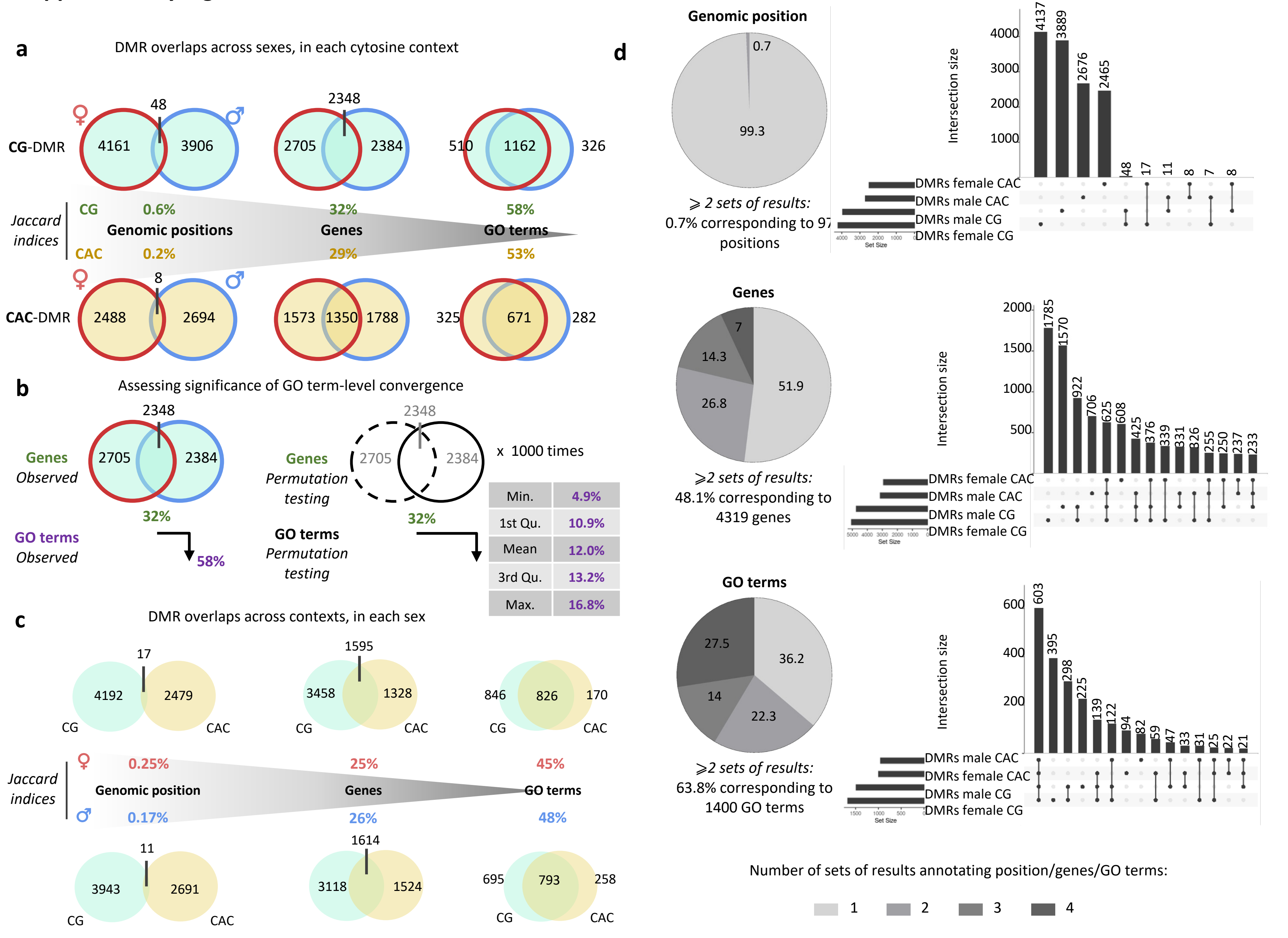

### Supplementary Figure 12

**Supplementary Figure 12**

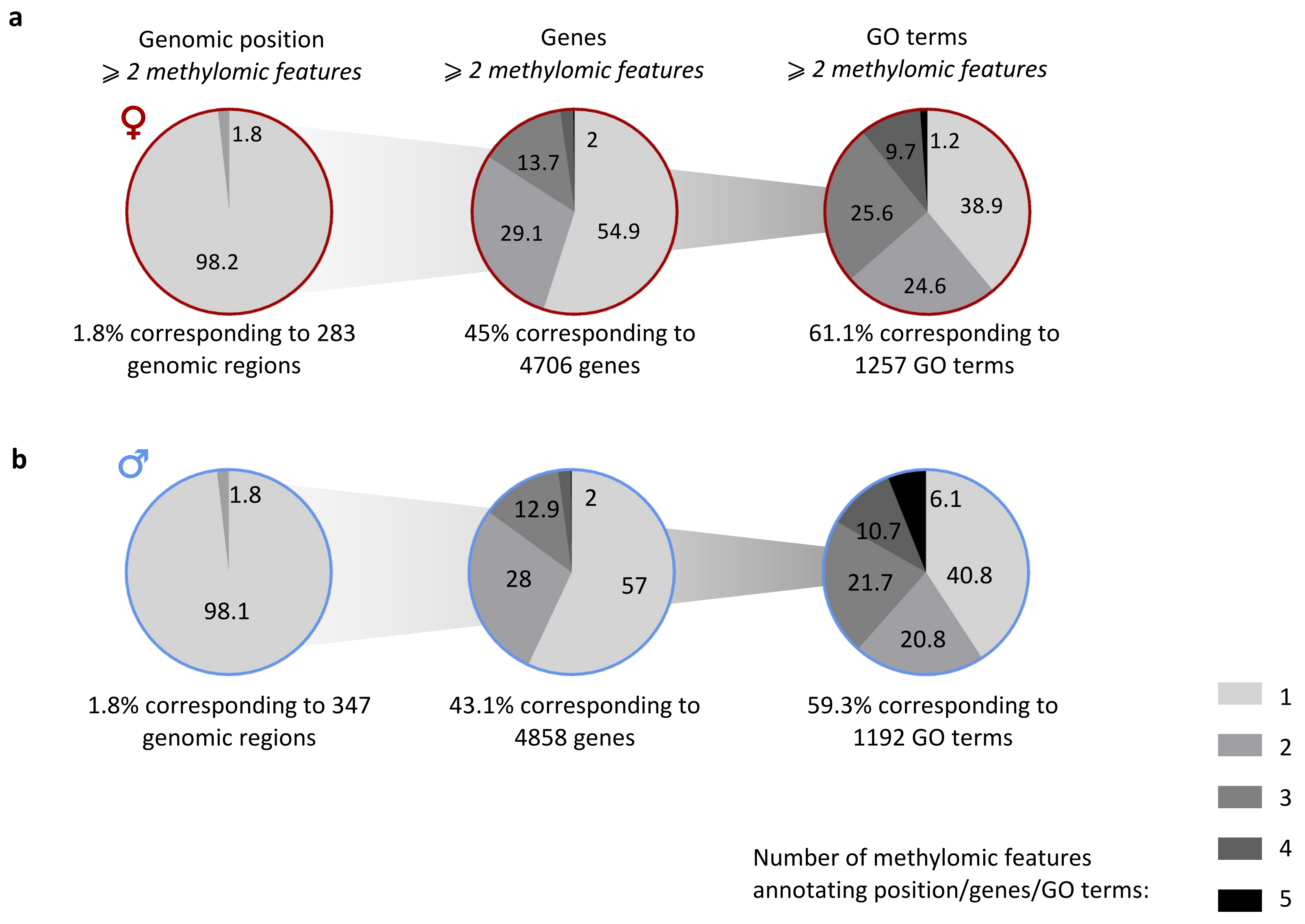

### Supplementary Figure 15

#### Supplementary Figure 15

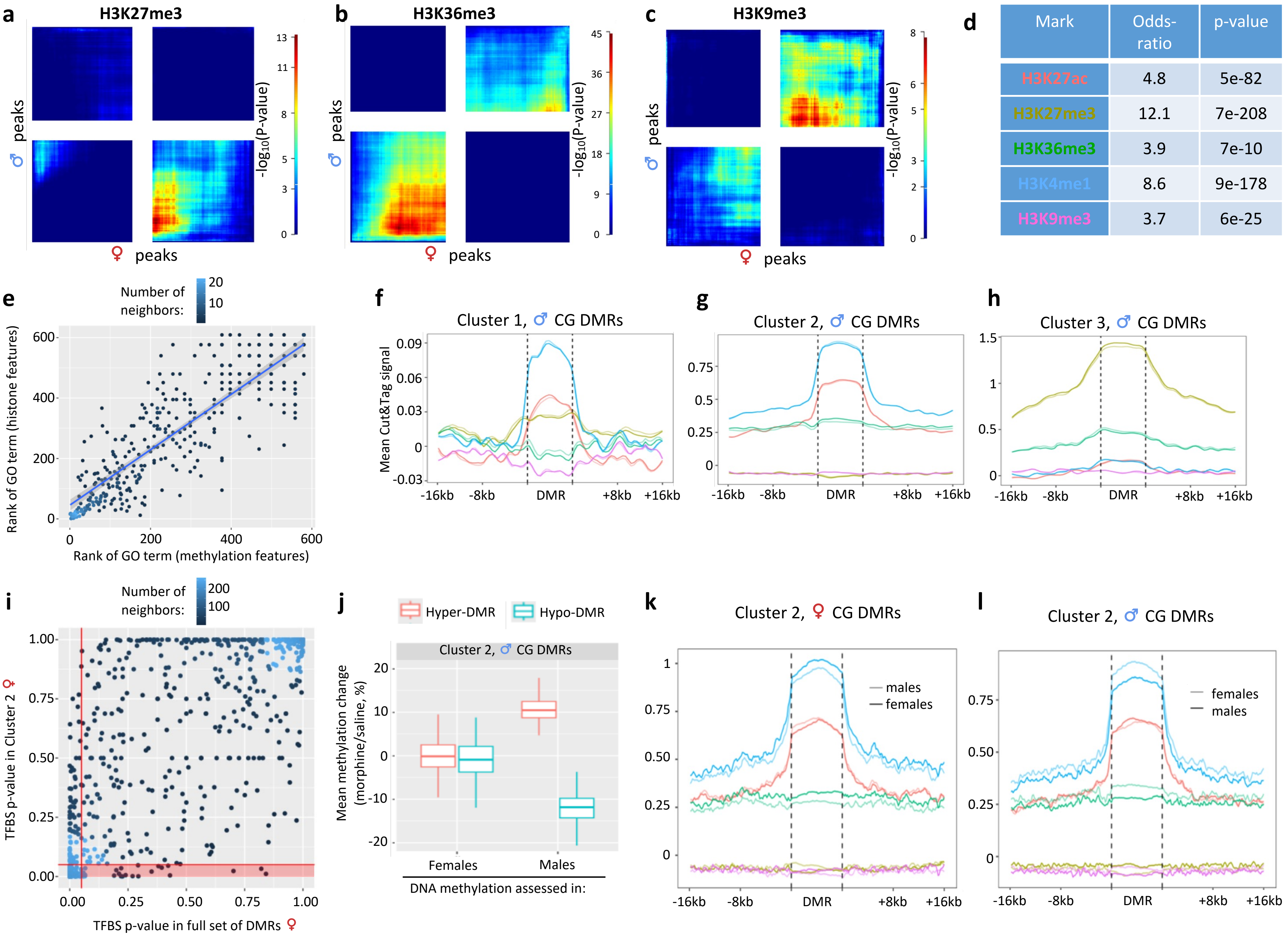


| Mark | Odds-ratio | p-value |
|---|---|---|
| H3K27ac | 4.8 | 5e-82 |
| H3K27me3 | 12.1 | 7e-208 |
| H3K36me3 | 3.9 | 7e-10 |
| H3K4me1 | 8.6 | 9e-178 |
| H3K9me3 | 3.7 | 6e-25 |

### Supplementary Figure 16

##### Supplementary Figure 16

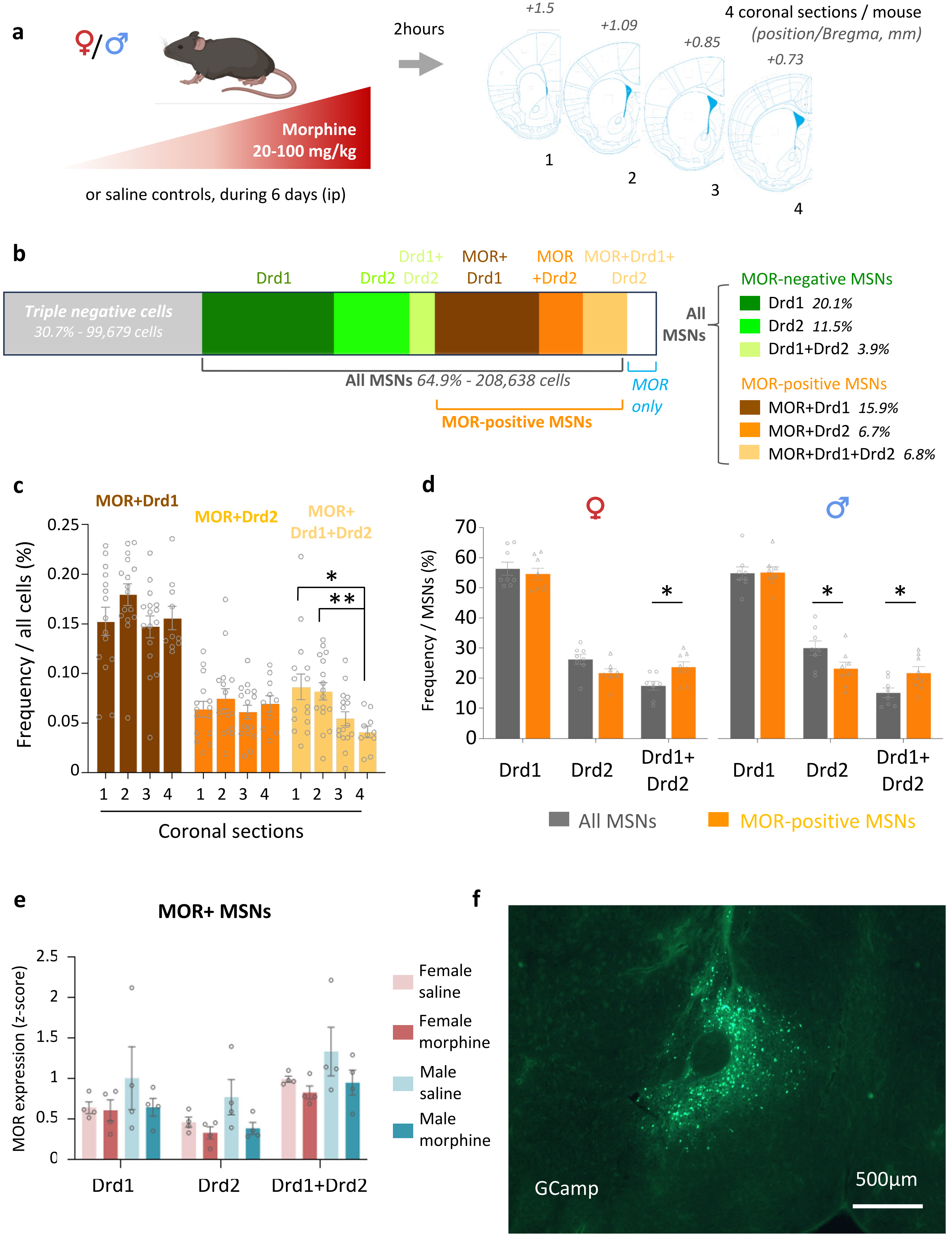

### Supplementary Figure 17

#### Supplementary Figure 17

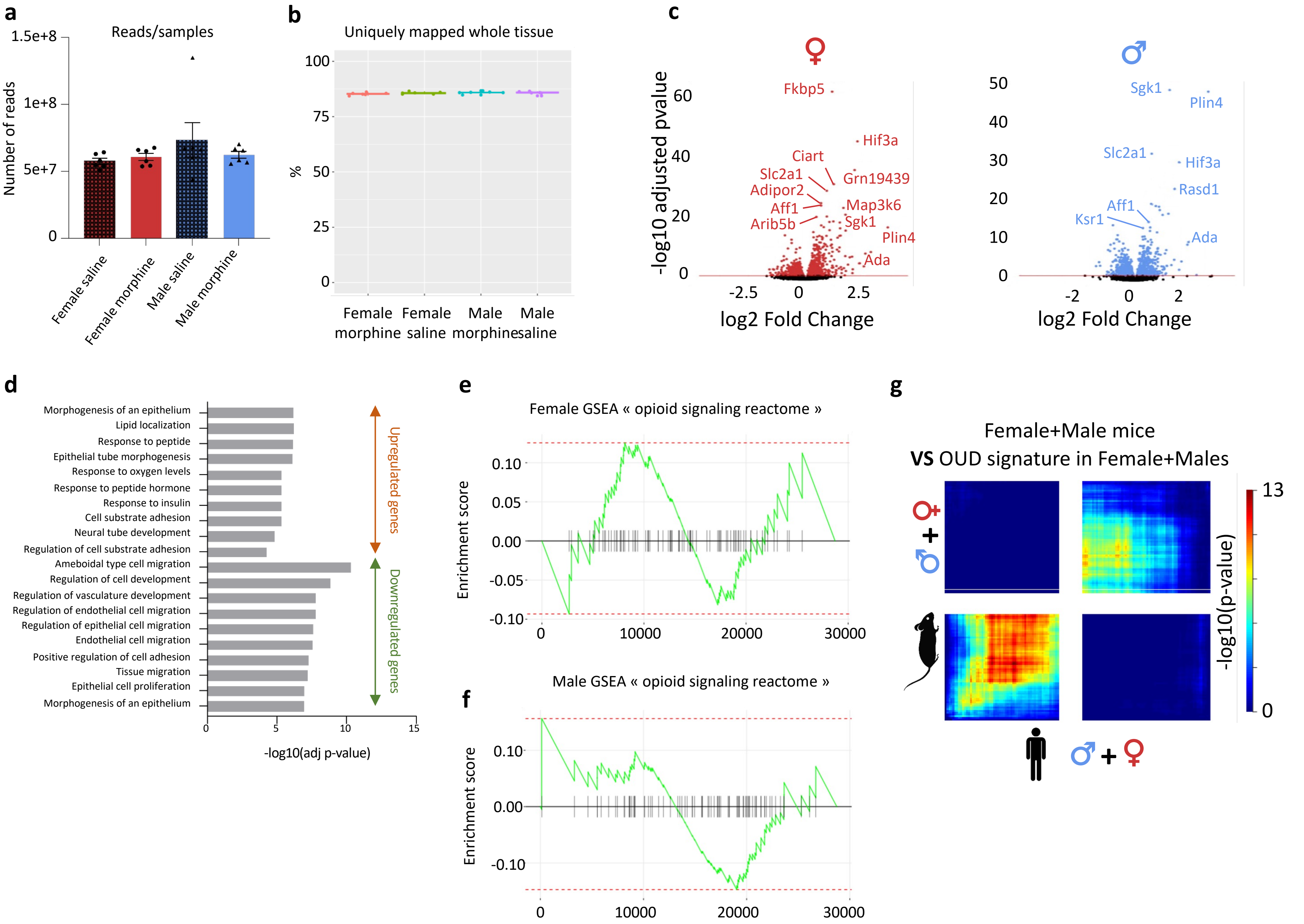

### Supplementary Figure 18

#### Supplementary Figure 18

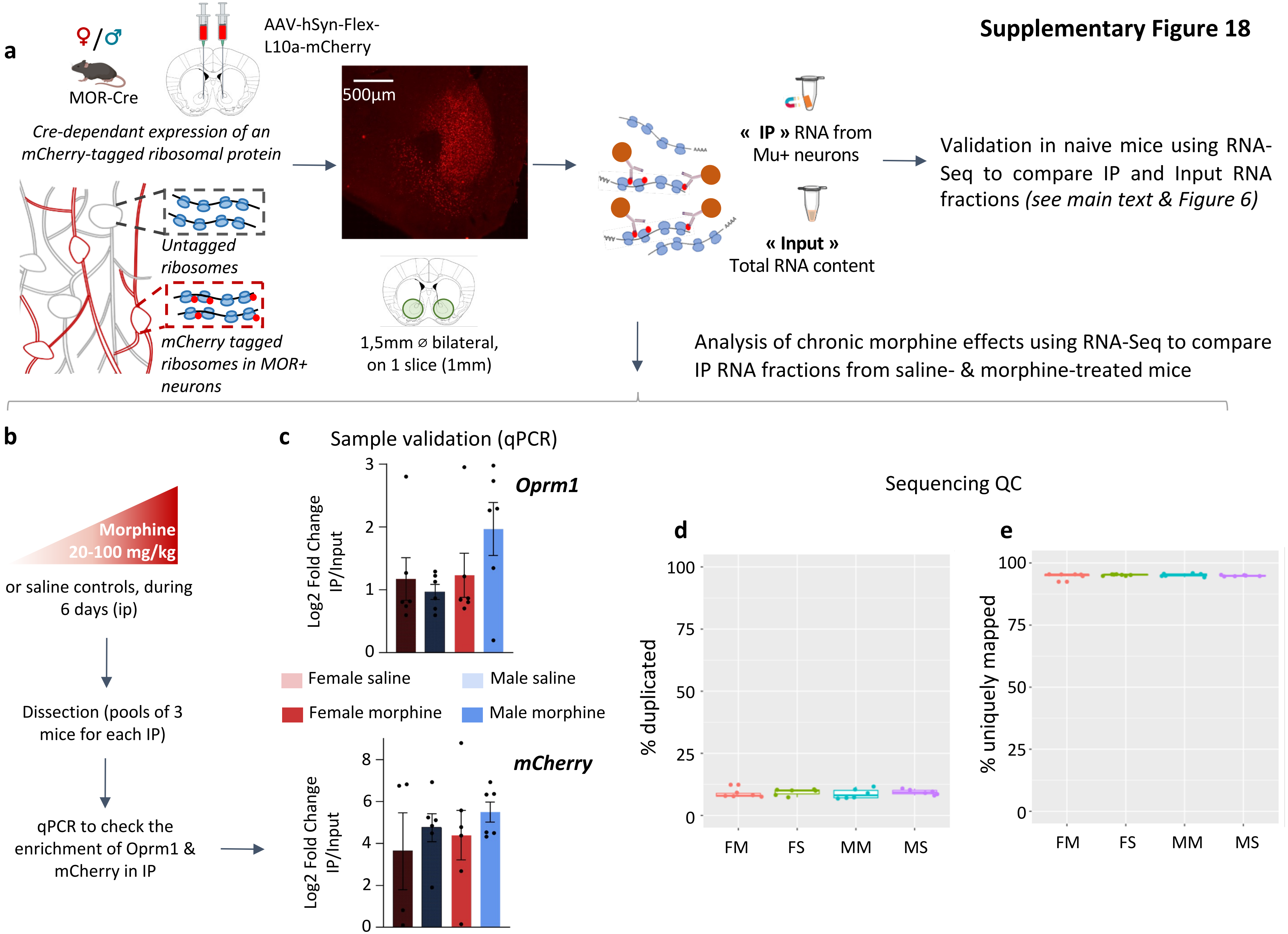

### Supplementary Figure 19

##### Supplementary Figure 19

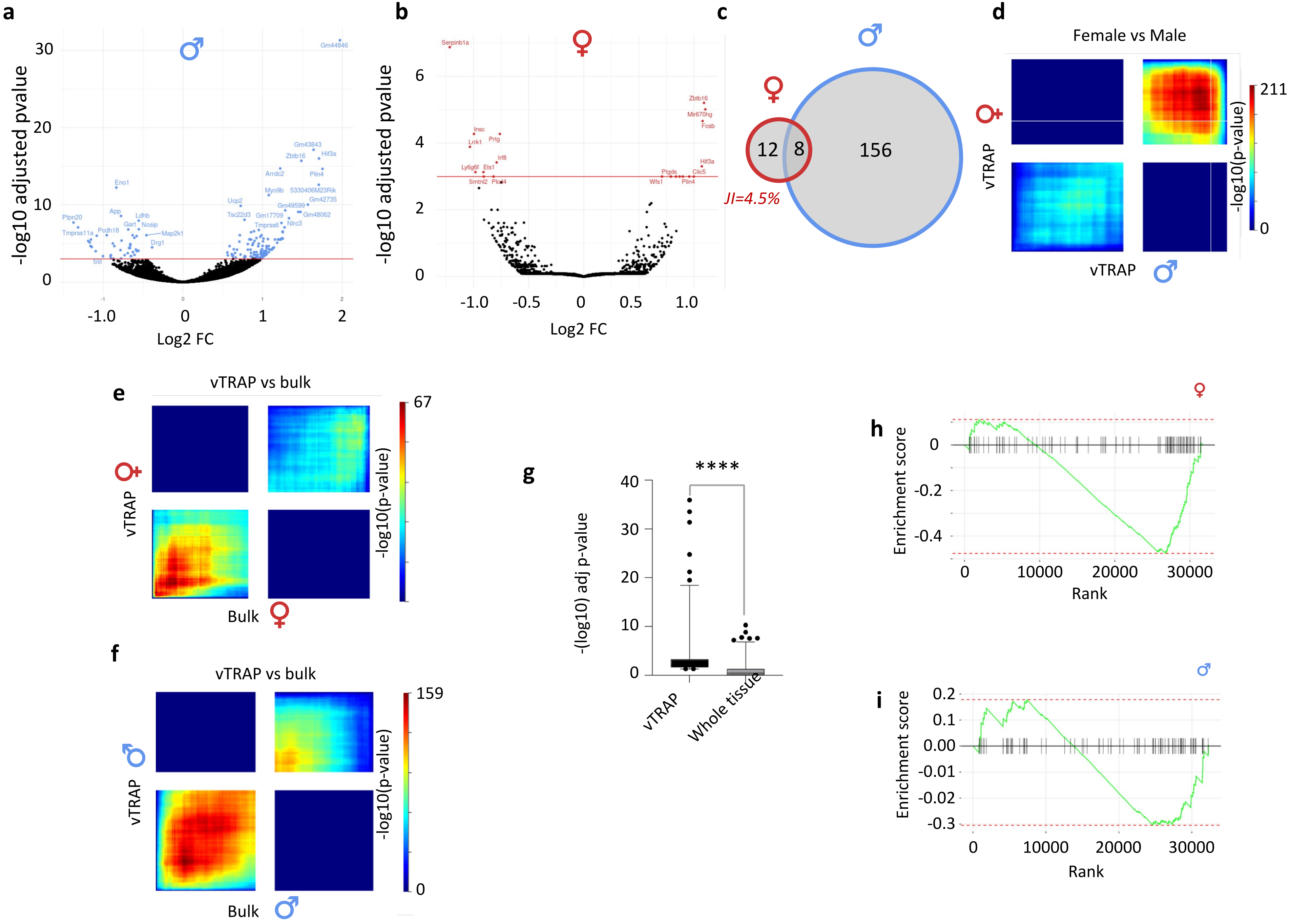

### Supplementary Figure 20

##### Supplementary Figure 20

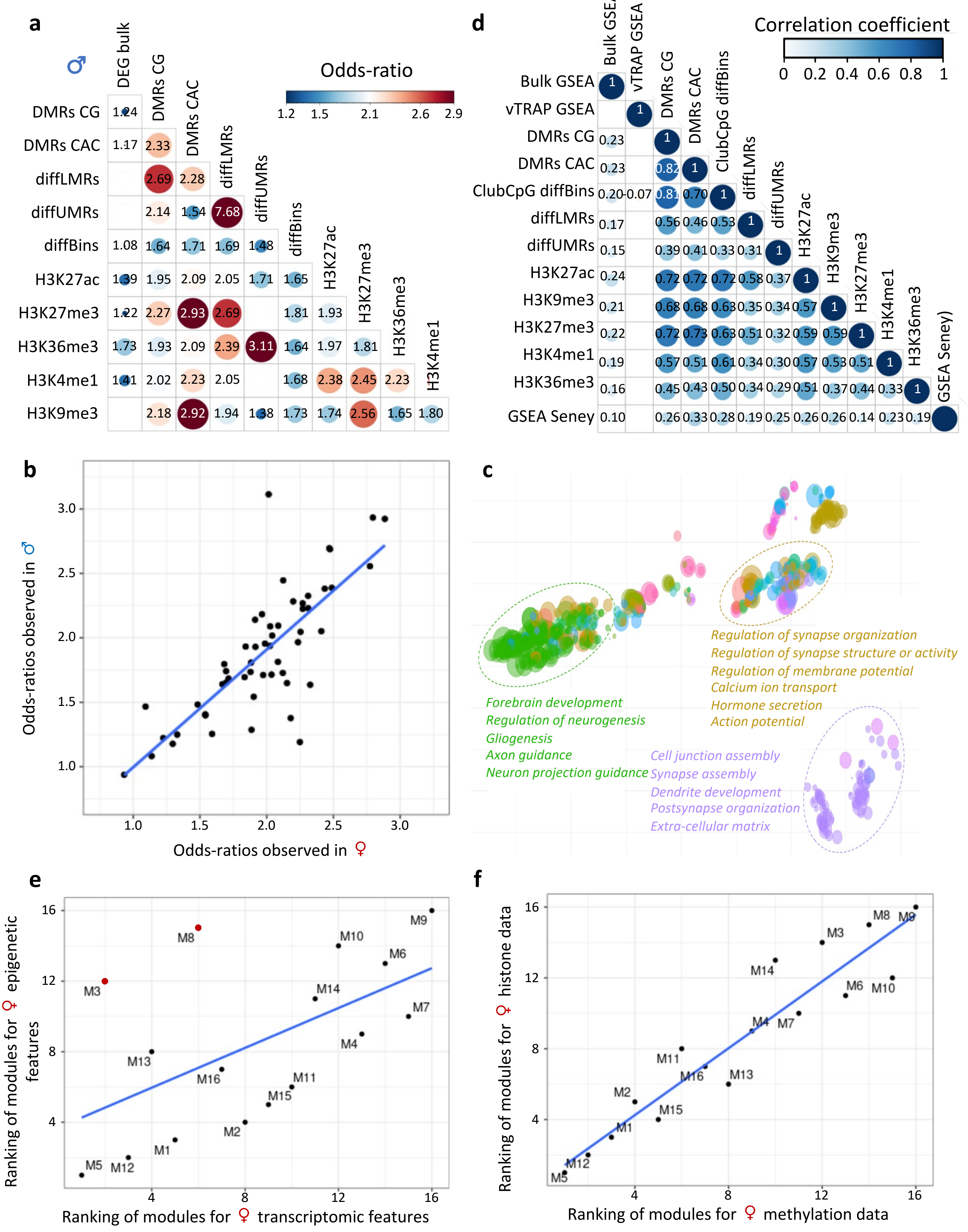
