## Supplementary Figure 5 for "Sex-divergent brain epigenetic reprogramming by chronic opioids"

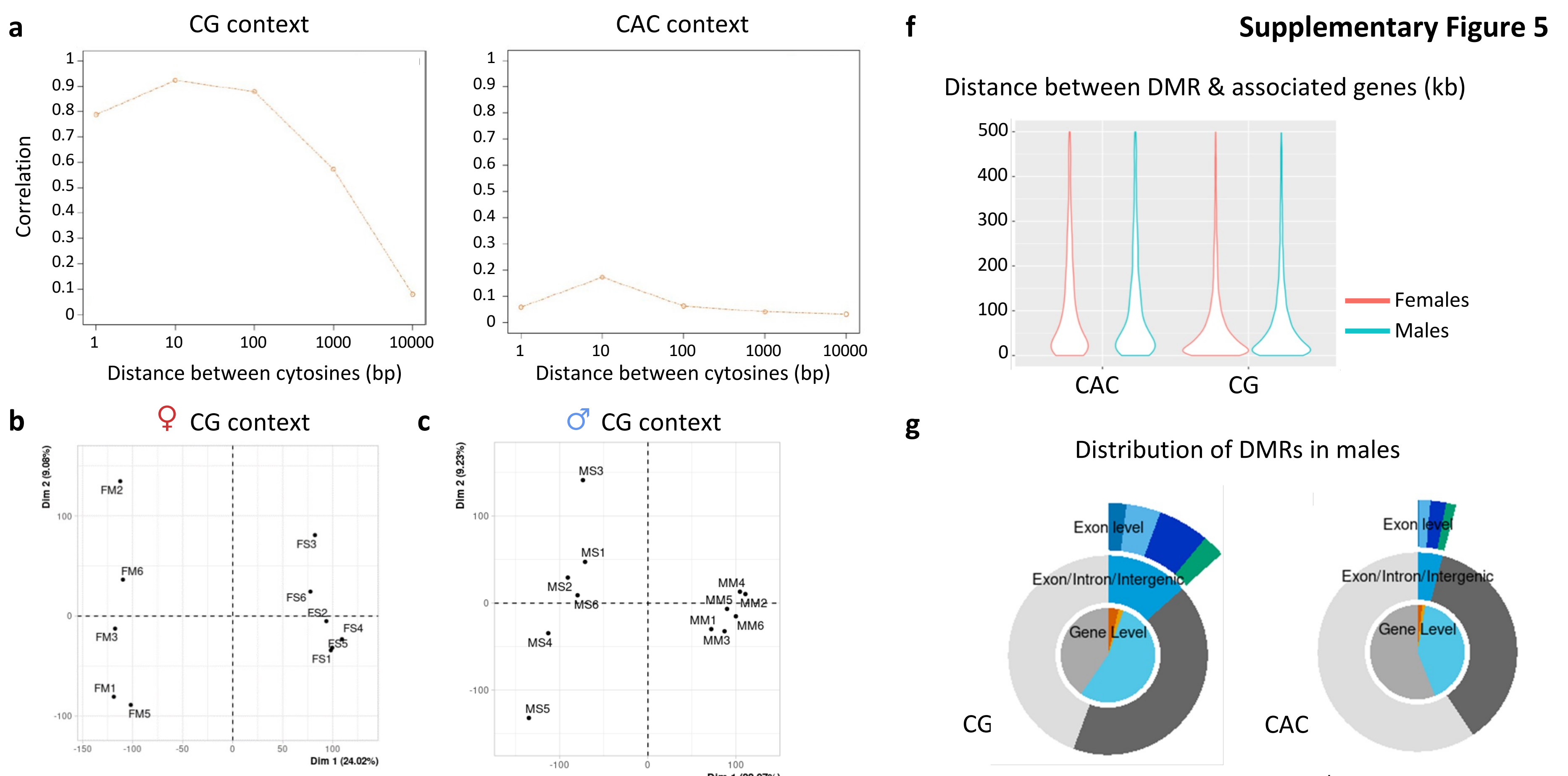


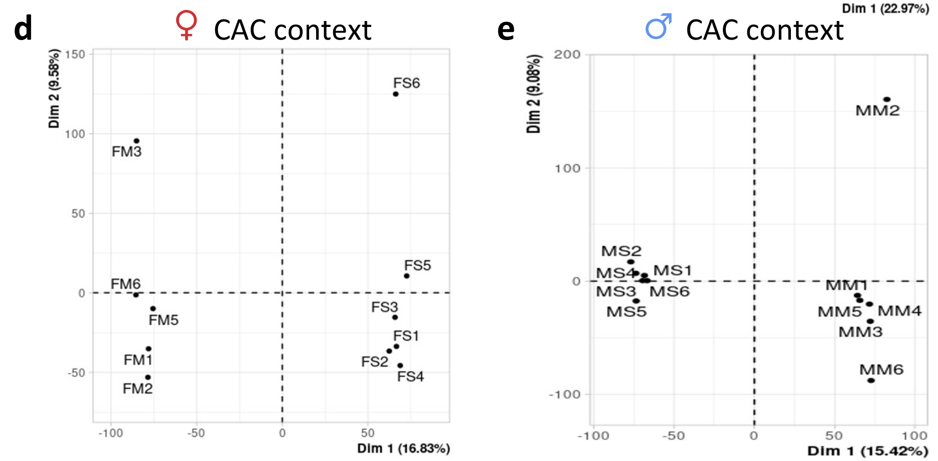


| | | CG-DMR (%) | CAC-DMR (%) |
|---|---|---|---|
| Outer circle | Other exon | 2.4 | 1.1 |
| | CDS | 5.4 | 1.7 |
| | 3'UTR | 3.6 | 1.2 |
| | 5'UTR | 1.9 | 0.2 |
| Middle circle | Intergenic | 44.4 | 59.4 |
| | Intron | 42.2 | 36.5 |
| | Exon | 13.4 | 4.1 |
| Inner circle | Distal intergenic | 40.4 | 56.1 |
| | Gene body | 54.4 | 41.3 |
| | Downstream | 1.6 | 1 |
| | Promoter | 3.6 | 1.6 |
