## Supplementary Figure 7 for "Sex-divergent brain epigenetic reprogramming by chronic opioids"

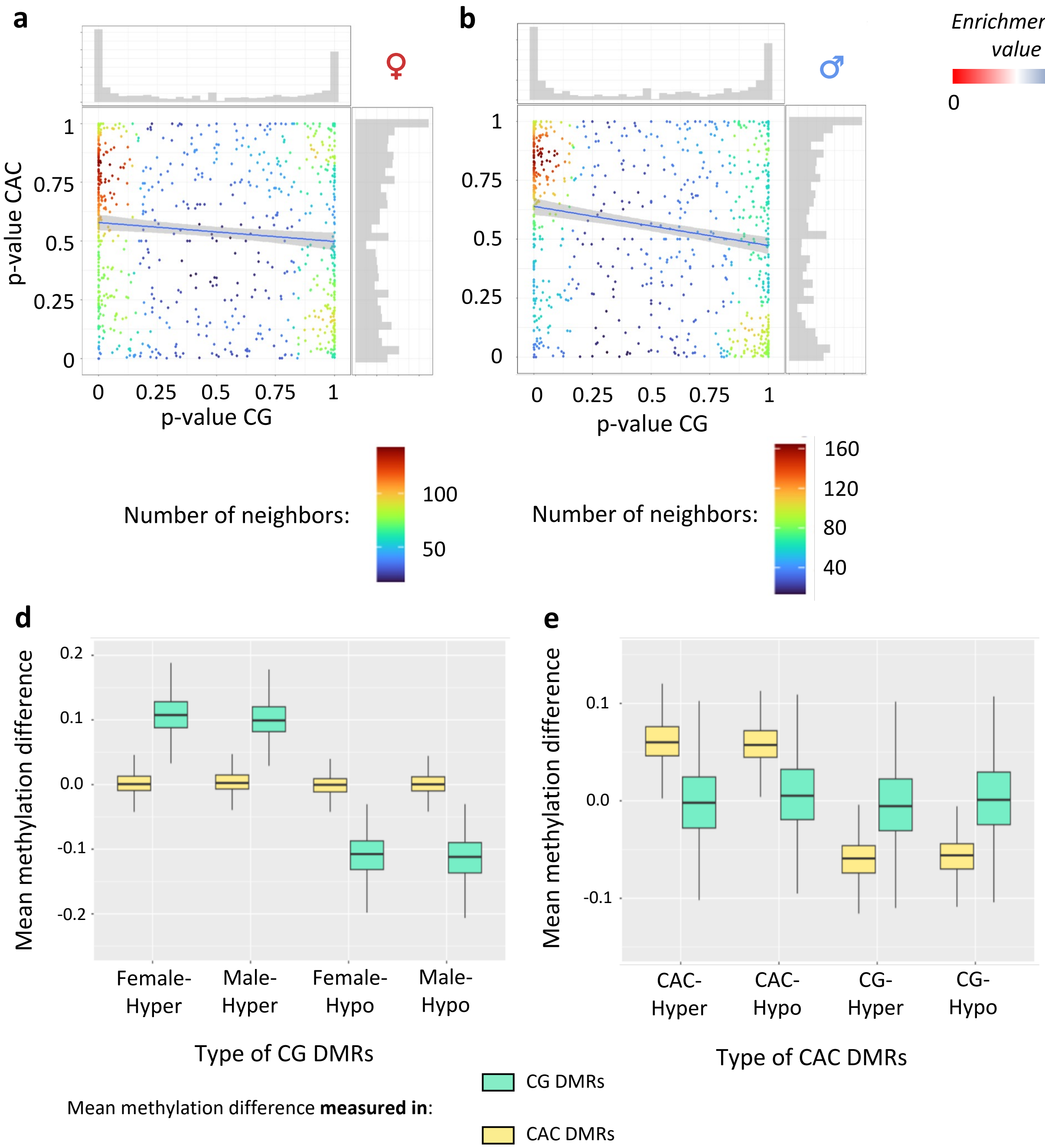


**c**

TFs among the 10 best in both sexes (CG-DMR)
TFs among the 10 best in both sexes (CAC-DMR)

*Enrichment p-value*: 0 – 1

| | | CG Fem. | CG Mal. | CAC Fem. | CAC Mal. |
|---|---|---|---|---|---|
| Female CG DMRs | ARNT_HIF1A | 1.3E-32 | 4.1E-27 | 8.7E-01 | 7.8E-01 |
| | AHR_ARNT | 3.0E-23 | 1.3E-17 | 2.3E-01 | 8.2E-01 |
| | GMEB1 | 4.7E-23 | 3.8E-14 | 3.4E-01 | 7.0E-01 |
| | HIF1A | 7.2E-17 | 1.5E-12 | 6.8E-01 | 8.3E-01 |
| | ATF1 | 3.2E-16 | 3.4E-12 | 5.4E-01 | 4.0E-01 |
| | TFDP1 | 1.4E-15 | 3.2E-17 | 6.1E-01 | 9.7E-01 |
| | E2F6 | 1.5E-14 | 7.6E-12 | 5.2E-01 | 7.1E-01 |
| | ZIC1_ZIC2 | 2.5E-14 | 4.3E-13 | 6.7E-01 | 7.9E-01 |
| | ZBTB14 | 1.0E-13 | 2.1E-14 | 2.5E-01 | 7.0E-01 |
| | ARNT | 1.8E-13 | 6.1E-14 | 8.0E-01 | 9.7E-01 |
| Male CG DMRs | ARNT_HIF1A | 1.3E-32 | 4.1E-27 | 8.7E-01 | 7.8E-01 |
| | AHR_ARNT | 3.0E-23 | 1.3E-17 | 2.3E-01 | 8.2E-01 |
| | TFDP1 | 1.4E-15 | 3.2E-17 | 6.1E-01 | 9.7E-01 |
| | ZNF263 | 1.7E-11 | 1.5E-14 | 2.9E-01 | 2.4E-01 |
| | ZBTB14 | 1.0E-13 | 2.1E-14 | 2.5E-01 | 7.0E-01 |
| | GMEB1 | 4.7E-23 | 3.8E-14 | 3.4E-01 | 7.0E-01 |
| | ARNT | 1.8E-13 | 6.1E-14 | 8.0E-01 | 9.7E-01 |
| | FOXN1 | 2.2E-11 | 3.2E-13 | 6.4E-01 | 4.7E-01 |
| | ZIC1_ZIC2 | 2.5E-14 | 4.3E-13 | 6.7E-01 | 7.9E-01 |
| | ZIC3 | 1.3E-12 | 9.6E-13 | 5.7E-01 | 5.8E-01 |
| Female CAC DMRs | KLF9 | 1.7E-03 | 9.3E-03 | 5.2E-07 | 8.8E-06 |
| | EGR1 | 2.3E-11 | 2.5E-08 | 3.4E-04 | 3.6E-05 |
| | PAX4 | 8.4E-01 | 8.1E-01 | 6.0E-04 | 7.7E-06 |
| | DLX2 | 2.9E-01 | 1.3E-01 | 7.2E-04 | 3.0E-03 |
| | DLX5 | 7.8E-01 | 3.1E-01 | 1.9E-03 | 2.1E-02 |
| | DRGX | 7.7E-01 | 9.3E-01 | 2.0E-03 | 1.5E-03 |
| | EOMES | 4.3E-01 | 5.4E-01 | 3.3E-03 | 3.6E-01 |
| | TBX3 | 2.5E-01 | 2.9E-01 | 3.5E-03 | 3.8E-01 |
| | LIN54 | 1.0E+00 | 1.0E+00 | 5.2E-03 | 1.5E-02 |
| | MGA | 1.2E-01 | 1.0E-01 | 5.4E-03 | 3.9E-02 |
| Male CAC DMRs | PAX4 | 8.4E-01 | 8.1E-01 | 6.0E-04 | 7.7E-06 |
| | KLF9 | 1.7E-03 | 9.3E-03 | 5.2E-07 | 8.8E-06 |
| | EGR1 | 2.3E-11 | 2.5E-08 | 3.4E-04 | 3.6E-05 |
| | BARX2 | 1.0E+00 | 1.0E+00 | 7.0E-02 | 5.2E-04 |
| | ALX4 | 1.3E-01 | 8.0E-01 | 4.0E-02 | 5.8E-04 |
| | LHX1 | 1.3E-01 | 3.7E-01 | 4.2E-02 | 9.7E-04 |
| | HOXB9 | 4.8E-01 | 2.0E-01 | 5.3E-01 | 9.7E-04 |
| | ARID3B | 1.0E+00 | 1.0E+00 | 9.5E-02 | 1.1E-03 |
| | ALX1 | 2.5E-01 | 6.5E-01 | 2.3E-02 | 1.2E-03 |
| | DRGX | 7.7E-01 | 9.3E-01 | 2.0E-03 | 1.5E-03 |
