## Supplementary Figure 10 for "Sex-divergent brain epigenetic reprogramming by chronic opioids"

**Threshold-free comparison of morphine effects in male & females (RedRibbon)**

**a** CG context, up/up quadrant

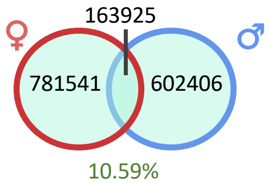


**b** CG context, down/down quadrant

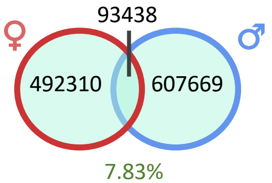


**c** CAC context, up/up quadrant

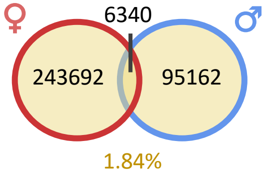


**d** CAC context, down/down quadrant

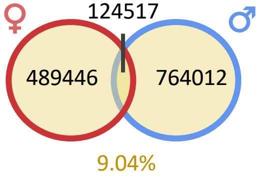


**e**

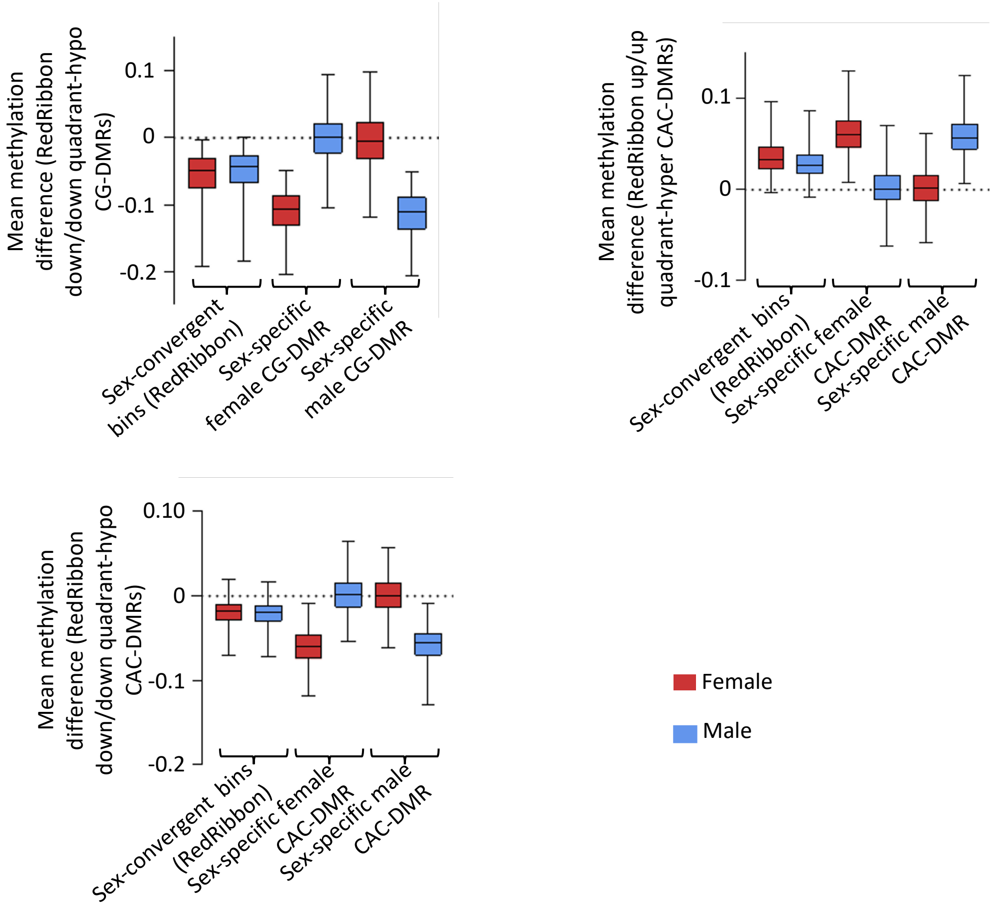


Permutation testing of distances between consensus DMR and sex-convergent 500-bp genomic bins (RedRibbon)

**f**

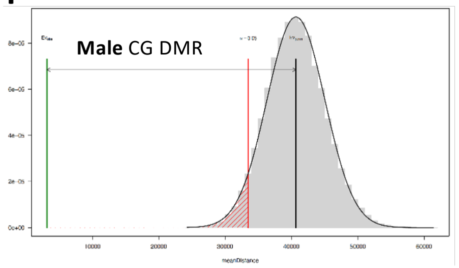


**g**

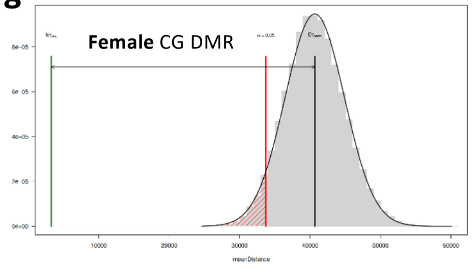
