## Supplementary Figure 13 for "Sex-divergent brain epigenetic reprogramming by chronic opioids"

**a**

| Mark | Million reads/ lib (mean+/-sem) | Number of peaks in M | Number of peaks in F |
|---|---|---|---|
| 27ac | 16.9 +/-0.299 | 109059 | 110273 |
| 27me3 | 15.0 +/-0.376 | 35275 | 42395 |
| 36me3 | 15.1 +/-0.449 | 27616 | 28630 |
| 4me1 | 17.9 +/-0.675 | 60070 | 62760 |
| 9me3 | 18.6 +/-0.582 | 74501 | 83215 |

**d**

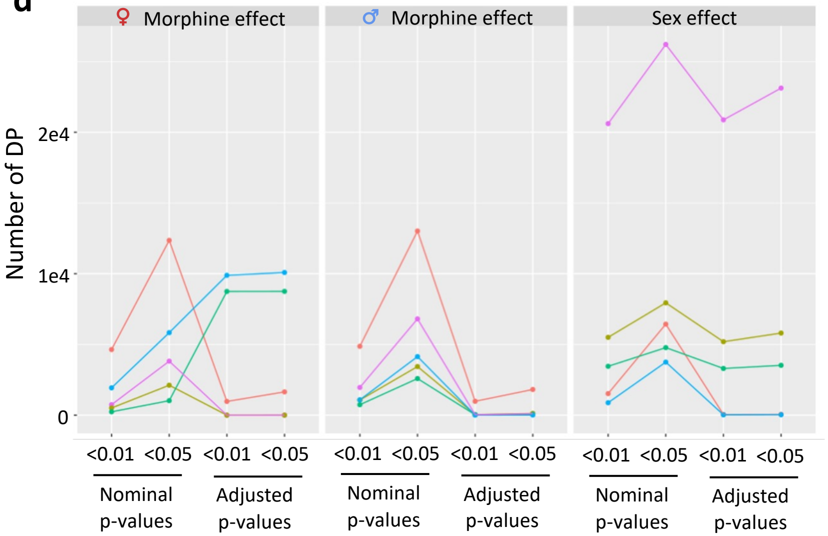


**f**

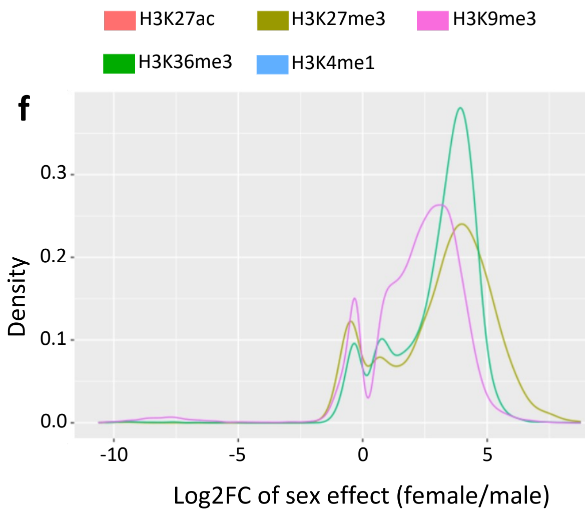


**b** Normalized by sex-specific IgG

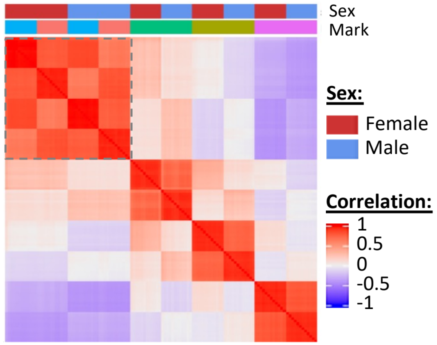


**e**

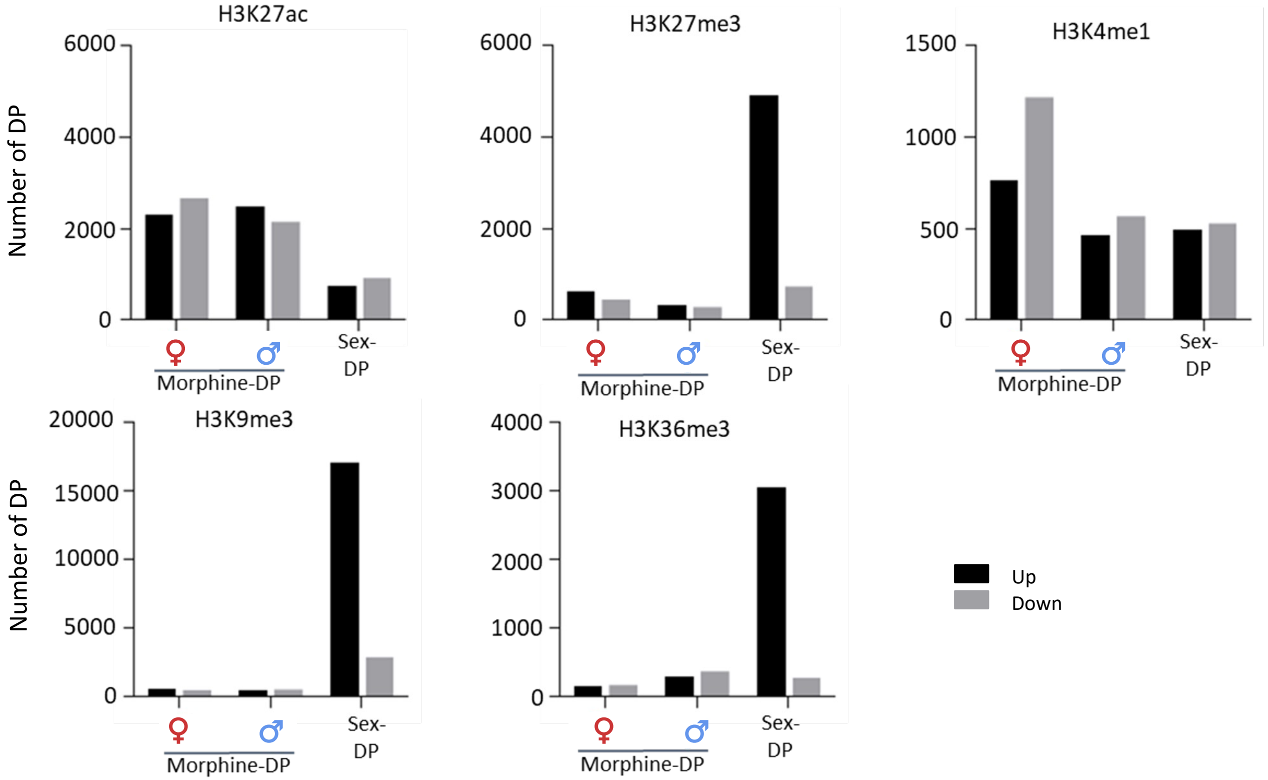


**c** Normalized by batch-specific IgG

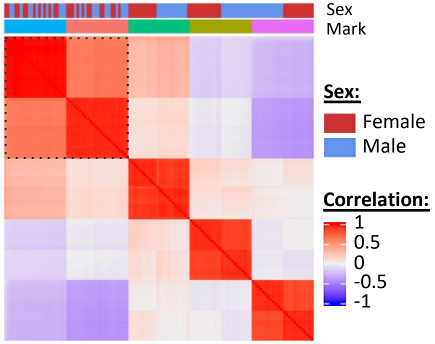
