## Supplementary Figure 14 for "Sex-divergent brain epigenetic reprogramming by chronic opioids"

**a**

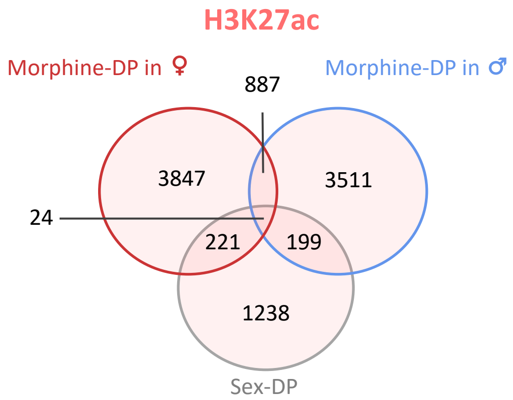


**b**

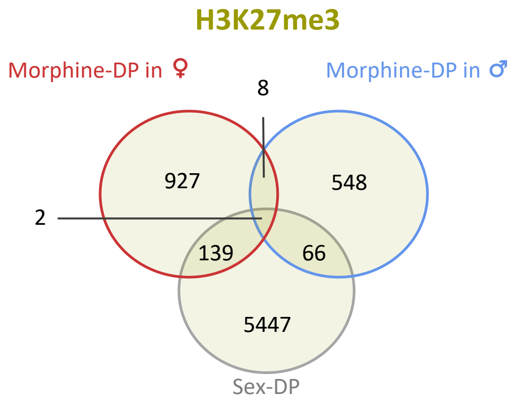


**c**

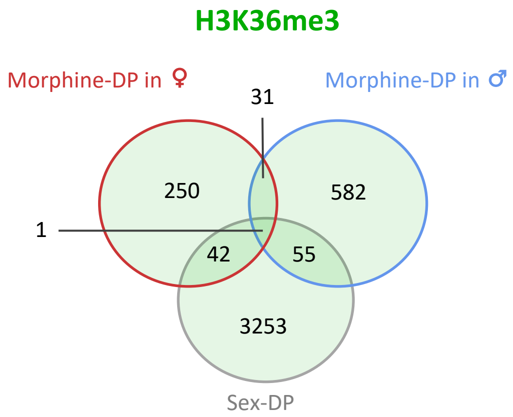


**d**

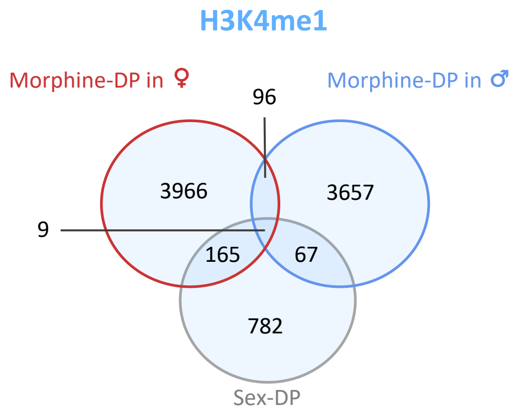


**e**

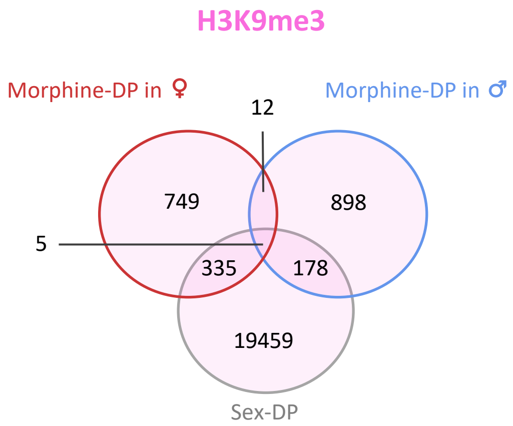


**e**

| Mark | Jaccard Index in females (%) | Jaccard Index in males (%) |
|---|---|---|
| K27ac | 3.8 | 3.7 |
| K36me3 | 1.1 | 1.4 |
| K9me3 | 1.6 | 0.9 |
| K27me3 | 2.1 | 1.1 |
| K4me1 | 6.1 | 3.8 |
